# AI-Driven Computational Design of Peptide-Based WWP1 Inhibitors as Promising Therapeutic Agents Against Breast Cancer, Including Triple-Negative Subtype

**DOI:** 10.64898/2026.08.08.742959

**Authors:** Enrico Mario Alessandro Fassi, Sara Mathlouthi, Elena Maspero, Edoardo Sisti, Giulia Tamboia, Giuseppe De Vita, Fabio Forlani, Simona Polo, Alessandro Gori, Kaliroi Peqini, Sara Pellegrino, Gabriella Roda, Jacopo Sgrignani, Andrea Cavalli, Luisa De Cola, Mariangela Garofalo, Giovanni Grazioso

## Abstract

Breast cancer (BC) is the second most common noncutaneous cancer and the second leading cause of cancer-related death in women. BC is classified into three primary subtypes, with triple-negative breast cancer (TNBC) having the poorest prognosis because it lacks specific targetable markers. Preclinical studies on TNBC indicated a common occurrence of diminished tumor-suppressor activity of PTEN, activating the PI3K/AKT/mTOR signaling pathway. Notably, published studies reveal that the WWP1 enzyme plays a pivotal role in driving PTEN degradation via ubiquitination, unveiling a promising therapeutic target for treating TNBC. In the search of new WWP1 inhibitors, we used artificial intelligence (AI)-driven computational strategies for de novo design of peptide-based WWP1 inhibitors and identified a hexapeptide, termed **WI23-B**, which demonstrated high nanomolar binding affinity to WWP1. In TR-FRET enzymatic assays, **WI23-B** inhibited WWP1 activity with an IC₅₀ of approximately 11 µM. In MCF7 and MDA-MB-231 breast cancer cell lines, **WI23-B** showed promising cytotoxic efficacy, particularly in combination with the PI3K inhibitor BYL719, also when it was loaded into nanocapsules. Collectively, these findings highlight **WI23-B** as a promising lead peptide with potent WWP1 inhibitory activity and synergistic antiproliferative effects when combined with PI3K inhibitors. While further structural optimization is required to enhance its potency and pharmacological properties, our results provide a strong foundation for the development of next-generation WWP1 inhibitors. Such agents have the potential to reshape therapeutic strategies for BC and TNBC by enabling more effective and less toxic treatment regimens, ultimately reducing the reliance on high-dose chemotherapy and minimizing adverse effects.

## 1. INTRODUCTION

Breast cancer (BC) represents the most frequently identified cancer in women and the second most prevalent cause of cancer-related death globally^1^. Over the years, considerable advancements have been achieved in treating BC displaying hormone receptors like estrogen receptor (ER) and progesterone receptor (PR), along with effectively targeting human epidermal growth factor receptor 2 (HER2). Nevertheless, triple-negative breast cancer (TNBC), lacking the expression of these receptors, carries a poorer prognosis compared to hormone receptor-positive BCs^2^. TNBC constitutes approximately 15% to 20% of all BC cases^3^. Within the initial 3 to 5 years post-diagnosis, over half of TNBC patients encounter relapse, and the median overall survival (OS) using current treatments is about 10.2 months^4,5^. Since TNBC lacks the receptors targeted by established endocrine or HER2-directed drugs, standard therapy typically involves general chemotherapy utilizing taxanes or anthracyclines. In 2011, Lehmann introduced six categories of TNBC subtypes, including basal-like (BL1 and BL2), immunomodulatory (IM), luminal androgen receptor (LAR), mesenchymal (M), and mesenchymal-stem-like (MSL) subtypes^6^. The latter two subtypes exhibit five primary pathways or functional changes: alterations in cell cycle regulation, amplification of growth factor receptors, modifications in the RAS/mitogen-activated protein kinase (MAPK) pathway, changes in DNA repair mechanisms, and alterations in the PI3K/AKT/mTOR and/or phosphatase and tensin homolog (PTEN) pathways. Notably, PTEN plays a crucial role in activating or deactivating PI3K^6,7^. Specifically, it has been shown that PTEN ubiquitination by the proto-oncogenic WW domain-containing E3 ubiquitin protein ligase 1 (WWP1) suppresses PTEN localization at the plasma membrane, consequently impairing its ability to counteract the proliferative, growth, and resistance to chemo- and immuno-therapy in cancer cells signals triggered by the PI3K/AKT/mTOR pathway activation^8,9^.

The PTEN gene encodes a lipid phosphatase enzyme that is pivotal in regulating intracellular signaling pathways controlling cell growth, survival, and proliferation, primarily by dephosphorylating the phospholipid PIP₃ and antagonizing the PI3K/AKT signaling axis, a pathway frequently hyperactivated in various cancers, including BC^10^. Under normal physiological conditions, PTEN acts as a potent tumor suppressor, maintaining cellular homeostasis by balancing proliferation and apoptosis^11^. However, loss or mutation of PTEN is a relatively common event in BC, particularly in more aggressive subtypes such as TNBC and HER2-positive tumors, and is often associated with poor prognosis and therapeutic resistance. Its inactivation contributes to uncontrolled tumor cell growth and may serve as both a prognostic marker and a potential therapeutic target^10,12–14^. Understanding the role of PTEN in breast tumorigenesis is therefore critical for developing more precise diagnostic tools and effective treatment strategies.

WWP1 is a member of the NEDD4 family of HECT-type E3 ubiquitin ligases, comprising four tandem WW domains and a catalytic HECT domain, and plays a critical role in post-translational regulation by ubiquitinating substrate proteins^15^. In recent years, WWP1 has garnered attention for its involvement in cancer progression where it has been found to be frequently overexpressed^16^. In BC, WWP1 is frequently amplified and overexpressed, particularly in HER2-positive and ER-positive tumors, and its elevated expression has been significantly associated with gene copy number gains, PTEN inactivation, and poor clinical outcomes, including increased proliferation, metastasis, and therapy resistance^17,18^. Functionally, WWP1 promotes breast cancer cell proliferation, anchorage-independent growth, and resistance to apoptosis, while its depletion induces caspase-mediated cell death in breast cancer cell lines such as MCF7 and HCC1500^19^. WWP1 was recently identified as an E3 ligase for PTEN, with the strongest interaction with it, compared to other members of the E3 ubiquitin ligase family^20^. In particular, WWP1 induces K27-linked polyubiquitination of PTEN at residues K342 and K344, impairing PTEN’s dimerization and membrane localization, without degrading the protein. This inactivation of PTEN disrupts its negative regulation of the PI3K/AKT pathway, thereby fostering unrestrained cell survival and proliferation^20^. The same authors tested whether pharmacologically inhibiting WWP1 would affect sensitivity to PI3K (a growth-promoting signal factor) inhibitors in both the original and resistant cell lines^21^. Their results showed that the WWP1 inhibition had a synergistic effect when combined with PI3K inhibitors like BKM120 or BYL719 in suppressing cell growth. Notably, this effect was more pronounced in the resistant MDA-MB-231 cells of BC with higher levels of WWP1 expression. In contrast, only a negligible synergistic effect was observed in MDA-MB-468 cells, which are known to be deficient in the tumor suppressor gene PTEN^21^. Overall, these results strongly suggest that the WWP1/PTEN pathway plays a critical role in both sensitivity to and acquired resistance to PI3K inhibitors^21^. Indeed, the prolonged treatment with PI3K inhibitors often results in acquired resistance, presenting a major challenge in clinical applications^22–24^. Recent studies have further elucidated WWP1’s role in metastasis. In luminal breast cancer, characterized by ER-positive status, WWP1-mediated PTEN polyubiquitination promotes bone tropism, leading to early bone metastasis by inhibiting PTEN membrane recruitment and dimer formation^25^.

Collectively, these findings position pharmacological WWP1 inhibition as an innovative strategy for reactivating tumor suppression pathways in certain cancers. However, to date, the only known WWP1 inhibitors are indole-3-carbinol (I3C), a natural compound present in cruciferous vegetables, and its structural analogs^20,26^. Although I3C is reported to bind WWP1 with a K_d_ of approximately 446 nM, reflecting high binding affinity, it exhibits low target selectivity and exerts pleiotropic effects beyond WWP1 inhibition. In fact, I3C is considered a dietary adjuvant, influencing a wide array of cellular processes including oxidative stress, inflammation, cell proliferation, differentiation, apoptosis, angiogenesis, and immune modulation^27–30^.

In this work, we employed classical and artificial intelligence (AI)-driven computational approaches to design novel WWP1 inhibitors endowed with a peptide-based structure. Following biophysical binding studies and TR-FRET enzymatic assays, we identified a lead peptide with high nanomolar binding affinity to WWP1, representing the first prototype for structure-guided development of anticancer WWP1 inhibitors. Notably, this peptide displayed cytotoxic activity in BC cell lines (MCF-7 and MDA-MB-231), with enhanced efficacy at higher concentrations and synergistic effects in combination with the PI3K inhibitor BYL719, outperforming I3C under comparable conditions.

## 2. MATERIALS AND METHODS

### 2.1. Computational system setup

To create the computational model of the target, the WWP1 X-ray structure was retrieved from the Protein Data Bank with PDB accession code 5HPS, which contains the HECT domain from residues I541 to E917^31^. The system also contains the “ubiquitin variant P1.1” (chain B) bound to WWP1, however, this component was not considered in this study. The protein structure was refined using the Protein Preparation Wizard in Maestro (release 2021-2, Schrödinger, LLC, New York, USA). This procedure included assigning protonation states at pH 7.4, filling in missing side chains, filling the missing loop composed by G605, L606, and D607 residues, resolving steric clashes, optimizing hydrogen bond assignments, and applying the OPLS4 force field for energy minimization.

### 2.2. Computational design of WWP1-inhibiting peptides

Aiming at identifying new peptides endowed with WWP1 inhibiting activity, a tetrapeptide library was initially built by means of an in-house developed python script. This was designed to reduce the number of peptides resulting from the combination of all natural amino acids (20^4^ = 160,000). In particular, the developed script generated all random sequence peptide with at least a tryptophan (Trp, W) residue to mimic the structure of the known inhibitor I3C (**Figure 1**), excluding also the ones containing repetitive amino acids. Specifically, the sequences should contain no more than two tryptophan residues, which cannot be adjacent, and no amino acid should be repeated three times consecutively. Moreover, to avoid the creation of intramolecular bonds and to reduce the possibility of peptide self-cyclization, the *N*- and *C*-termini were protected by acetylation and amidation, respectively. These rules allowed us to dramatically reduce the number of generated peptides from 160,000 to 28,481 tetrapeptides. Then, three approaches were applied to predict their affinity to WWP1 enzyme.

*1) Classical approach utilizing docking, MD simulations, and MM-GBSA calculations.* The whole peptide library (28,481) was docked into the WWP1 catalytic site using the “peptide docking” protocol of GLIDE tool of Maestro (release 2021-2, Schrödinger, LLC, New York, USA), acquiring a glide score (*Gscore*) for each peptide composing the library^32^. The binding site corresponds to that described by Lee et al., who used computational analyses to predict that I3C binds in the hydrophobic pocket surrounded by F577, Y628, L630, N650, and Y656 residues^20^. The docking protocol was validated through re-docking of I3C. The best 100 peptides ranked by Gscore were simulated in complex with WWP1 by 250 ns-long MD simulations (Supporting Information, **Table S1**). MM-GBSA calculations were performed to estimate the peptides binding free energy (ΔG) values considering the last 500 frames (i.e., 100 ns) of MD simulation. The 10 peptides showing the ΔG values equal or lower than – 30 kcal/mol were selected for extending their MD simulations to 500 ns, to better sample the conformational space of the WWP1/peptide complexes and attain more robust prediction. Finally, the peptides showing ΔG values lower or equal than –40 kcal/mol were selected and subjected to an additional independent 500 ns-long MD simulation (Replica 2), in order to increase the statistical significance of the computational studies.
*2) Application of a deep learning (DL) algorithm.* 500 peptides, randomly selected among the whole peptide library developed (28,481 tetrapeptides), were docked in the predicted I3C binding site of WWP1 protein^20^. Then, 150 among the best and worst tetrapeptides (ranked by Gscore) constituted the training set used to instruct a deep learning algorithm (DEEPCHEM tool of Maestro, Schrödinger, LLC, New York, USA) capable of generating a prediction model of the Gscore of the whole 28,481 members library. The DL algorithm was trained for 4 h, using the random split method, and finally the Gscores of the whole peptide library of 28,481 tetrapeptides were predicted (r^2^ = 0.54). Among these, the 200 peptides with the lowest Gscore values were docked into the WWP1 target protein using GLIDE (release 2021-2, Schrödinger, LLC, New York, USA). From the most promising candidates, 10 peptides were selected for a single 500 ns-long MD simulation in complex with WWP1 and their ΔG values were calculated using MM-GBSA approach.
*3) Application of the “Peptide QSAR” tool.* In this protocol, the machine learning (ML) based “Peptide QSAR” tool, implemented in Maestro (release 2021-2, Schrödinger, LLC, New York, USA), was used to generate two prediction models, using the Gscore obtained after “peptide docking” calculations of 500 peptides randomly selected among the whole tetrapeptide’s library: i) the best and worst 50 peptides by Gscore resulted in a model with r^2^ = 0.44 and q^2^ = 0.60; ii) the best and worst 25 peptides selected by Gscore resulted in a model with r^2^ = 0.63 and q^2^ = 0.70. Finally, an MD simulation of 500 ns and MM-GBSA calculations were accomplished to estimate the ΔG values of the best scored peptides of both subsets.

**Figure 1.**
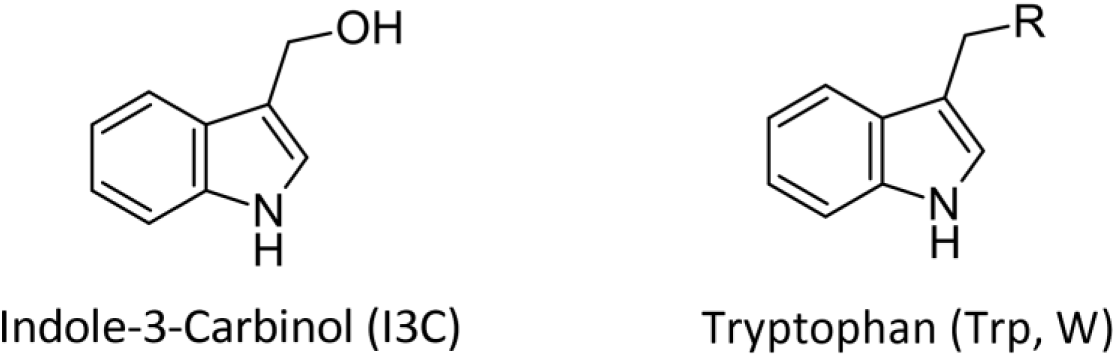
Comparison of the chemical structures between I3C (left) and the Trp amino acid residue (right).

### 2.3. Molecular Dynamics (MD) simulations

The WWP1/peptide complex model was immersed in a water box using the tleap script of Amber22 package and the ff14SB AMBER^33^ force field was applied to the protein, while the TIP3P model represented the explicit water molecules^34^. Van der Waals and short-range electrostatic interactions were computed within an 8 Å cutoff, and long-range electrostatic interactions were incorporated using the particle mesh Ewald method^35^. Hydrogen atom-related bonds were constrained using the SHAKE algorithm, enabling a 2 fs time step. After an initial solvent-only minimization run, the WWP1 model underwent geometric optimization, starting with side chain conformations and progressing to the entire protein structure. Prior to starting the production run of MD simulations, the system was equilibrated at 300 K in isocore conditions (NVT) for 40 ps, followed by isothermal-isobaric ensemble simulations at 300 K. MD simulations were performed using the pmemd.cuda module of Amber22 under periodic boundary conditions and MD trajectories were visually examined using VMD software^36^. The last 1000 MD trajectory frames, corresponding to 200 ns of MD simulation, were utilized for binding energy calculations using the “single trajectory” protocol of the MM-GBSA approach^37^. This protocol employs an ensemble of structures (snapshots) to account for multiple possible conformations of the WWP1/peptide complex using the MM-PBSA.py module^38^ of the Amber22 package, with parameters kept at default values. Entropic contribution was not included in the calculation of the ΔG due to its high computational cost and limited effect on accuracy^39^.

### 2.4. Cluster analysis

The peptide conformations from two independent 500 ns-long MD simulations were clustered considering the backbone atoms and applying the *Gromos* algorithm available in the GROMACS software package^40,41^. After multiple trials and careful evaluation, a suitable root mean square deviation (RMSD) cut-off was chosen to well differentiate peptide conformations while minimizing single-member clusters (Supporting Information, **Table S2**).

### 2.5. Expression and Purification of HECT domain of human WWP1

The *E. coli* strain BL21(DE3)^42^ harbouring the plasmid HECT_WWP1_FF_pET-28a was used for the overexpression of the HECT domain of human WWP1. The sequence of the HECT-domain coding region has been obtained through the retrotranslation of sequence of the protein used for the determination of the 3D structure (PDB accession code 1ND7)^43^. It was optimized for *E. coli* codon usage, synthesized and cloned (GenScript Biotech BV, Leiden, Netherlands) into *Nde*I and *Bam*HI sites of the expression plasmid pET28a(+) (Novagen) to give HECT_WWP1_FF_pET-28a. This plasmid contains the region coding for the 546-917 region of WWP1 (Uniprot accession code Q9H0M0-1) fused at 5’ end to a region coding for the N-terminus His_6_ tag. The cells were cultured in liquid LB medium in the presence of 25 µg/mL kanamycin to A_600_ 0.8 at 37 °C in an orbital shaker (140 rpm), then cooled and induced using 0.5 mM IPTG. After 20 h incubation at 20 °C, cells were harvested by centrifugation (4,000×*g*, 15 min, 4 °C), washed 2-fold with phosphate-buffered saline (PBS; 150 mM NaCl, 15 mM NaH_2_PO_4_, pH 7.2) and stored at -30 °C. For crude extract preparation, thawed collected cells were suspended in 3.3 volumes of degassed extraction buffer (50 mM Tris-HCl, 500 mM NaCl, 1 mM EDTA, pH 8) containing 0.02 mg/mL lysozyme, incubated for 30 min on ice and disrupted by sonication in a Soniprep 150 instrument (MSE, London, UK; 9.5 mm probe), involving seven 30 sec sonication cycles with an amplitude of 16 μm, followed by 2 min on-ice cooling periods. Insoluble debris material was removed using two consecutive 30 min centrifugation periods (16,000×*g* at 4 °C) and the supernatant (i.e., crude extract) was submitted to ion metal affinity chromatography (IMAC) following the method described by Rossi and co-workers^44^ with some modifications that were implemented taking inspiration from works described in some publications on WWP1^43,45^. The IMAC was monitored at 280 nm and was carried out in 50 mM Tris-HCl, 500 mM NaCl (pH 8; running buffer). Elution biphasic linear gradient of imidazole concentration was achieved applying running buffer containing 300 mM imidazole. It was from 0 to 165 mM imidazole in 80 mL and then to 300 mM imidazole in 20 mL. In these conditions, the protein (His_6_-WWP1_HECT_) eluted at 138 ± 3 mM imidazole (at ∼67 mL of the elution gradient). Eluted protein was extensively desalted, exchanged in 50 mM Tris-HCl, 150 mM NaCl (pH 7.4), frozen in liquid nitrogen and stored in aliquots (200-600 µL) at −80 °C. Quality of overexpression and purification was checked by colorimetric protein quantification^46^, using bovine serum albumin as standard, and SDS-PAGE analyses. An amount of 6.8 ± 0.3 mg soluble His_6_-WWP1_HECT_ was yielded per liter of cell culture (∼5 g wet cell pellet).

### 2.6. Peptides

**WI3**, **WI6**, **WI13**, **WI14**, **WI23**, **WI25**, **WI26**, and **WI29** tetrapeptides were acquired from Proteogenix (Schiltigheim, France) which were supplied after trifluoroacetic acid (TFA) removal and with a reported purity of ≥95%. The **WI23-B** and **WI23-E** hexapeptides were obtained from GenScript (Piscataway, NJ, USA), supplied with TFA removed and a reported purity of ≥95%. Instead, **WI6-B** hexapeptide was synthesized *in-house* by the following procedure. Peptide synthesis was carried out on a 2-CTC resin using standard Fmoc-based solid phase protocols, with a synthesis scale of 0.15 mmol. For activation, solutions of 0.5 M Oxyma and 0.5 M DIC were employed, while Fmoc deprotection was accomplished with 20% piperidine in DMF. After sequential coupling steps, the peptides were cleaved from the resin by applying a mixture comprising 92.5% TFA, 2.5% TIS, 2.5% thioanisole, and 2.5% water, followed by precipitation in cold diethyl ether. Crude peptides were isolated via centrifugation and subsequently purified by HPLC using a C-18 Phenomenex column. Unless otherwise indicated, all reagents and materials were sourced from Sigma-Aldrich (St. Louis, MO, USA).

### 2.7. Microscale thermophoresis (MST)

The measurement of binding affinity (K_d_) between the target protein (His_6_-WWP1_HECT_) and peptides was conducted through MST^47^. The protein was labeled using the Monolith His-Tag Labeling Kit RED-tris-NTA 2nd Generation (MO-L018), acquired from NanoTemper Technologies (München, Germany), for 30 min at room temperature (RT). MST experiments were carried out using Monolith NT.115^Pico^ instrument (NanoTemper Technologies, München, Germany). In the preliminary binding check experiments, a fixed 10 nM concentration of the red-labeled His_6_-WWP1_HECT_ protein was mixed with each tetrapeptide at the constant concentration of 100 µM applying the “Expert Mode” available in MO.Control (v1.6; NanoTemper Technologies, München, Germany). I3C, at the concentration of 100 µM, was also tested to serve as positive control^20^. The assay was considered positive for a ligand when the response amplitude (RA), determined as the difference in normalized fluorescence (Fnorm) between the ligand-bound and unbound states of His_6_-WWP1_HECT_, was greater than or equal to 2.5 value. In the binding affinity experiments, a constant concentration of the red-labeled His_6_-WWP1_HECT_ protein was combined with sixteen 1:1 serial dilutions of peptides. The tetrapeptides were evaluated across a concentration range from 156 μM to 0.00477 μM, whereas the hexapeptides were assessed from 100 μM down to 0.0122 μM. The protein-peptide mixture was incubated for 60-90 min at room temperature. MST analysis utilized premium capillaries with specific experimental settings: an excitation power of 20% and medium MST power (40%). The K_d_ values were calculated based on at least two independent experiments, and data analysis was performed using the NanoTemper MO.Affinity Analysis software (v2.3; NanoTemper Technologies, München, Germany) using the K_d_ model for fitting the data. In both binding check and binding affinity experiments, the protein and peptides were dissolved in PBS-T buffer (phosphate-buffered saline + 0.05% Tween™ 20; NanoTemper Technologies, München, Germany) and 2.5% dimethyl sulfoxide (DMSO; Product No. D8418, Sigma-Aldrich, Saint Louis, MO, USA). Each peptide was tested to exclude any auto-fluorescence capable of interfering with K_d_ measurements, adopting the protocol suggested by NanoTemper Technologies.

### 2.8. Surface plasmon resonance (SPR)

SPR experiments were conducted using a Biacore 8K system (Cytiva, Marlborough, MA, USA). His_6_-WWP1_HECT_ (200 nM) in 10 mM sodium acetate (pH 4.5) was immobilized on a CM5 sensor chip (GE Healthcare, Chicago, IL, USA) via standard amine coupling. All the tetrapeptides were injected in a single kinetic experiment at five concentrations (1.56, 3.12, 6.25, 12.5, and 25 µM), over the immobilized His_6_-WWP1_HECT_ at a flow rate of 30 µL/min. In the case of **WI3** and **WI6**, they were tested at concentrations up to 50 μM (3.12, 6.25, 12.5, 25, and 50 μM). Each injection had a contact time of 120 s, followed by a dissociation phase of 600 s. Experiments were performed at 25 °C. In the case of **WI6-B**, **WI23-B**, and **WI23-E** hexapeptides, a single kinetic experiment was performed where the peptides were injected at six concentrations (0.195, 0.39, 0.78, 1.56, 3.125, 6.25, and 12.5 µM) using the same protocol previously described for the tetrapeptides. After blank subtraction, binding data were analyzed and fitted using Biacore Insight Evaluation Software (v5.0.18; Cytiva, Marlborough, MA, USA).

### 2.9. Protein expression and purification for TR-FRET assay

The coding region for the complete HECT domain of WWP1 protein (aa 546-922 of WWP1, Uniprot accession code Q9H0M0-1) was amplified by PCR from a pCR3.1 YFP-WWP1 construct using the following oligos: FOR: gcggatccGGCTTTAGGTGGAAG / REV: TAGAAGGCACAGTCGAGG and cloned into pGEX6P1 vector (Cytiva) using *Bam*HI and EcoRI restriction enzymes. Glutathione S-transferase (GST)-WWP1_HECT_ protein was expressed in *E. coli* Rosetta cells (Novagen). The cells were cultured in liquid LB medium in the presence of 100 µg/mL ampicillin at 37 °C to A600 0.6 in an orbital shaker (140 rpm), then cooled at 18 °C and induced using 0.5 mM IPTG. After 16 h incubation, cells were harvested by centrifugation (4000 ×g, 15 min, 4 °C). Cell pellets were resuspended in lysis buffer (50 mM Na-HEPES, pH 7.5, 200 mM NaCl, 1 mM EDTA, 0.1% NP40, 5% glycerol and Protease Inhibitor Cocktail set III (Calbiochem, San Diego, CA)). Sonicated lysates were cleared by centrifugation at 45,000×g for 45 min. Supernatants were incubated with glutathione-Sepharose beads (Cytiva). After 2 h at 4 °C, beads were washed with lysis buffer followed by a wash with PBS and finally equilibrated in cleavage buffer (20 mM Tris-HCl, pH 7.4, 150 mM NaCl, 1 mM EDTA, 1 mM DTT and 5% glycerol). To cleave off GST, HRV 3C protease (produced in house) was added in a ratio (w/w) 1:50 (protease:substrate) and incubated for 16 h at 4 °C. The cleaved protein were concentrated in Amicon Ultra Centrifugal Filters (MW cut-off 30 kDa) (Merck KGaA, Darmstadt, Germany) and loaded onto a Superdex 200 size exclusion chromatography column (Cytiva, Marlborough, MA, USA) equilibrated with 20 mM Tris-HCl, pH 8.0, 200 mM NaCl, 1 mM EDTA, 5% glycerol, 1 mM DTT. Fractions containing purified protein were collected and concentrated. The E1 (Ube1, Addgene clone 34965), E2 (Ube2D3) enzymes and Ub production and purification were previously described in the paper of Taibi and co-workers^48^.

### 2.10. Ubiquitination assay based on time-resolved fluorescence resonance energy transfer (Ub-TR-FRET) assay

TR-FRET reactions were set up in a white OptiPlate-384 well Microplate (Optiplate, Perkin Elmer, #6007290) using EnVision microplate reader with the following setting: Excitation Filter - UV2 (TRF) 320 nm; Emission Filter 1 - APC 665 nm; Emission Filter 2 - Europium 615 nm; Delay Time 50 μs; Window Time 400 μs. Briefly, enzymes are bacterially expressed, purified and mix at various concentrations (final concentration, 15 nM E1, 350 nM of E2, 200 nM WWP1_HECT_ as E3) together with TRF-Ubiquitin-mix (mixture of Eu-cryptate ubiquitin, Cy5-ubiquitin and wild-type ubiquitin, 100x SouthBayBio) in TR-FRET buffer (25 mM Tris-HCl, pH 7.6, 5 mM MgCl_2_, 100 mM NaCl, 0.01% Tween20), generating a master mix. Peptides were dissolved in 100% DMSO at high concentration and serially diluted in 96-well plates (from 100 µM to 0.005 µM), ensuring that the final DMSO concentration in the reaction mixture never exceeded 2%. 50 µL/well of the master mix is dispensed in a 384 well plate together with 3 µL of diluted peptides or TR-FRET buffer (with the same amount of DMSO) were added and pre-incubated for 1 h. The reaction was initiated with the addition ATP in TR-FRET buffer (2 mM final concentration) and was allowed to proceed for 150 min at RT. Kinetic raw data were analyzed using GraphPad Prism software version 4.0 (GraphPad Software, San Diego, CA). Due to the sigmoidal shape of the curve representing the reaction kinetics, we evaluated the effect of the inhibitors by calculating the area under the curve (AUC) of the reaction. Data are reported as %signal/background, where the background corresponds to the signal obtained from the same enzymatic mix without ATP. IC_50_ values were calculated using GraphPad Prism considering for each tested dose the corresponding AUC. For experiments shown in **Figure 5B**, the reaction products at the end of the measurement were further analyzed using SDS-PAGE followed by anti-ubiquitin immunoblot. Coomassie staining of the membrane is used as a loading control.

### 2.11. Immunoblotting

Proteins were separated by SDS-PAGE (NuPAGE 4-12% Bis-Tris Gel, Invitrogen) and blotted onto PVDF membrane. Filters were denatured for 30 min at 4°C in 6 M Guanidine-HCl pH 7.6, 5 mM ß-mercaptoethanol and 1 mM PMSF prior to blocking in 5% BSA in TBS-T (TBS, 0.1% tween-20) for 16 h. Membranes were incubated with primary anti Ub antibody (anti-Ub, clone ZTA10, generated in house^49^ diluted 1:5 in 5% BSA in TBS-T for 1 hour at RT followed by an incubation for 30 min at RT with horseradish peroxidase-conjugated secondary antibody in TBS-T (Goat anti-mouse IgG HRP, Bio-Rad® dilution 1:10000). Protein detection was carried out using the ECL Select™ Western Blotting Detection Reagent (Cytiva Amersham™), and signals were visualized using the Chemidoc Imaging System (Bio-Rad®). To verify equal protein loading, filters were stained with ImperialTM Protein Stain (Thermo Scientific®) and used as a loading reference.

### 2.12. Cell lines

Human BC cells MCF-7 (luminal A subtype) and MDA-MB-231 (triple negative subtype) were kindly provided by Prof. Rinner from the Medical Graz University (Austria) and cultured in DMEM (Sigma Aldrich, St. Louis, Missouri, USA) supplemented with 1% L-Glutamine (Aurogene, Rome, Italy), 1% penicillin/streptomycin (Aurogene, Rome, Italy) and 10% fetal bovine serum (FBS, Sigma Aldrich, St. Louis, Missouri, USA).

### 2.13. Cell viability assay (I3C, WI23-B and WI23-E)

MCF-7 and MDA-MB-231 cells were seeded at a concentration of 1 × 10^4^ cells/well on a 96 well plate and maintained under cell culture conditions overnight. The following day, cells were treated as follows: *(i)* cell culture medium; *(ii)* BYL719 (0, 0.2, 0.4, 0.6, 0.8, 1, 1.2 μM) and I3C (0, 40, 80, 120, 160, 200 μM) in combination; *(iii)* BYL719 (0, 0.2, 0.4, 0.6, 0.8, 1, 1.2 μM) and **WI23-B** or **WI23-E** (0, 40, 80, 120, 160, 200 μM) in combination. 72 h post treatment MTS cytotoxicity assay was performed according to the manufacturer’s protocol using CellTiter96 AQueous One Solution Cell Proliferation Assay (Promega, Madison, Wisconsin, USA). Spectrophotometric analysis was performed through absorbance recording using Viktor^®^ Nivo^™^ Plate Reader at 490 nm (Perkin Elmer, Waltham, MA, USA). Experiments were performed in triplicates for each condition.

### 2.14. Synthesis of empty nanocapsules (empty-NCs) and peptide loaded nanocapsules

The breakable hybrid organosilica nanocapsules (NCs) were synthesized using the Stöber process at room temperature in a W/O microemulsion (following the protocol developed by Prasetyanto et al.^50^) prepared by mixing 1.77 mL of TRITON X-100, 7.5 ml of cyclohexane and 1.8 mL of n-hexanol in a 50 mL round-bottom flask and stirring at 250 rpm on a magnetic stirrer for 30 min. Separately, 1.1 mg of **WI23-B** or 1.2 mg of **WI23-E** were dissolved in 600 μL of water, then mixed with 40 μL of tetraethyl orthosilicate (TEOS) and 60 μL of bis[3- (triethoxysilyl)propyl]disulfide. After shaking, this mixture was added to the organic solution. The hydrolysis of TEOS was initiated by the addition of 50 μL of aq. NH_3_ 28 wt%, and the mixture was stirred at room temperature overnight. 20 mL of acetone was subsequently added to precipitate the ssNCs and the material was recovered by centrifugation. All the round bottom flask content is poured in a centrifuge Teflon tube, then centrifuged for 30 min, 4 °C at 35,000 rcf. After removing the supernatant, 2 mL of ethanol were added in the tube, then they are sonicated for about 2 min. All the Teflon tubes content is poured in an empty Eppendorf tube and centrifuged for 30 min at 14,100 rcf, then the supernatant is removed. This washing passage was performed twice with ethanol and twice with water. Finally, the empty-NCs and peptide loaded NCs (**WI23-B@NC** and **WI23-E@NC**) were stored at 4°C in water/EtOH 1:1.

### 2.15. Cell viability assay (empty-NCs, WI23-B@NC and WI23E@NC)

The same procedure described in chapter 2.13. was employed, with exception that cells were treated as follows: (i) cell culture medium; (ii) BYL719 (0, 0.2, 0.4, 0.6, 0.8, 1, 1.2 μM) in combination with empty-NCs that were used as controls and tested at 0 mg/mL, 0.25 mg/mL, 0.75 mg/mL and 1.25 mg/mL corresponding approximately to 0, 40, 120 and 200 µM of peptide equivalents; *(iii)* BYL719 (0, 0.2, 0.4, 0.6, 0.8, 1, 1.2 μM) in combination with peptide-loaded nanocapsules (**WI23-B@NC**, **WI23-E@NC**) at concentration of 0, 40, 120, 200 μM.

### 2.16. Synthesis of the carboxyfluorescein(CF)-labeled peptide WI23-B

Microwave-assisted automated peptide synthesis was performed using the Fmoc/tBu protection group strategy on a Rink amide resin with a loading capacity of 0.59 mmol/g with a Liberty Blue synthesizer using a scale of 0.1 mmol. The amino acids concentration was equal to 0.2 M in DMF. DIC and Oxyma were used as coupling reagents (respectively 0.5 and 1 M in DMF) while for the deprotection 20% piperidine in DMF was used. Couplings were performed at 75 °C using 170 W for 15 seconds and then at 90 °C using 40 W for 110 seconds and a double coupling was performed for both Arg residue and the Fmoc-O_2_Oc-OH linker. Deprotection was performed at 75 °C using 155 W for 15 seconds and then at 90 °C using 50 W for 50 seconds. The coupling with 5(6)-carboxyfluorescein (CF) was performed using 10 equivalents of CF, 10 of DIC and 10 of Oxyma. The reaction was left shaking for 2 h in the dark. Then the resin was washed (DMF, DCM, DMF) and finally 20% piperidine was added for 45 min to depolymerize the carboxyfluorescein (CF) polymer. The cleavage was then performed using 3 mL of cleavage cocktail for **WI23-B** (trifluoroacetic acid/triisopropylsilane(TIPS)/thioanisole/H_2_O/phenol; 82:5.5:5:3) for 4.5 h at room temperature. After the cleavage, the peptide was precipitated from ice-cold diethyl ether and recovered by centrifugation at 4 °C. The obtained **CF-[WI23-B]** was purified using a Kinetex C-18 column from Phenomenex (10 µm, 250 × 21.2 mm) by RP-HPLC using a gradient elution of 05-60% solvent B (solvent A: water/trifluoroacetic acid 100: 0.1; solvent B: acetonitrile/trifluoroacetic acid 100:0.1) over 20 min at a flow rate of 20 mL/min. The purified **CF-[WI23-B]** were freeze-dried and stored at 0°C. Afterwards, it was analyzed using analytical HPLC (5% solvent B for 3 min; 05-60% solvent B over 20 min) and ESI mass spectrometry (ESI-MS). The UV chromatogram and ESI-MS spectrum are available in the Supporting Information.

### 2.17. Encapsulation of CF-[WI23-B] into Cy5-functionalized nanocapsules (Cy5-NCs)

The same procedure described in chapter 2.14. was employed for the encapsulation of 1.1 mg of **CF-[WI23-B]**. For the Cy5 functionalization of the NCs, 0.5 mg of Cy5-NHS ester was dissolved in 200 µL of anhydrous DMSO in a 2 mL Eppendorf tube. Subsequently, 4.5 µL of APTES was added to the DMSO solution, and the reaction mixture was stirred for 30 min at room temperature in the dark. Then, 30 µL of the resulting mixture was added to the aqueous **CF-[WI23-B]@Cy5-NC** suspension and stirred overnight at 400 rpm in the dark. After the reaction, the ssNCs were centrifuged twice in 2 mL of Milli-Q water, twice in 2 mL of ethanol, and twice in a 1:1 (v/v) ethanol/Milli-Q water solution. Finally, the **CF-[WI23-B]@Cy5-NC** were resuspended in Milli-Q water.

### 2.18. Confocal microscopy studies

MCF-7 cells were seeded onto 24-chamber glass slides (Sigma-Aldrich, St. Louis, MO, USA) at a density of 1×10⁵ cells/well in complete growth medium and allowed to adhere overnight. The following day, cells were treated with increasing concentrations (0, 40, 120, 200 μM) of CF-labeled WI23-B either as free peptide (**CF-[WI23-B]**) or encapsulated into Cy5-labeled silica nanocapsules (**CF-[WI23-B]@NC**). Untreated cells served as negative controls. After 24 and 48 h of incubation, cells were washed three times with PBS and fixed with 4% (w/v) paraformaldehyde in PBS for 15 min at RT. Following fixation, cell nuclei were stained with Hoechst (Thermo Fisher Scientific, Waltham, MA, USA) for 10 min at 37 °C. Cells were washed three times with PBS between each staining step. The chambered coverslips were then mounted onto microscope slides (Menzel-Gläser, Braunschweig, Germany) using ProLong™ Glass Antifade Mountant (Invitrogen, Thermo Fisher Scientific, Waltham, MA, USA). Confocal imaging was performed using a Zeiss LSM800 laser scanning confocal microscope equipped with a 63× oil immersion objective. Fluorescence was sequentially acquired using 405 nm, 488 nm and 640 nm laser lines for Hoechst, CF and Cy5 excitation, respectively. Image analyses were performed using ImageJ software^51^.

## 3. RESULTS AND DISCUSSION

To identify new peptide-based WWP1 inhibitors, we constructed a library of tetrapeptides with random sequences, each containing at least one Trp residue, due to its structural similarity to I3C (**Figure 1**).

The design of the most promising WWP1 inhibitors was carried out by applying three different computational procedures, spanning from the classical approach of only docking and MD simulations to the application of AI algorithms (see Material and Methods).

### 3.1. Computational design of WWP1-inhibiting peptides by classical approach

By the classical approach, the whole peptide library was docked into the WWP1 binding site, and the best 100 peptides, according to Gscore, were subjected to MD simulations and MM-GBSA calculations, to estimate their ΔG values (see Material and Methods for details). **Table 1** displays the 10 most promising peptides, designated by the abbreviation “WI”, which stands for “WWP1 Inhibitors”. The complete list of all the 100 tetrapeptides subjected to a single 250-ns long MD simulation is reported in the Supporting Information, **Table S1**.

**Table 1.** Binding free energy (ΔG) of the 10 most promising peptides out of all the 100 peptides simulated.

| Peptide | Sequence | $\Delta G_{\text{Rep1}} \pm \text{SEM}^1$<br>(0-250 ns) | $\Delta G_{\text{Rep1}} \pm \text{SEM}^1$<br>(250-500 ns) | $\Delta G_{\text{Rep2}} \pm \text{SEM}^1$<br>(0-500 ns) | Average<br>$\Delta G^1$ |
| --- | --- | --- | --- | --- | --- |
| WI1 | Ace-TRYW-NMe | $-32.0 \pm 0.2$ | $-30.7 \pm 0.3$ | / | / |
| WI2 | Ace-QYYW-NMe | $-30.3 \pm 0.1$ | $-30.6 \pm 0.2$ | / | / |
| WI3 | Ace-TWYE-NMe | $-37.7 \pm 0.3$ | $-41.2 \pm 0.2$ | $-36.8 \pm 0.3$ | $-39.0$ |
| WI4 | Ace-WQYL-NMe | $-34.9 \pm 0.3$ | $-42.1 \pm 0.1$ | unbound | / |
| WI5 | Ace-RRWE-NMe | $-39.3 \pm 0.3$ | $-31.3 \pm 0.3$ | / | / |
| WI6 | Ace-SWYT-NMe | $-40.3 \pm 0.2$ | $-40.2 \pm 0.2$ | $-41.1 \pm 0.3$ | $-40.7$ |
| WI7 | Ace-SCWQ-NMe | $-30.5 \pm 0.2$ | unbound | / | / |
| WI8 | Ace-WQHR-NMe | $-39.2 \pm 0.4$ | $-44.3 \pm 0.4$ | unbound | / |
| WI9 | Ace-NWRW-NMe | $-32.4 \pm 0.4$ | $-34.2 \pm 0.3$ | / | / |
| WI10 | Ace-IHYW-NMe | $-38.4 \pm 0.2$ | $-43.2 \pm 0.2$ | unbound | / |

<sup>1</sup> kcal/mol; SEM = Standard Error of Mean.

As shown in **Table 1**, only two peptides (namely, **WI3** and **WI6**) exhibited appreciable ΔG values and maintained stability at the active site throughout the MD simulations, which was corroborated in two independent replicas. Consequently, only these two peptides were selected and acquired for further experimental evaluation.

### 3.2. Computational design of WWP1-inhibiting peptides by AI

On parallel to the classical computational pipeline, two new different protocols, characterized by a significant lower computational time cost, were adopted. In these approaches (approaches 2 and 3, Material and Methods section), the Gscore of 500 peptides, randomly selected in the previous peptide library, was used to generate ML or DL prediction models capable of guessing the Gscore of the complete peptide library. These protocols, with varying performance, allowed the prediction of peptide’s Gscore without performing computationally demanding docking calculations on the entire peptide library. By the application of DL algorithm (approach 2), only two peptides out of the 10 most promising peptides by AI-predicted Gscore, namely **WI13** and **WI14**, showed appreciable ΔG values of –35.8 and –42.2 kcal/mol, respectively (**Table 2**). Consequently, also in this case only these two peptides were acquired for further experimental assays. To note, we deliberately performed only a single MD simulation for these AI-based predictions, fully embracing the rationale of integrating AI to minimize computational time and cost.

**Table 2.** Binding free energy (ΔG) of the 10 most promising peptides using the DeepChem-based approach.

| Peptide | Sequence | AI-Predicted Gscore<sup>1</sup> | $\Delta G_{\text{Rep1}} \pm \text{SEM}^1$<br>(0-500 ns) |
| --- | --- | --- | --- |
| **WI11** | Ace-SQYW-NMe | -8.57 | *unbound* |
| **WI12** | Ace-YNKW-NMe | -8.55 | *unbound* |
| **WI13** | Ace-WRQY-NMe | -8.55 | $-35.8 \pm 0.3$ |
| **WI14** | Ace-QWRY-NMe | -8.45 | $-42.2 \pm 0.4$ |
| **WI15** | Ace-WKYN-NMe | -8.45 | *unbound* |
| **WI16** | Ace-KQYW-NMe | -8.44 | *unbound* |
| **WI17** | Ace-WYWQ-NMe | -8.43 | $-19.9 \pm 0.1$ |
| **WI18** | Ace-WFYQ-NMe | -8.42 | *unbound* |
| **WI19** | Ace-SWYQ-NMe | -8.39 | $-21.4 \pm 0.2$ |
| **WI20** | Ace-WNKY-NMe | -8.37 | *unbound* |

<sup>1</sup> kcal/mol; SEM = Standard Error of Mean.

By the application of “Peptide QSAR” algorithm (approach 3), the best two peptides retrieved from each subset (namely, **WI23**, **WI25**, **WI26**, and **WI29**) that displayed the lowest ΔG values together with the highest stability over two independent MD simulations (**Table 3**), were selected to be purchased for further experimental assays.

**Table 3.** Binding free energy (ΔG) of the 10 most promising peptides using two different Peptide QSAR models.

| Peptide | Sequence | AI-Predicted Gscore<sup>1</sup> | $\Delta G_{\text{Rep1}} \pm \text{SEM}^1$<br>(0-500 ns) |
| --- | --- | --- | --- |
| **WI21** | Ace-RKWE-NMe | -8.08 | *unbound* |
| **WI22** | Ace-WRKH-NMe | -7.95 | *unbound* |
| **WI23** | Ace-RKWQ-NMe | -7.90 | $-28.0 \pm 0.3$ |
| **WI24** | Ace-KRWN-NMe | -7.89 | *unbound* |
| **WI25** | Ace-YRWQ-NMe | −7.87 | −25.5 ± 0.3 |
| **WI26** | Ace-FWYC-NMe | −8.51 | −38.7 ± 0.3 |
| **WI27** | Ace-WFYC-NMe | −8.24 | *unbound* |
| **WI28** | Ace-CYRW-NMe | −8.04 | −31.3 ± 0.3 |
| **WI29** | Ace-YWKH-NMe | −7.96 | −36.7 ± 0.3 |
| **WI30** | Ace-WYKC-NMe | −7.91 | −19.0 ± 0.3 |

<sup>1</sup> kcal/mol; SEM = Standard Error of Mean.

In conclusion, by the computational approaches described before, eight peptides were selected and acquired from Proteogenix (Schiltigheim, France) for further experimental assays, aiming at identifying new peptides endowed with WWP1 inhibiting activity.

### 3.3. Biophysical experiments

Preliminary microscale thermophoresis (MST) “binding check” assays were conducted on the chemically synthesized eight peptides and I3C (positive control) at a fixed concentration of 100 µM (see Materials and Methods). Interestingly, all peptides displayed Fnorm values that significantly differed from that of His_6_-WWP1_HECT_ alone, indicating appreciable interactions with the target under these conditions (Supporting Information, **Figure S1**). To further quantify these interactions, MST “binding affinity” experiments were subsequently performed on the entire synthetic peptide library to determine their K_d_ values. Nevertheless, among all tested peptides, only **WI23** yielded a clear, complete and unambiguous binding curve, showing a K_d_ value of 4.5 ± 1.0 µM (**Figure 2A**). The full MST analysis report of **WI23** is available in the Supporting Information, **Figure S2**. Then, surface plasmon resonance (SPR) experiments were also performed to further validate and determine the K_d_ values of the peptides, yielding results consistent with those obtained from MST binding check assays. In fact, SPR data confirmed that all peptides bound to His_6_-WWP1_HECT_ with K_d_ values in the micromolar range (Supporting Information, **Figure S3**). However, closer examination of the SPR sensorgrams revealed that the observed K_d_ values are greater than the peptide concentration range tested. Notably, only in the case of **WI6** it was observable a complete binding curve in the concentration range examined, displaying a K_d_ of 6.4 µM (**Figure 2B**).

**Figure 2.**
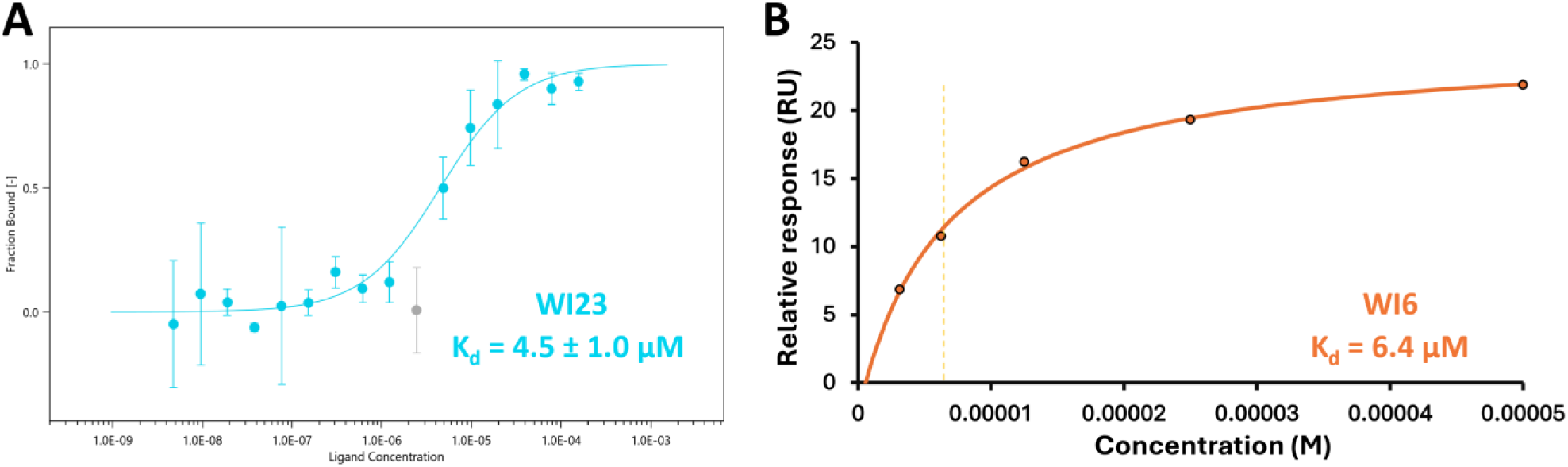
(A) MST binding affinity curve of the peptide **WI23** performed in two independent replicates. The seventh concentration point (2.44 µM), shown in grey, was excluded from K_d_ analysis as it represents a clear outlier. **(B)** SPR affinity plot of peptide **WI6**. The K_d_ value is highlighted in the graph and indicated by light orange dashed lines.

### 3.4. Design of new WI6 and WI23 analogues

Considering the promising binding affinity displayed by **WI6** and **WI23** peptides, additional computational investigations were done to identify new peptides with improved affinity on WWP1. Firstly, cluster analysis was performed to identify the most representative structure of the complex during the MD simulations (see Materials and Methods). For **WI23**, an additional independent MD simulation of 500 ns was carried out to further enhance conformational sampling and thus improve the statistical robustness of the results, while, for **WI6**, two independent MD simulations were already completed prior to this analysis. The representative conformations of the most populated cluster comprised the 74% and 90% of the generated conformational ensemble of **WI23** and **WI6**, respectively. In the case of **WI23**, the residue W_3_ remains firmly anchored within the hydrophobic pocket and establishes multiple hydrophobic interactions with residues F577, Y628, C629, N650, and Y656 of the WWP1 protein. In addition, it is capable of forming an H-bond with S647 as well as a π-π stacking interaction with Y656. Regarding the other residues, R_1_ and K_2_ establish H-bonds and salt bridges with the carboxylate side chains of E578 and D652, respectively, whereas Q_4_, together with its protected *C*-terminus tail, remains fully exposed to the solvent (**Figure 3A**).

**Figure 3.**
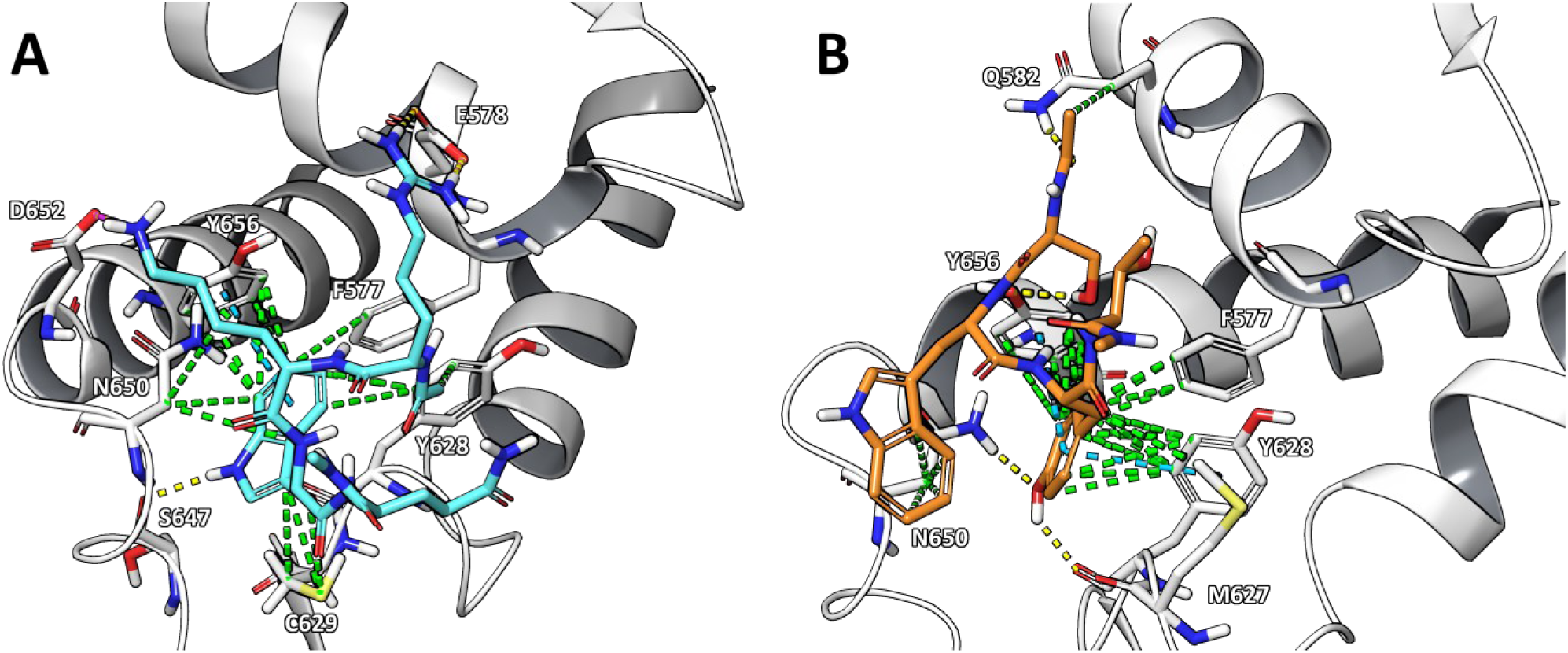
The representative structure of the most populated cluster, considering two independent 500 ns-long MD simulations, of **(A) WI23**, and **(B) WI6**, which account for 74% and 90% of the total conformational ensemble, respectively. H-bond, salt bridges, π-cation and hydrophobic interactions are shown as yellow, purple, cyan and green, respectively. WWP1 protein is represented as light grey cartoon and sticks.

In the case of **WI6**, the peptide shifts slightly from its original position. Within the hydrophobic pocket, residue Y_3_, replacing W_2_, occupies the binding site and forms multiple hydrophobic interactions with F577, Y628, and Y656, as well as π-π stacking interactions with the latter two residues. The hydroxyl group of Y_3_ functions as a H-bond acceptor with N650 and as a donor to the backbone carboxyl group of M627. This observation suggests that peptides enriched in tyrosine residues may represent promising scaffolds for the development of novel WWP1 inhibitors. Among the remaining residues, only S_1_ establishes a hydrogen bond with Q582, whereas W_2_ and T_4_ remain largely solvent-exposed. W_2_ nonetheless engages in limited hydrophobic contacts with the side chain of N650 (**Figure 3B**). All the interactions occurring between the peptides and WWP1 protein are shown in the Supporting Information, **Figure S4**.

Based on the analysis of the 3D binding modes and interactions occurring between the peptides and the WWP1 protein, in both cases, we decided to extend the peptide sequence at the *N*-terminus in order to reach additional residues, previously inaccessible by the tetrapeptides, thereby potentially increasing the number of interactions and consequently increasing binding affinity. Specifically, the peptide sequences were extended by adding two glycine residues at the *N*-terminus with the aim of reaching new protein regions. For **WI23**, the extension was designed to reach: (i) a cavity formed by E578, F581, Q582, and the hydroxyl group of Y656, located close to the hydrophobic pocket; and (ii) a small pocket surrounded by residues Q574, E622, N625, and M627 (Supporting Information, **Figure S5A**). For **WI6**, the *N*-terminal extension was aimed to establish new contacts mainly with D579 and R601 (Supporting Information, **Figure S5B**). Subsequently, both Gly residues were systematically replaced by all natural amino acids using the “affinity maturation protocol” available in the Bioluminate software within Maestro (release 2021-2, Schrödinger, LLC, New York, USA), and the mutants were ranked according to their predicted ΔAffinity values (**Table 4**). This protocol has been extensively validated in our prior published studies, demonstrating robust performance and substantially enhancing the binding affinity of the peptides towards the target^52,53^. Finally, the most promising hexapeptides were simulated in complex with WWP1 through two independent 500 ns-long MD simulations replicates, and their ΔG values were estimated by MM-GBSA approach (**Table 4**).

**Table 4.** ΔG values of the new hexapeptides attained by the application of the affinity maturation protocol.

| Peptide | Sequence | ΔAffinity<sup>1</sup> | ΔStability<sup>1</sup> | ΔG<sub>Rep1</sub> ± SEM<sup>1</sup><br>(0-500 ns) | ΔG<sub>Rep2</sub> ± SEM<sup>1</sup><br>(0-500 ns) | Average<br>ΔG<sup>1</sup> |
| --- | --- | --- | --- | --- | --- | --- |
| **WI6** | Ace-SWYT-NMe | / | / | $-40.2 \pm 0.2$ | $-41.1 \pm 0.3$ | $-40.7$ |
| **WI6-A** | Ace-GYSWYT-NMe | -9.58 | +0.43 | *unbound* | *unbound* | / |
| **WI6-B** | Ace-GRSWYT-NMe | -4.36 | +1.58 | $-52.0 \pm 0.5$ | $-49.5 \pm 0.3$ | $-50.7$ |
| **WI6-C** | Ace-GMSWYT-NMe | -2.86 | +0.38 | $-30.6 \pm 0.2$ | *unbound* | / |
| **WI23** | Ace-RKWQ-NMe | / | / | $-28.0 \pm 0.3$ | $-30.7 \pm 0.3$ | $-29.3$ |
| **WI23-A** | Ace-RWRKWQ-NMe | -32.33 | -10.16 | *unbound* | $-33.2 \pm 0.4$ | / |
| **WI23-B** | Ace-RFRKWQ-NMe | -27.81 | -6.38 | $-49.5 \pm 0.3$ | $-45.4 \pm 0.4$ | $-47.5$ |
| **WI23-C** | Ace-RMRKWQ-NMe | -27.47 | -0.87 | $-40.8 \pm 0.2$ | $-28.6 \pm 0.3$ | $-34.7$ |
| **WI23-D** | Ace-WGRKWQ-NMe | -11.56 | -6.57 | *unbound* | *unbound* | / |
| **WI23-E** | Ace-RSRKWQ-NMe | -10.03 | +5.49 | $-42.7 \pm 0.2$ | $-46.9 \pm 0.2$ | $-44.8$ |
| **WI23-F** | Ace-QARKWQ-NMe | -4.60 | +1.52 | *unbound* | *unbound* | / |
| **WI23-G** | Ace-NARKWQ-NMe | -2.01 | -3.22 | *unbound* | *unbound* | / |

<sup>1</sup> kcal/mol; SEM = Standard Error of Mean.

As shown in **Table 4**, the peptide **WI6-B** exhibited an average ΔG value approximately 10 kcal/mol lower than its parent peptide **WI6**, while the **WI23** analogues, **WI23-B** and **WI23-E**, showed reductions of 18.2 and 15.5 kcal/mol, respectively, compared to **WI23**. Consistent with these predictions, these peptides were synthesized and subjected to further biophysical examination.

### 3.5. Hexapeptides characterization

SPR experiments revealed well-defined affinity curves for both **WI23-B** (K_d_ = 920 nM, **Figure 4A**) and **WI23-E** (K_d_ = 625 nM, **Figure 4B**), indicating strong interactions with WWP1 in the sub-micromolar range. Interestingly, only in the case of **WI23-B** it was observed a clear kinetic profile, from which a K_d_ of 1.3 µM was derived (**Figure 4C**). Unexpectedly, no binding curve was detected for **WI6-B**, suggesting either a lack of interaction under the tested conditions or a binding mode not adequately captured by the SPR setup. To corroborate these findings, additional MST experiments were performed. The results were fully consistent with the SPR data, in fact, both **WI23-B** (**Figure 4D**, and Supporting Information, **Figure S6**) and **WI23-E** (Supporting Information, **Figure S7**) displayed comparable K_d_ values in the high nanomolar range, while no measurable binding was observed for **WI6-B**.

**Figure 4.**
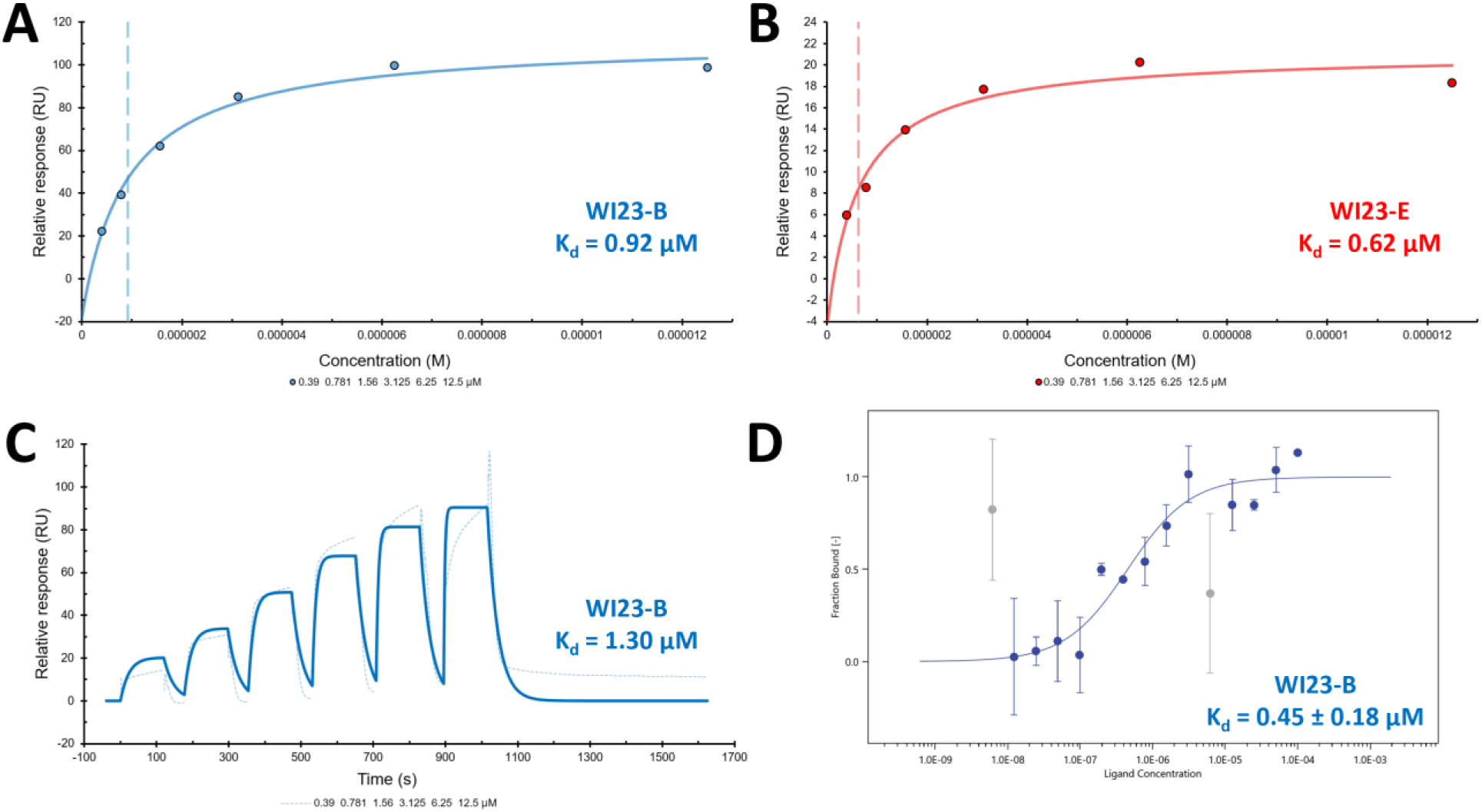
(A-B) SPR affinity plots of **WI23-B** and **WI23-E**, respectively, on His_6_-WWP1_HECT_. (**C**) SPR kinetic plot of **WI23-B** on His_6_-WWP1_HECT_. (**D**) MST binding curve of **WI23-B** on His_6_-WWP1_HECT_ in two independent replicas, the grey points were discarded since they could be considered as outliers.

**Figure 5.**
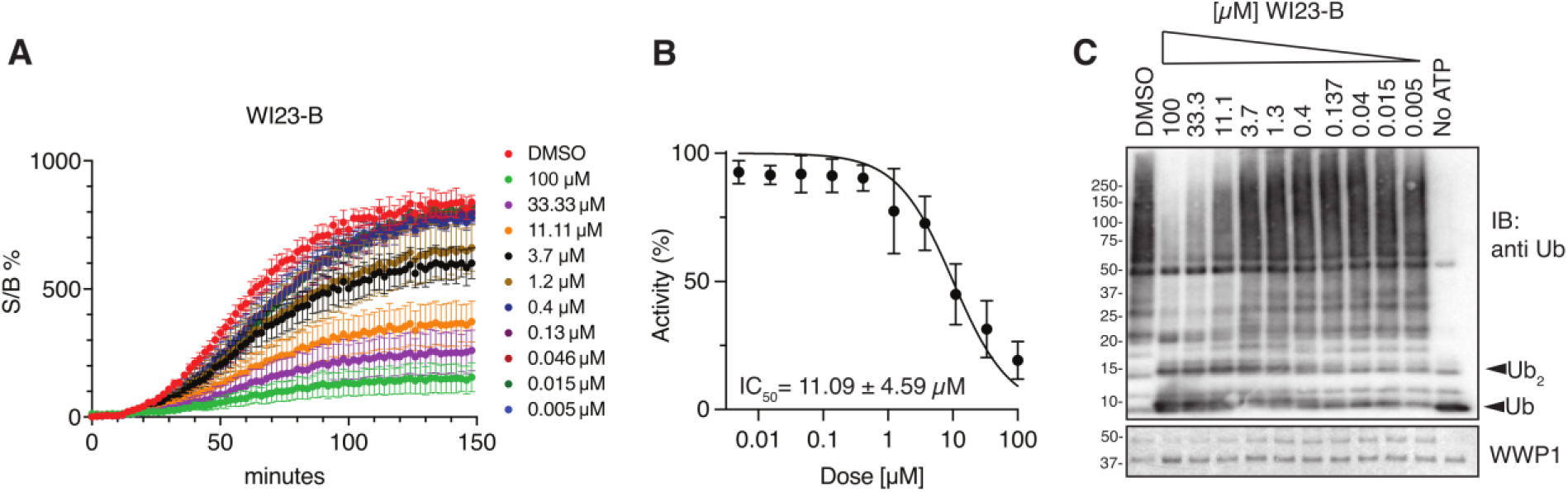
**(A)** Ub-TR-FRET assay with WWP1_HECT_ pre-incubated with an increasing concentration of **WI23-B** peptide for 60 min before ATP addition and FRET reaction measurement. Plots show single time-point measurements of FRET signal to background ratio (% Signal/Background) as function of time (minutes) (mean ± SEM, n=2). **(B)** Dose-response curve of WWP1_HECT_ activity in presence of **WI23-B** peptide. The values are expressed as percentage of activity, calculated as percentage of the area under the curve (% AUC) normalized to DMSO treated sample at the different doses. Black circles represent calculated values of two independent experiments (mean ± SEM). Calculated IC_50_ value is 11.09 ± 4.59 µM. **(C)** Immunoblot analysis of the samples collected at 150 min of TR-FRET reaction after Laemmli buffer addition. Top panel, mono-, di- and polyUb are indicated. Bottom panel, Coomassie staining showing comparable loading of WWP1_HECT_ protein.

In conclusion, although **WI23-E** demonstrated a slightly higher affinity than **WI23-B** in biophysical assays, only **WI23-B** displayed a clear kinetic profile in SPR assays and a distinctly binding curve in MST measurements. Therefore, between the two peptides, only **WI23-B** was chosen for evaluation of WWP1 inhibitory activity via TR-FRET experiments. This assay measures ubiquitin-chain formation by the ubiquitin (Ub) machinery (E1, E2, E3) using a mix of fluorescently labelled and unlabelled monomeric Ub that result in a FRET signal increase as consequence of the Ub chain formation and elongation. After determining the optimal enzyme concentration to achieve the highest signal-to-background ratio, **WI23-B** peptide inhibitory profile was tested across a concentration range from 0.005 to 100 µM (**Figure 5A**). The IC₅₀ was calculated by correlating the AUC for each tested dose with the percentage of signal relative to the DMSO-treated control, yielding an IC₅₀ of 11.09 ± 4.59 µM (**Figure 5B**). At this concentration, immunoblot analysis further confirmed reduced polyubiquitin smear intensity alongside an accumulation of monomeric Ub species (**Figure 5C**).

### 3.6. In vitro evaluation of WI23-B and WI23-E in BC models

Previous studies have reported that BC prognosis and disease progression are closely associated with WWP1 expression, which negatively regulates the tumor suppressor PTEN^16^. Consequently, to evaluate the *in vitro* efficacy of **WI23-B** and **WI23-E**, MTS cytotoxic assays were performed on two representative BC cell lines: the luminal A subtype MCF-7 and the triple-negative subtype MDA-MB-231. These two models were selected to capture the heterogeneity of BC and to assess whether the activity of the peptides varies according to the molecular subtype^54^. Moreover, the cytotoxic effects of **WI23-B** and **WI23-E** were assessed both alone and in combination with increasing concentrations of BYL719 (**Figure 6B,C**), a selective inhibitor targeting the alpha isoform of the PI3K enzyme, which is currently undergoing clinical trials for the treatment of various solid tumors, including BC^55^. These data were also compared with those obtained testing the reference compound I3C under the same experimental conditions (**Figure 6A**)^21^. The obtained data demonstrated that increasing concentrations of BYL719 alone produced concentration-dependent cytotoxic effects on both BC cell lines, as expected (**Figure 6**, black line). When treated with escalating concentrations of the individual compounds in the absence of BYL719, we observed distinct dose-dependent reductions in cell viability across both cell lines. I3C decreased viability of MCF-7 cells to about 50% at 120 µM, whereas the same concentration reduced viability to `∼`70% in MDA-MB-231 cells (**Figure 6A**). In contrast, **WI23-B** elicited a markedly stronger cytotoxic effect, particularly in the TNBC model: at 120 µM, cell viability declined to `∼`41% in MCF-7 and `∼`53% in MDA-MB-231, indicating a more potent activity relative to I3C under equivalent conditions (**Figure 6B**). In contrast, **WI23-E** displayed weaker cytotoxic activity, with viability values of `∼`71% in MCF-7 and ∼79% in MDA-MB-231 at 120 µM. Notably, in this assay, both cell lines were intrinsically less responsive to BYL719 alone at the highest concentration tested (1.2 µM; **Figure 6C**, black line), showing viability levels of approximately 60% in MCF-7 and 70% in MDA-MB-231. These values are markedly higher than those observed for I3C and WI23-B under comparable conditions (40-50% for MCF-7 and 45-65% for MDA-MB-231), further underscoring the relatively modest activity of **WI23-E**. Interestingly, at the highest concentration tested (200 µM), both I3C and **WI23-B** markedly reduced cell viability in MCF-7 cells to `∼`20%. In MDA-MB-231 cells, **WI23-B** remained slightly more potent than I3C, lowering viability to `∼`23% compared with `∼`30% for I3C (**Figure 6A,B**). In contrast, **WI23-E** exhibited substantially weaker activity at the same concentration, reducing cell viability only to `∼`42% in MCF-7 and `∼`48% in MDA-MB-231 (**Figure 6C**).

**Figure 6.**
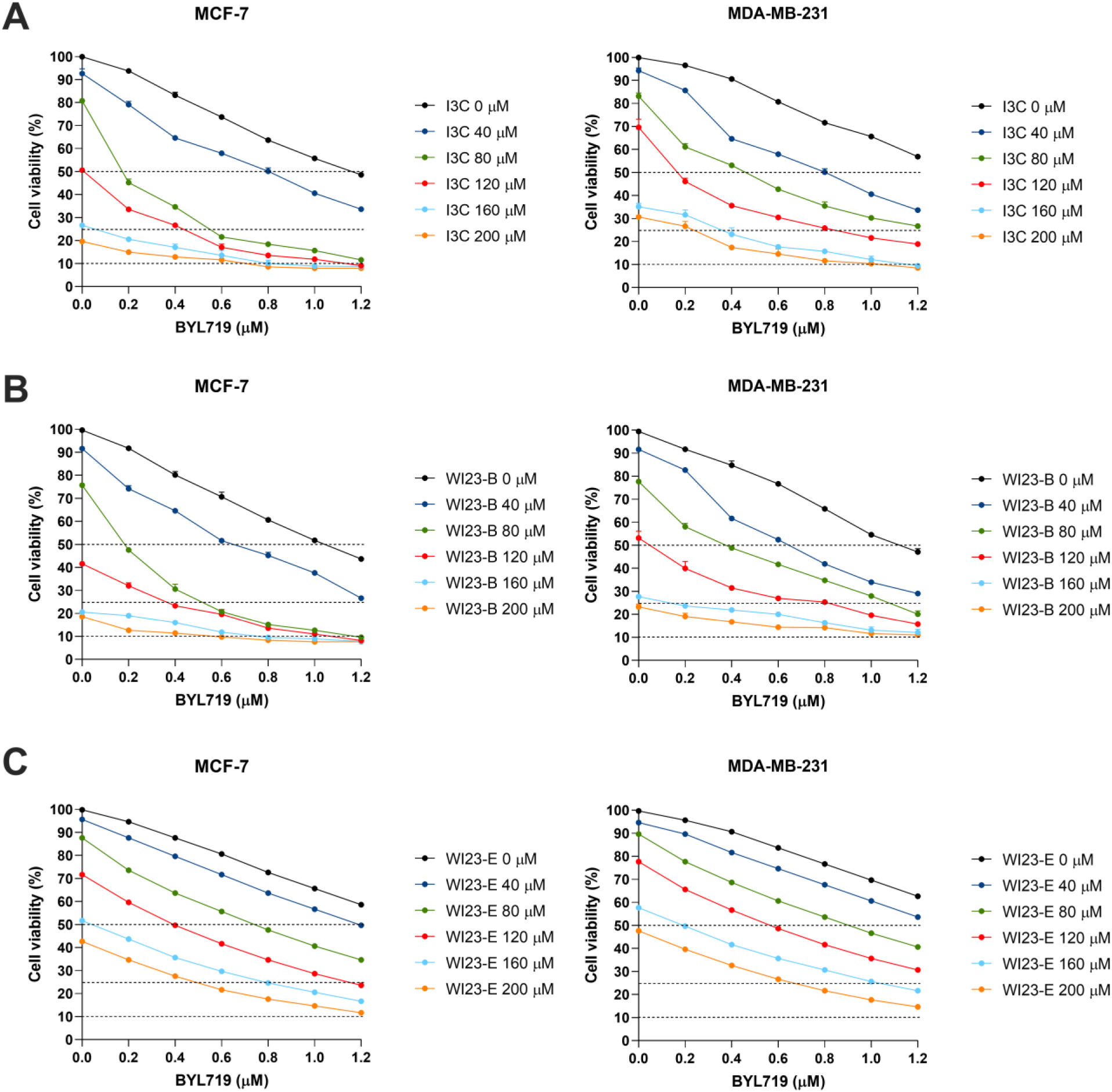
Evaluation of combination treatments using an MTS cytotoxicity assay for I3C, **WI23-B**, and **WI23-E** in combination with BYL719, assessing their effects on cell viability in BC cell lines. Cell viability of MCF-7 and MDA-MB-231 cells treated with increasing concentrations of BYL719 (0, 0.2, 0.4, 0.6, 0.8, 1 and 1.2 µM) in combination with different concentrations (0, 40, 80, 120, 160 and 200 µM) of the compounds **(A)** I3C, **(B) WI23-B** and **(C) WI23-E**. Analysis was performed 72 h after treatment. Data are expressed as the percentage of viable cells, determined using CellTiter 96 Aqueous One Solution Cell Proliferation Assay (MTS) and measuring the absorbance at 490 nm with a 96-well plate spectrophotometer Viktor Nivo^TM^.

In the case of the compound/BYL719 combinations, a pronounced synergistic cytotoxic effect was observed. At the highest concentrations tested (200 µM for the compounds and 1.2 µM BYL719), cell viability consistently decreased to about 10% across both cell lines, indicating a substantial enhancement of the antiproliferative activity relative to single-agent treatments (**Figure 6**). Co-administration with BYL719 substantially reduced the amount of antitumoral agent required to achieve comparable cytotoxic effects, thereby potentially limiting treatment-associated side effects. Notably, when considering the concentration of BYL719 needed to reduce cell viability by 50%, **WI23-B** enabled a marked dose-sparing effect in MCF-7 cells: the same level of inhibition obtained with 1 µM BYL719 alone was achieved either with 40 µM **WI23-B** plus 0.6 µM BYL719 (a 40% reduction) or with 80 µM **WI23-B** plus 0.2 µM BYL719 (an 80% reduction). A similar trend was observed in the TNBC model, where 40 µM **WI23-B** allowed a 46% reduction in BYL719 concentration (0.6 µM instead of 1.1 µM), and 80 µM **WI23-B** enabled a 64% reduction (0.4 µM instead of 1.1 µM), as shown in **Figure 6B**. Comparable but slightly weaker dose-sparing effects were observed for I3C (**Figure 6A**). In contrast, **WI23-E** did not allow a reliable estimation of this parameter, as BYL719 alone did not reach 50% inhibition at the highest concentration tested (**Figure 6C**).

In MDA-MB-231 cells, the overall response was reduced, consistent with the lower sensitivity of this triple-negative model; however, both peptides maintained a detectable effect combined with BYL719 (**Figure 6B,C**). Furthermore, the cytotoxic effect of **WI23-B** was very similar to that of the natural compound I3C, suggesting that **WI23-B** could operate through the same mechanism of action as I3C, by inhibiting WWP1-mediated dimerization of PTEN, which in turn impede PTEN’s tumor-suppressive function^20^. This mechanism, combined with its ability to potentiate the PI3Kα inhibition, makes both peptides promising therapeutic candidates for fighting BC^21^.

### 3.7. Synthesis of peptide-loaded nanocapsules (NCs)

Silica nanoparticles are widely recognized as versatile drug delivery platforms, thanks to their high surface area, tunable porosity, and straightforward functionalization chemistry^56^. In this work, the synthesized organosilica nanocapsules consist of a hydrophilic core, in which the peptides are encapsulated, surrounded by a denser silica shell that provides structural integrity and protection of the payload until cellular internalization. In addition, the incorporation of disulfide bridges within the silica framework confers redox-responsive degradability, enabling controlled and sustained release of the encapsulated species^56^. Within this context, the peptides **WI23-B** and **WI23-E** were encapsulated in organosilica nanocapsules (NCs) to assess their performance as delivery systems (see Materials and Methods section for details) (**Figure 7**). To allow tracking of the NCs internalization and the intracellular peptide release, **WI23-B** was conjugated to carboxyfluorescein (CF, see Materials and Methods for details) and the NCs containing **CF-[WI23-B]** were further functionalized with Cy5 (**CF-[WI23-B]@Cy5-NC**), to enable *in vitro* studies of nanoparticles uptake and intracellular localization (**Figure 7**).

**Figure 7.**
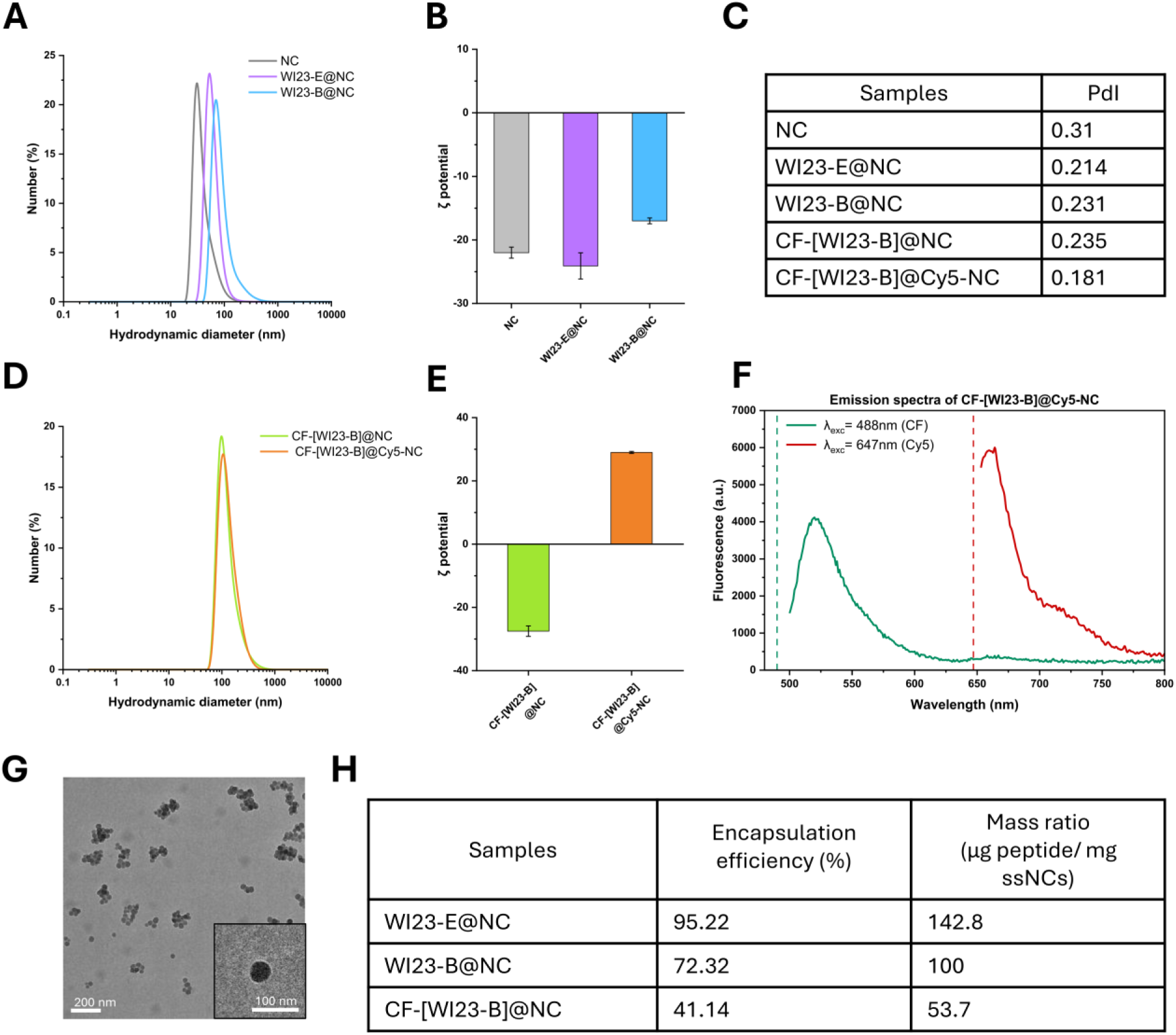
Physicochemical characterization of empty and peptide-loaded organosilica nanocapsules**. (A)** Hydrodynamic diameter of empty-NCs and NCs loaded with **WI23-E** or **WI23-B**, and **(B)** corresponding ζ-potential measurements. **(C)** Polydispersity index (PDI) of all formulations. **(D)** Hydrodynamic diameter of **CF-[WI23-B]@NC** before and after post-synthesis Cy5 functionalization, with **(E)** corresponding ζ-potential measurements. **(F)** Emission spectra of **CF-[WI23-B]@Cy5-NC** under excitation at 488 nm and 647 nm, confirming successful incorporation of CF and Cy5. (G) TEM image of **CF-[WI23-B]@NC**. **(H)** Encapsulation efficiencies and mass ratio for all the formulations.

The hydrodynamic diameter of NCs increased with the encapsulation of WI23-E and WI23-B, reaching approximately 80 nm and 90 nm, respectively, compared to empty-NCs control (**Figure 7A**). All formulations exhibited a strongly negative ζ-potential, with a slight reduction in magnitude for **WI23-B@NC** (**Figure 7B**).

Encapsulation of **WI23-B** conjugated with CF (**CF-[WI23-B]@NC**) resulted in a hydrodynamic diameter comparable to the one measured for **WI23-B@NC**, indicating that CF labeling did not affect nanoparticles size. Following the peptide encapsulation, Cy5 was conjugated on the outer surface of NCs (**CF-[WI23-B]@Cy5-NC**), resulting in a slight increase in the hydrodynamic diameter and a shift in the surface charge toward positive ζ-potential values, likely due to residual protonated amine groups from the APTES linker that are not fully shielded by Cy5 conjugation (**Figure 7D,E**). All samples showed low polydispersity indices ranging from 0.18 to 0.31, consistent with homogeneous and well-dispersed nanoparticles population (**Figure 7C**).

Fluorescence characterization further confirmed successful labeling. The emission spectra recorded from the **CF-[WI23-B]@Cy5-NC** under sequential excitation at 488 nm and 647 nm selectively activated CF and Cy5, respectively, with no evidence of cross-excitation, confirming the coexistence of both fluorophores within the system (**Figure 7F**).

Consistently, TEM analysis of **CF-[WI23-B]@NC** confirmed the expected nanoparticles morphology and revealed an average particle size of approximately 50 nm (**Figure 7G**). Finally, the encapsulation efficiencies and corresponding mass ratios for all the formulations are reported in **Figure 7H**.

### 3.8 In vitro evaluation of WI23-B@NC and WI23-E@NC in BC models

To further investigate whether the enhanced cytotoxic activity observed for **WI23-B** could be translated into a nanocapsule-based delivery system, we assessed the effects of novel **WI23-B** and **WI23-E**-loaded nanocapsules in combination with increasing concentrations of BYL719 in both MCF-7 and MDA-MB-231 cell lines (**Figure 8**). As shown, treatment with empty-NCs did not significantly affect cell viability in either model, confirming their compatibility and suitability as delivery systems. The reduction in cell viability observed in these conditions is mainly attributable to the activity of BYL719 alone, since increasing concentrations of empty-NCs did not substantially alter the cytotoxic profile (**Figure 8A**, black line). Indeed, cell viability remained relatively stable across all empty-NCs concentrations, demonstrating that the nanocapsule itself is not intrinsically toxic and does not exert antiproliferative effects in the absence of peptide cargo.

**Figure 8.**
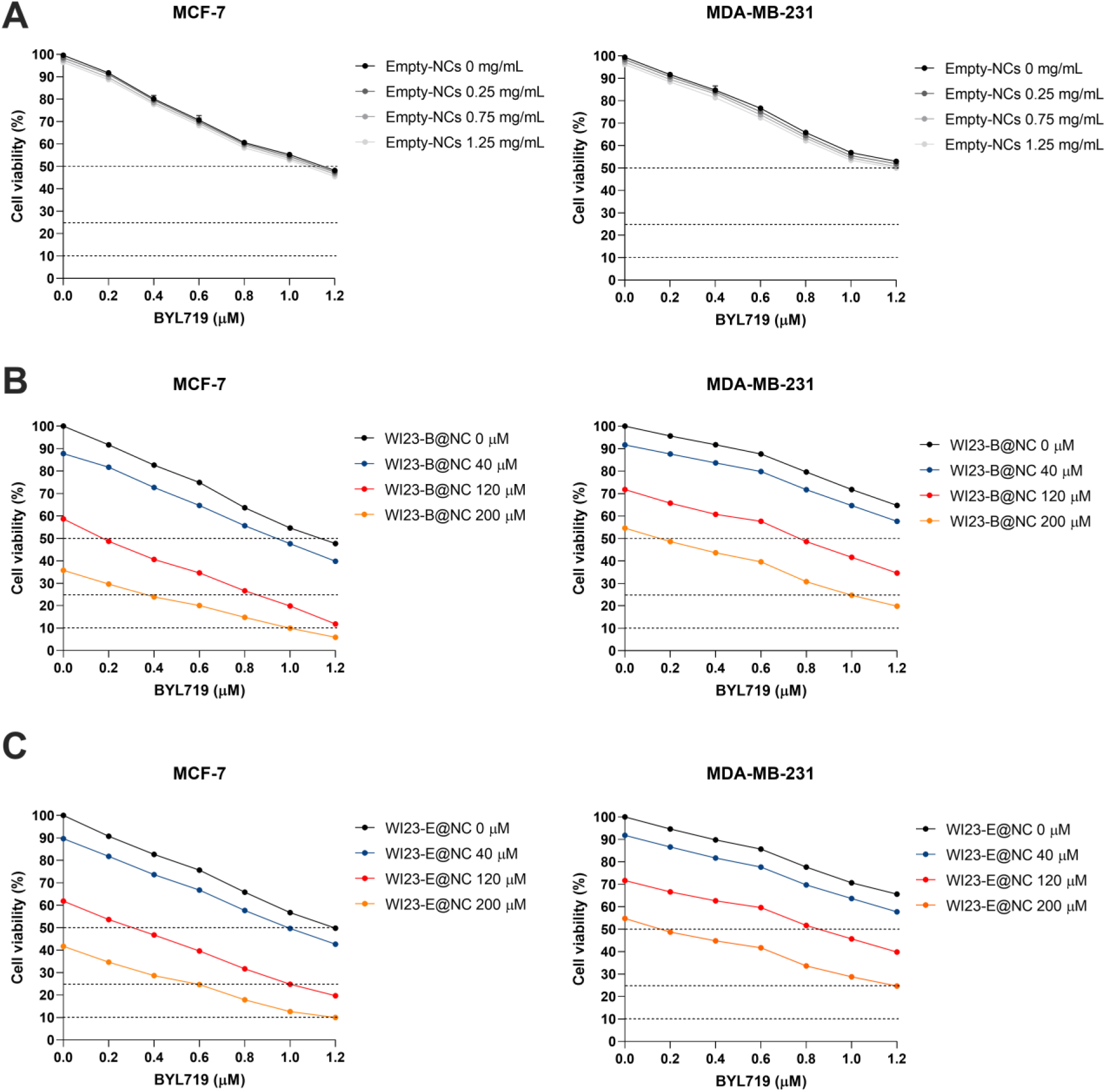
Effect of peptide-loaded nanocapsules in combination with BYL719 on cell viability in BC cell lines. Cell viability of MCF-7 and MDA-MB-231 cells treated with increasing concentrations of BYL719 (0, 0.2, 0.4, 0.6, 0.8, 1 and 1.2 µM) in combination with different concentrations of nanocapsule-loaded peptides (0, 40, 120, and 200 µM). **(A)** Empty-NCs used as control at a concentration of 0 mg/mL, 0.25 mg/mL, 0.75 mg/mL and 1.25 mg/mL which corresponds to approximately 0, 40, 120 and 200 µM of peptide equivalents. **(B) WI23-B** loaded nanocapsules (**WI23-B@NC**). **(C) WI23-E** loaded nanocapsules (**WI23-E@NC**). Cell viability is expressed as percentage relative to untreated controls. Analysis was performed 72 h after treatment. Data are expressed as the percentage of viable cells, determined using CellTiter 96 Aqueous One Solution Cell Proliferation Assay (MTS) and measuring the absorbance at 490 nm with a 96-well plate spectrophotometer Viktor Nivo^TM^.

In MCF-7 cells, both **WI23-B@NC** and **WI23-E@NC** induced a clear dose-dependent reduction in cell viability, which was further enhanced upon co-treatment with BYL719. Among the tested nanoformulations, **WI23-B@NC** showed a slightly stronger antiproliferative effect than **WI23-E@NC**, in fact, without BYL719, it reduced cell viability to `∼`59% and `∼`35% at 120 µM and 200 µM, respectively (**Figure 8B**), compared with

`∼`62% and `∼`42% for **WI23-E@NC** (**Figure 8C**). The reduced impact on cell viability observed for the nanoencapsulated peptide relative to the free peptide (**Figure 6**) can be explained by the intrinsic release kinetics of the NCs. These formulations require a defined temporal window to liberate their payload and allow the active peptide to accumulate intracellularly. Because both treatments were evaluated at the same 72 h endpoint, the nanoformulation likely delivered its cargo more gradually, resulting in a lower effective concentration of the peptide during the assay period. However, in the case of the combination with BYL719 at higher concentrations, both peptides maintained a cytotoxic activity relatively close to that of the free peptide (**Figure 6B,C**), indicating that encapsulation preserves a substantial fraction of their biological efficacy (**Figure 8B,C**). This suggests that once the NCs have released a sufficient amount of peptide, an event that requires a defined temporal window, the combined treatment can reach intracellular concentrations comparable to those achieved with the free peptide, thereby sustaining a similar antiproliferative response.

In MDA-MB-231 cells, a similar trend was observed, with **WI23-B@NC** showing a slightly stronger antiproliferative effect than **WI23-E@NC**, by approximately 5%, particularly when combined with BYL719 at higher concentrations (**Figure 8B,C**). The overall response to the peptide-loaded NCs was more attenuated, both when administered alone and in combination with BYL719, consistent with the intrinsically lower sensitivity of this TNBC model compared with MCF-7 (**Figure 8**). However, even in this model, co-administration with BYL719 at the highest concentrations led to a pronounced reduction in cell viability, reaching `∼`20% and `∼`25% for **WI23-B@NC** and **WI23-E@NC**, respectively (**Figure 8B,C**). It is worth noting that, in this case, the cells appear less responsive at baseline. Indeed, BYL719 alone reduced cell viability only to `∼`70% (**Figure 8**, black line), whereas in the previous experiments yielded values of `∼`50% and `∼`60% (**Figure 6**, black line).

Overall, these results demonstrate that NC loading preserves the biological activity of **WI23-B** and **WI23-E** while enabling their effective combinatorial use with PI3Kα inhibition. **WI23-B@NC** emerged as the most potent formulation in both cell lines, retaining strong antiproliferative effects and maintaining substantial activity after encapsulation. Nanocapsule systems are designed to enhance peptide stability, protect the cargo from premature degradation, and modulate its release profile, features that may become particularly advantageous in more complex biological environments or *in vivo* settings, where controlled delivery can improve pharmacokinetics and therapeutic persistence. Collectively, these findings reinforce the concept that targeting the WWP1/PTEN axis sensitizes BC cells to PI3Kα inhibition. By relieving WWP1-mediated PTEN suppression, the peptides restore or enhance PTEN activity, thereby strengthening the negative regulation of PI3K signaling. As a consequence, cells become more vulnerable to pharmacological PI3Kα blockade, displaying a markedly amplified antiproliferative response when the two interventions are combined. This cooperative effect highlights the central role of PTEN reactivation in reshaping PI3K pathway dependency and provides mechanistic support for therapeutic strategies that pair WWP1 inhibition with PI3Kα-targeted agents.

### 3.9. Cellular uptake of CF-[WI23-B] and CF-[WI23-B]@Cy5-NC in MCF-7 cells

To prove the intracellular delivery of **WI23-B** and to better understand the differences observed between the free and nanocapsule-loaded peptide in the cytotoxicity assays, confocal microscopy analyses were performed in MCF-7 cells, the most responsive BC model (**Figure 9**). Cells were incubated with increasing concentrations of **CF-[WI23-B]** either as free peptide or encapsulated into Cy5-conjugated nanocapsules (**CF-[WI23-B]@Cy5-NC**), and fluorescence was monitored after 24 and 48 h. These time point were selected based on previous studies from De Cola et al., demonstrating degradation of silica nanocapsule platform after endocytic internalization, ultimately promoting intracellular cargo release^57^.

**Figure 9.**
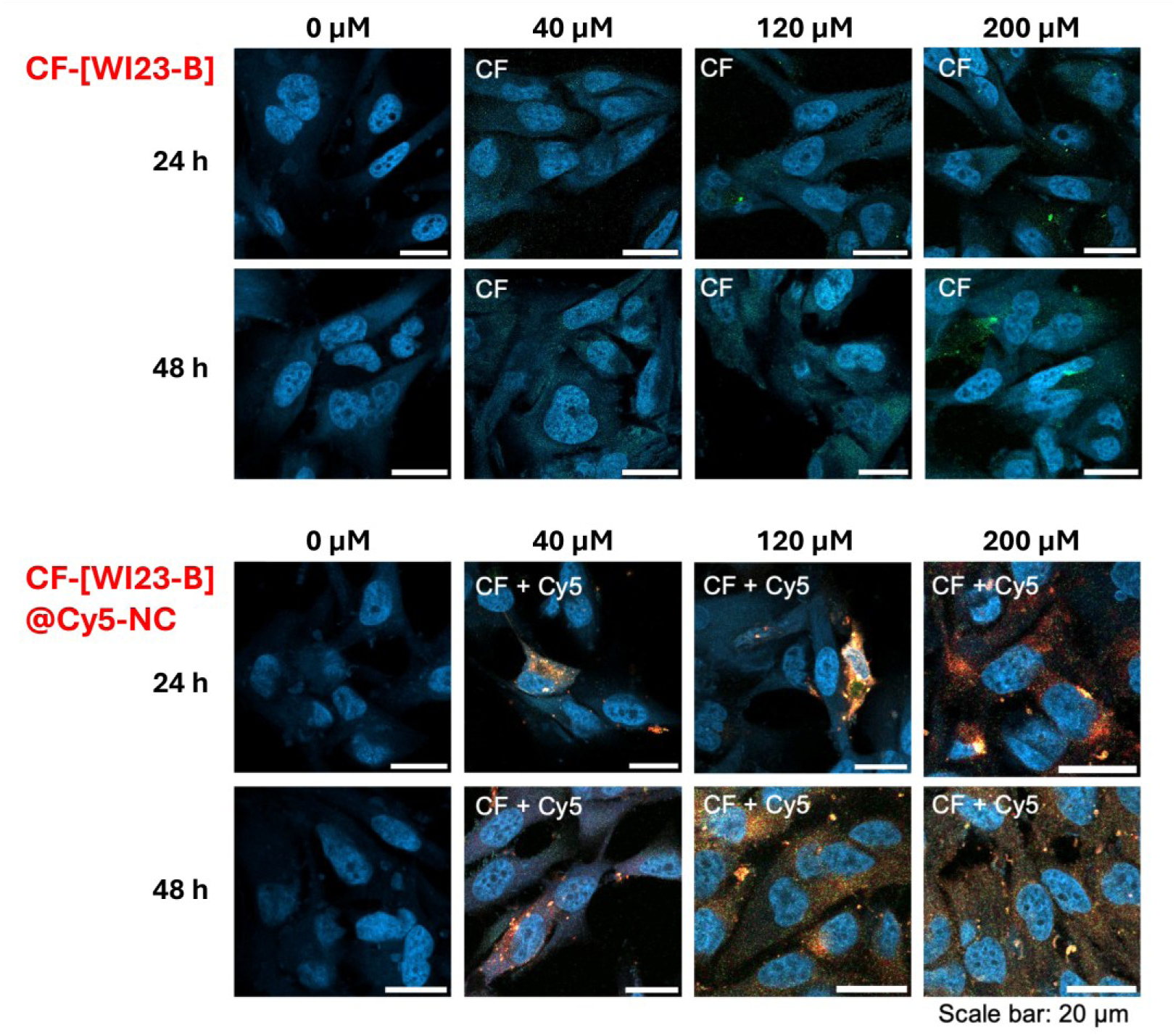
Intracellular uptake of free WI23-B peptide and WI23-B-loaded nanocapsules in MCF-7 cells evaluated by confocal microscopy. MCF-7 cells were incubated with increasing concentrations (0, 40, 120, 200 µM) of CF-labeled **WI23-B** either as free peptide (upper panels) or encapsulated into Cy5-labeled silica nanocapsules (**CF-[WI23-B]@Cy5-NC**, lower panels). Images were acquired after 24 and 48 h of treatment. Nuclei were stained with Hoechst (blue), free peptide fluorescence is shown in green (CF), whereas **CF-[WI23-B]@Cy5-NC** were visualized by merged CF (green) and Cy5 (red) signals. Scale bar: 20 µm.

As shown in **Figure 9**, intracellular fluorescence was detected in cells treated with both free **CF-[WI23-B]** and **CF-[WI23-B]@Cy5-NC**. However, important differences in intracellular distribution were observed, aligning with the related cytotoxicity. The free peptide produced a relatively weak and diffuse CF signal throughout the cytoplasm, with only a modest increase at higher concentrations and longer incubation times. In contrast, **CF-[WI23-B]@Cy5-NC** exhibited a markedly stronger intracellular fluorescence that progressively increased. The intense Cy5 signal confirmed the efficient cellular internalization of the nanocapsules, while overlapping CF fluorescence demonstrated successful intracellular delivery of the encapsulated peptide.

Overall, these observations demonstrate that peptide encapsulation markedly enhances the intracellular accumulation of **WI23-B** compared with the free peptide. The progressive increase in CF fluorescence and overlapping with Cy5-NC signal supports the hypothesis of a gradual intracellular release of the peptide following endocytic uptake. These findings confirm that the developed nanocarrier efficiently transports **WI23-B** into BC cells and represents a suitable delivery systems for intracellular peptide administration.

### 3.10. WI23-B binding mode and hints for designing new analogues

To depict the binding mode of **WI23-B** in complex with WWP1, a clustering procedure was applied to the trajectories obtained by two independent 500 ns-long MD simulations. The representative structure of the most populated cluster, accounting on about the 74% of the sampled conformations, was selected for the interaction’s description (**Figure 10**). As can be noted in this figure, the W_5_ residue of **WI23-B** was deeply and precisely positioned within the hydrophobic pocket, establishing extensive hydrophobic interactions with residues F577, Y628, L630, N650, and Y656. In addition, the aromatic bicyclic ring of W_5_ created π-π stacking interactions with Y628. Both R_1_ and R_3_ arginine residues created strong H-bond-assisted salt bridges with the side chains E622 and E578, respectively; notably, R_3_ establishes also an additional H-bond with E578 through its backbone. The amide group of Q_6_’s side-chain established two H-bonds with N650 and D652 while the side chain of residue F_2_ was projected close to the hydrophobic area shaped by F577, Y628, and Y656. Moreover, it created additional hydrophobic contacts with E578 and F581, establishing a broad interaction network not previously observed in the case of the tetrapeptide series. The same F_2_ residue established an H-bond by means of the backbone -NH group with the hydroxyl group of Y628, improving the conformational stability of the peptide. Only the residue K_4_ of **WI23-B** was fully solvent-exposed and devoid of any intermolecular interactions with WWP1. However, its polarity can be advantageous to enhance the peptide water solubility.

**Figure 10.**
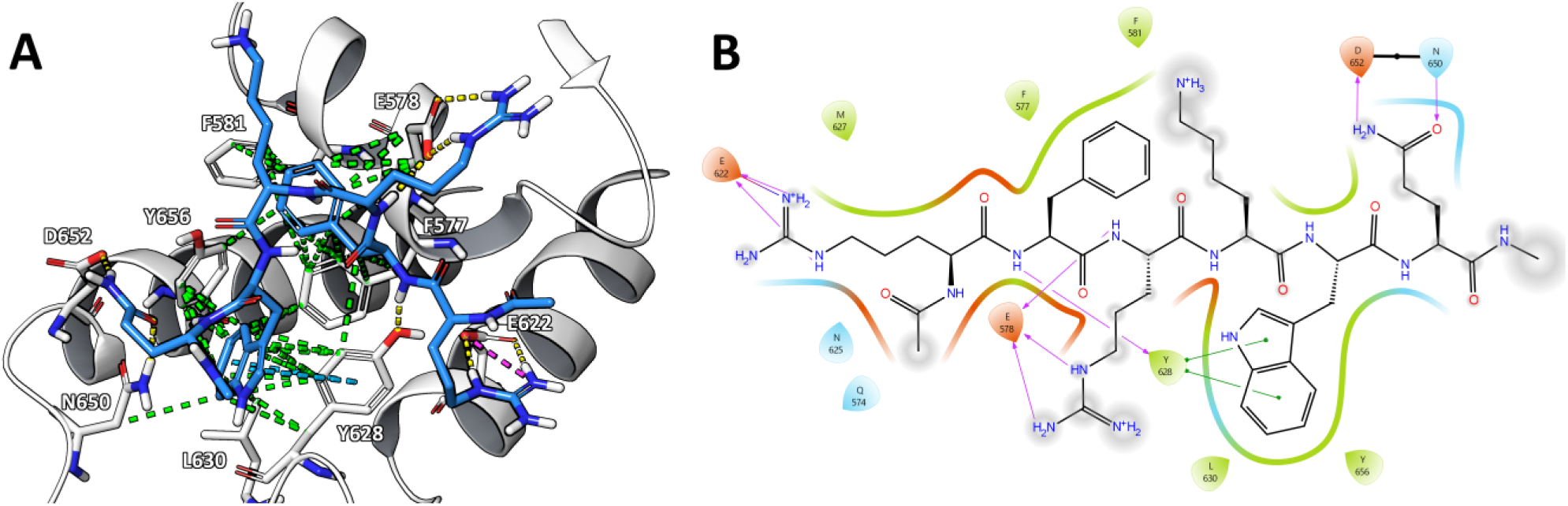
**(A)** The representative structure of the most populated cluster of **WI23-B** (with sequence Ace-RFRKWQ-NMe, light blue sticks) in complex with WWP1 (light grey cartoon and sticks), considering 1 µs of MD simulations. H-bond, salt bridges, π-π stacking and hydrophobic interactions are represented as yellow, purple, cyan and green dashed lines, respectively. **(B)** Ligand interaction diagram of **WI23-B** with WWP1 protein, applying a residue cutoff equal to 3.0 Å. H-bond interactions are highlighted by purple arrows, while salt bridges and π-π stacking interactions are indicated by blue/red and green lines, respectively. Atoms exposed to the solvent are highlighted by grey circles.

The analysis of **WI23-B**’s interactions with WWP1 (**Figure 10**) showed that the two additional residues (R_1_ and F_2_) in the hexapeptide created more interactions with WWP1’s residues than the tetrapeptide homolog **WI23** (**Figure 3A**). This likely accounts for the more favorable predicted ΔG value (**Table 4**) and the 5- to 10-fold lower K_d_ measured by SPR and MST assays, respectively (**Figure 4**). Interestingly, based on the **WI23-B** predicted binding mode with WWP1, the peptide’s N-terminus is oriented toward a polar area of the protein containing residues R573, Q574, and E603 (Supporting Information, **Figure S8**). This suggests that introducing additional residues at the N-terminus of the peptide could be a viable strategy to further enhance binding affinity. These new residues could form novel interactions (e.g., H-bonds) thereby increasing the overall complementarity with the target protein WWP1. At variance, the peptide C-terminus was fully solvent-exposed, indicating that the homologation at this portion would not be productive.

## 4. CONCLUSIONS

Targeting the ubiquitin-proteasome system has proven successful for the treatment of cancer, as exemplified by the clinically approved 26S proteasome inhibitors bortezomib^58^ and carfilzomib^59^, and inhibitors of ubiquitin-conjugating enzymes^60^. With nearly 600 distinct E3 ligases encoded in the human genome, these latter exhibit considerable structural and functional diversity, posing significant challenges for inhibition through conventional high-throughput screening approaches. Within the HECT E3 ligase family specifically, only a few small-molecule inhibitors have been mechanistically characterized^61,62^, including new covalent inhibitors which exhibited nanomolar potencym, and selectivity against NEDD4^63^. In this paper, we demonstrated the power of AI-driven computational strategies for the *de novo* design of peptide-based WWP1 inhibitors as potential therapeutic agents for BC. Starting from a rationally filtered peptide library, we combined classical computational methods (such as molecular docking and MD simulations) with ML or DL algorithms to efficiently predict and select candidate sequences with high affinity for WWP1. In particular, all eight tetrapeptides selected through three different approaches exhibited binding in the micromolar range, with **WI6** and **WI23** displaying clear K_d_ values of 6.4 µM (SPR) and 4.5 µM (MST), respectively. The sequences of these two peptides were subsequently optimized to enhance potential interactions with WWP1, yielding a series of hexapeptides. Among these, the **WI23** derivatives (**WI23-B** and **WI23-E**) demonstrated high-nanomolar affinities in both MST and SPR assays, with **WI23-B** showing the most unambiguous binding curves (K_d_ = 0.45 µM and 0.92 µM). **WI23-B** was therefore selected as the lead peptide for further TR-FRET enzymatic assays and biological studies in BC cell lines. Functional enzymatic assays confirmed WWP1 inhibition by **WI23-B**, displaying an IC₅₀ value in the low micromolar range (about 11 µM). Notably, **WI23-B** displayed the best cytotoxic effects in both BC cell lines (MCF7 and MDA-MB-231), even compared with free **WI23-E** or nanocapsule-loaded, especially when used in combination with the PI3K inhibitor BYL719, mirroring and in some cases surpassing the efficacy of I3C. Confocal microscopy in MCF-7 cells demonstrated that free **WI23-B** readily enters cells, while nanocarrier-encapsulated **WI23-B** achieves markedly higher intracellular accumulation. These results indicate that the developed nanocarrier efficiently delivers **WI23-B** into BC cells and constitutes a suitable system for intracellular peptide administration.

These findings confirm the WWP1/PTEN axis as a promising therapeutic target in BC and demonstrate the potential of peptide-based WWP1 inhibitors to enhance the activity of existing PI3K-targeted therapies. The identification of **WI23-B** as a lead compound provides a strong basis for further optimization, with the goal of generating a new class of peptide inhibitors capable of improving treatment outcomes while reducing toxicity. The structure of **WI23-B** can be used as a template for designing peptidomimetics, small molecules or peptide nucleic acids (PNAs) that replicate the key features responsible for binding to WWP1. To support this, structural studies are planned to validate the computational findings that led to the identification of **WI23-B** and to guide medicinal chemistry efforts aimed at discovering potent, selective, and bioavailable drug candidates. Taken together, the results showed in this paper open new avenues for the design of anticancer strategies that could improve survival and quality of life in difficult-to-treat BCs, including the TNBC subtype. Notably, the identification of novel WWP1 inhibitors aimed at fighting TNBC has the potential to transform the treatment paradigm for this aggressive subtype of cancer, offering promising prospects for the development of new anticancer protocols with reduced toxicity and improved life expectancy for patients.

## Data Availability Statement

All data is contained within the article and the Supporting Information.

## Acknowledgments

E.M.A.F would like to thank the Fondazione Umberto Veronesi for its precious support to his research activities. G.G. and E.M.A.F would like to thank INDACO and CINECA for providing high-performance computing resources and support. Authors would thank Arianna Baroni for her assistance in the computational analysis and preparation of His_6_-WWP1_HECT_.

