## Supporting Information for "AI-Driven Computational Design of Peptide-Based WWP1 Inhibitors as Promising Therapeutic Agents Against Breast Cancer, Including Triple-Negative Subtype"


**Table S1**. Binding free energy (ΔG) of all the 100 tetrapeptides subjected to MD simulation. In light green, are highlighted the 10 peptides with ΔG values ≤ –30.0 kcal/mol that were selected for extending their MD simulations to 500 ns. All the peptides were protected in the *N*- and *C*-termini by acetylation (Ace-) and amidation (-NMe), respectively.

| Peptide | Sequence | ΔG ± SEM[1] (0–250 ns) | Peptide | Sequence | ΔG ± SEM[1] (0–250 ns) |
|---|---|---|---|---|---|
| 001 | RKWQ | –27.5 ± 0.3 | 051 | SWYV | *unbound* |
| 002 | YYQW | –20.5 ± 0.2 | 052 | YYWC | –20.1 ± 0.2 |
| 003 | WQYE | *unbound* | 053 | QFYW | –24.2 ± 0.3 |
| 004 | NKKW | –15.4 ± 0.5 | 054 | RAYW | –19.9 ± 0.4 |
| 005 | YWQY | *unbound* | 055 | WHQT | *unbound* |
| 006 | WYYQ | –24.0 ± 0.3 | 056 | WHKH | –20.3 ± 0.2 |
| 007 | YKAW | –28.0 ± 0.2 | 057 | WFYW | *unbound* |
| 008 | QHYW | –17.6 ± 0.2 | 058 | WSQK | –17.5 ± 0.4 |
| 009 | WQKC | –26.4 ± 0.3 | 059 | YFQW | –18.2 ± 0.3 |
| 010 | KTYW | *unbound* | 060 | YKWR | –22.0 ± 0.5 |
| 011 (**WI1**) | TRYW | –32.0 ± 0.2 | 061 | FNKW | –22.2 ± 0.2 |
| 012 (**WI2**) | QYYW | –30.3 ± 0.1 | 062 | AWYC | *unbound* |
| 013 | WYYK | –22.0 ± 0.3 | 063 | QWYM | *unbound* |
| 014 (**WI3**) | TWYE | –37.7 ± 0.3 | 064 (**WI7**) | SCWQ | –30.5 ± 0.2 |
| 015 | WNYF | –21.0 ± 0.2 | 065 | NWYW | *unbound* |
| 016 | WYHN | *unbound* | 066 | KWYF | *unbound* |
| 017 | SQYW | *unbound* | 067 | YWYQ | –13.3 ± 0.3 |
| 018 | QWYA | –29.3 ± 0.4 | 068 | WYST | –20.2 ± 0.3 |
| 019 (**WI4**) | WQYL | –34.9 ± 0.3 | 069 | YYMW | *unbound* |
| 020 | RFYW | –8.3 ± 0.2 | 070 | NYYW | –25.2 ± 0.3 |

| | | | | | |
|---|---|---|---|---|---|
| 021 | WQYV | −17.6 ± 0.2 | 071 | WYKK | −21.3 ± 0.4 |
| 022 | WFRW | −26.3 ± 0.3 | 072 (**WI8**) | WQHR | −39.2 ± 0.4 |
| 023 | TKFW | −17.3 ± 0.2 | 073 | YWSF | −23.2 ± 0.3 |
| 024 | WYKQ | *unbound* | 074 | QWQY | −20.7 ± 0.5 |
| 025 | WYWQ | −17.7 ± 0.2 | 075 | QYWT | −23.2 ± 0.2 |
| 026 | QFQW | *unbound* | 076 | WTQY | −24.3 ± 0.4 |
| 027 | LYQW | −18.5 ± 0.3 | 077 | RWHY | −26.2 ± 0.3 |
| 028 | QWRS | *unbound* | 078 | WYRW | *unbound* |
| 029 | YFSW | −15.9 ± 0.2 | 079 | WFWH | −24.5 ± 0.2 |
| 030 | YWTK | *unbound* | 080 | KWTQ | *unbound* |
| 031 | MYYW | *unbound* | 081 | KRFW | −26.5 ± 0.3 |
| 032 | WYQQ | −17.8 ± 0.2 | 082 | NYHW | −15.2 ± 0.3 |
| 033 | WFKS | *unbound* | 083 | DYYW | −20.3 ± 0.2 |
| 034 | SYQW | −22.5 ± 0.2 | 084 | QYKW | *unbound* |
| 035 | NQWQ | *unbound* | 085 | TWYH | *unbound* |
| 036 | KYWP | *unbound* | 086 | RQWM | −23.7 ± 0.3 |
| 037 (**WI5**) | RRWE | −39.3 ± 0.3 | 087 | RYWS | −29.9 ± 0.3 |
| 038 | KFFW | *unbound* | 088 | QWKW | −22.6 ± 0.3 |
| 039 | KYNW | −19.9 ± 0.3 | 089 | WFKN | −17.6 ± 0.3 |
| 040 | WYRN | −23.9 ± 0.2 | 090 | WQYQ | *unbound* |
| 041 | WNYN | −27.7 ± 0.3 | 091 | WQYT | *unbound* |
| 042 | VNYW | *unbound* | 092 | EFWQ | *unbound* |
| 043 (**WI6**) | SWYT | −40.3 ± 0.2 | 093 | KMYW | −23.3 ± 0.3 |
| 044 | QHWH | −19.4 ± 0.1 | 094 | QNTW | −22.9 ± 0.4 |
| 045 | SNYW | −21.5 ± 0.2 | 095 | EWYQ | *unbound* |
| 046 | QWHY | −24.5 ± 0.2 | 096 | WYKH | *unbound* |
| 047 | FWYM | −23.5 ± 0.2 | 097 | WYNT | −22.5 ± 0.2 |
| 048 | WFKH | *unbound* | 098 (**WI9**) | NWRW | −32.4 ± 0.4 |
| 049 | WYYC | *unbound* | 099 (**WI10**) | IHYW | −38.4 ± 0.2 |
| 050 | MNYW | *unbound* | 100 | AWYM | −21.2 ± 0.2 |

[1] kcal/mol; SEM = Standard Error of Mean.

**Table S2**. Results of the cluster analysis carried out considering different cut-off levels. The values of the three main populated cluster groups are expressed in percentage (%). In bold is highlighted the chosen cut-off level.

| **WI6** | | | | | **WI23** | | | | | **WI23-B** | | | | |
|---|---|---|---|---|---|---|---|---|---|---|---|---|---|---|
| **cut-off** | **N°** | **1°** | **2°** | **3°** | **cut-off** | **N°** | **1°** | **2°** | **3°** | **cut-off** | **N°** | **1°** | **2°** | **3°** |
| 0.4 Å | 203 | 44.3 | 16.6 | 3.0 | 1.0 Å | 15 | 57.7 | 30.0 | 4.4 | 1.4 Å | 7 | 54.5 | 39.8 | 2.4 |
| 0.6 Å | 52 | 63.1 | 11.0 | 5.9 | 1.2 Å | 8 | 60.3 | 35.3 | 1.7 | 1.6 Å | 5 | 62.4 | 33.6 | 3.8 |
| 0.8 Å | 20 | 77.9 | 9.8 | 4.2 | 1.4 Å | 6 | 62.9 | 36.1 | 0.7 | **1.8 Å** | **4** | **73.9** | **22.2** | **3.9** |
| **1.0 Å** | **8** | **90.2** | **4.1** | **3.4** | **1.6 Å** | **3** | **74.3** | **21.5** | **4.1** | 2.0 Å | 3 | 90.4 | 7.3 | 2.3 |
| 1.2 Å | 5 | 96.2 | 3.2 | 0.4 | 1.8 Å | 3 | 87.6 | 12.1 | 0.2 | 2.2 Å | 3 | 98.1 | 1.0 | 0.9 |
| 1.4 Å | 3 | 98.6 | 1.3 | 0.1 | 2.0 Å | 3 | 97.8 | 1.8 | 0.4 | 2.4 Å | 3 | 99.8 | 0.1 | 0.1 |

N° = Number of clusters.

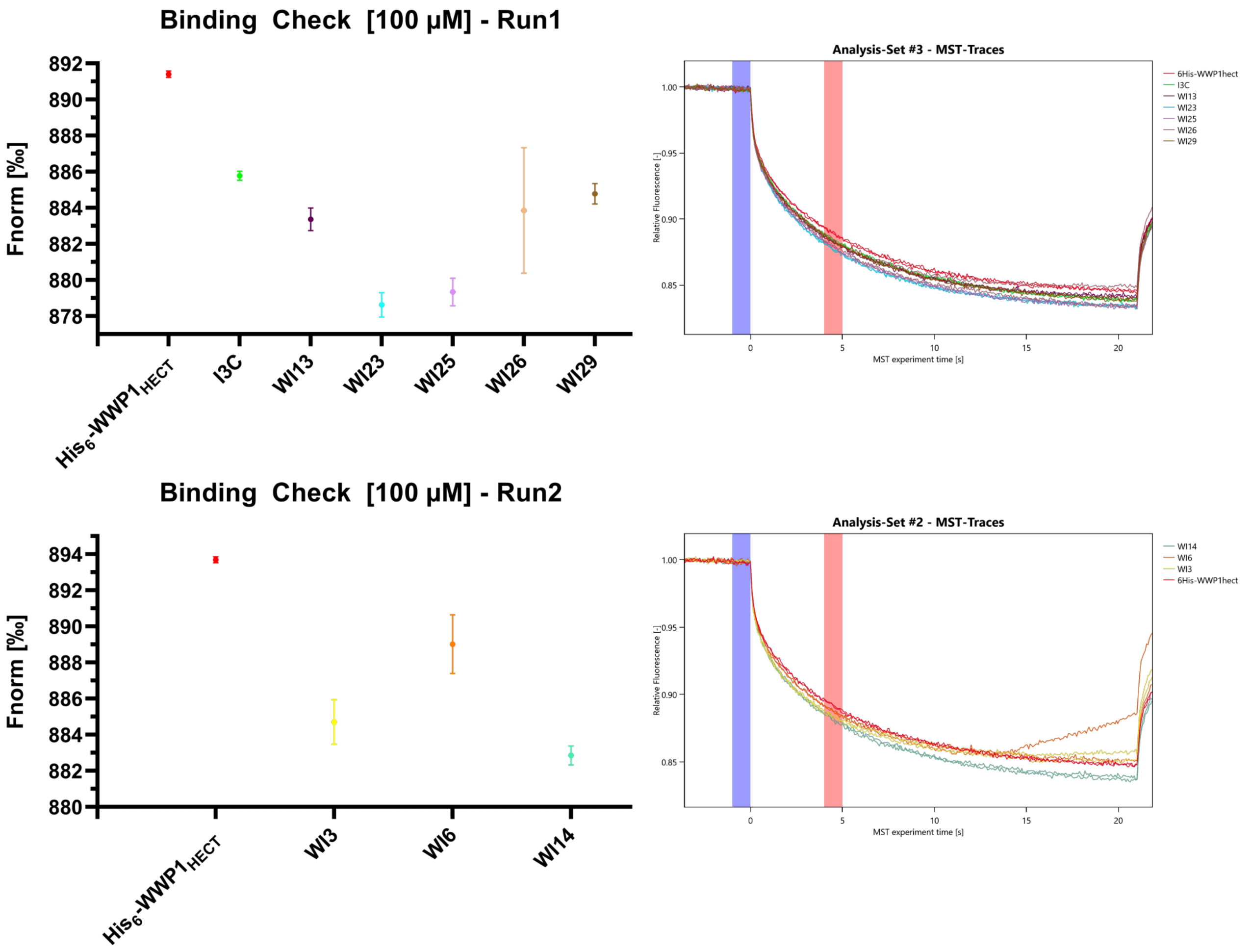


**Figure S1**. MST binding check experiments (left) with corresponding MST traces (right). All ligands were tested at a concentration of 100 µM. Since the analysis focused on the time point after 5 s, a difference in Fnorm of at least 2.5 is required to indicate binding.

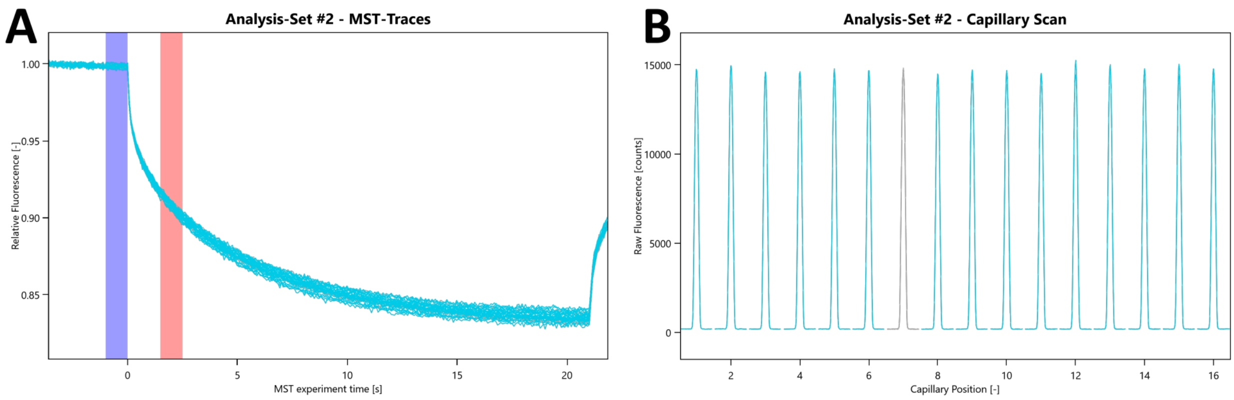


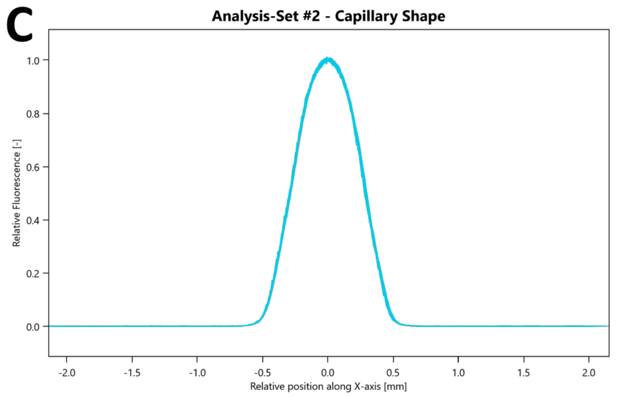


D

| | |
|---|---|
| Target Name: | His6-WWP1hect |
| Target Concentration: | 10 nM |
| Ligand Name: | WI23 |
| Ligand Concentration: | 156 µM to 0.00477 µM |
| n: | 2 |
| Comments: | |
| Excitation Power: | 20% |
| MST Power: | 40% |
| Temperature: | 25.0°C |
| Kd: | 4.5057E-06 |
| Kd Confidence: | ± 9.8953E-07 |
| Response Amplitude: | 3.6097733 |
| TargetConc: | 1E-08[Fixed] |
| Unbound: | 910.31 |
| Bound: | 906.7 |
| Std. Error of Regression: | 0.22063416 |
| Reduced $\chi^2$: | 2.749866 |
| Signal to Noise: | 17.671791 |

**Figure S2**. MST binding affinity assay report of **WI23** on His$_6$-WWP1$_{HECT}$. **(A)** MST traces, **(B)** capillary scan, **(C)** capillary shape, and **(D)** dataset overview. Concentration points highlighted in grey were excluded from the analysis as they were identified as outliers.

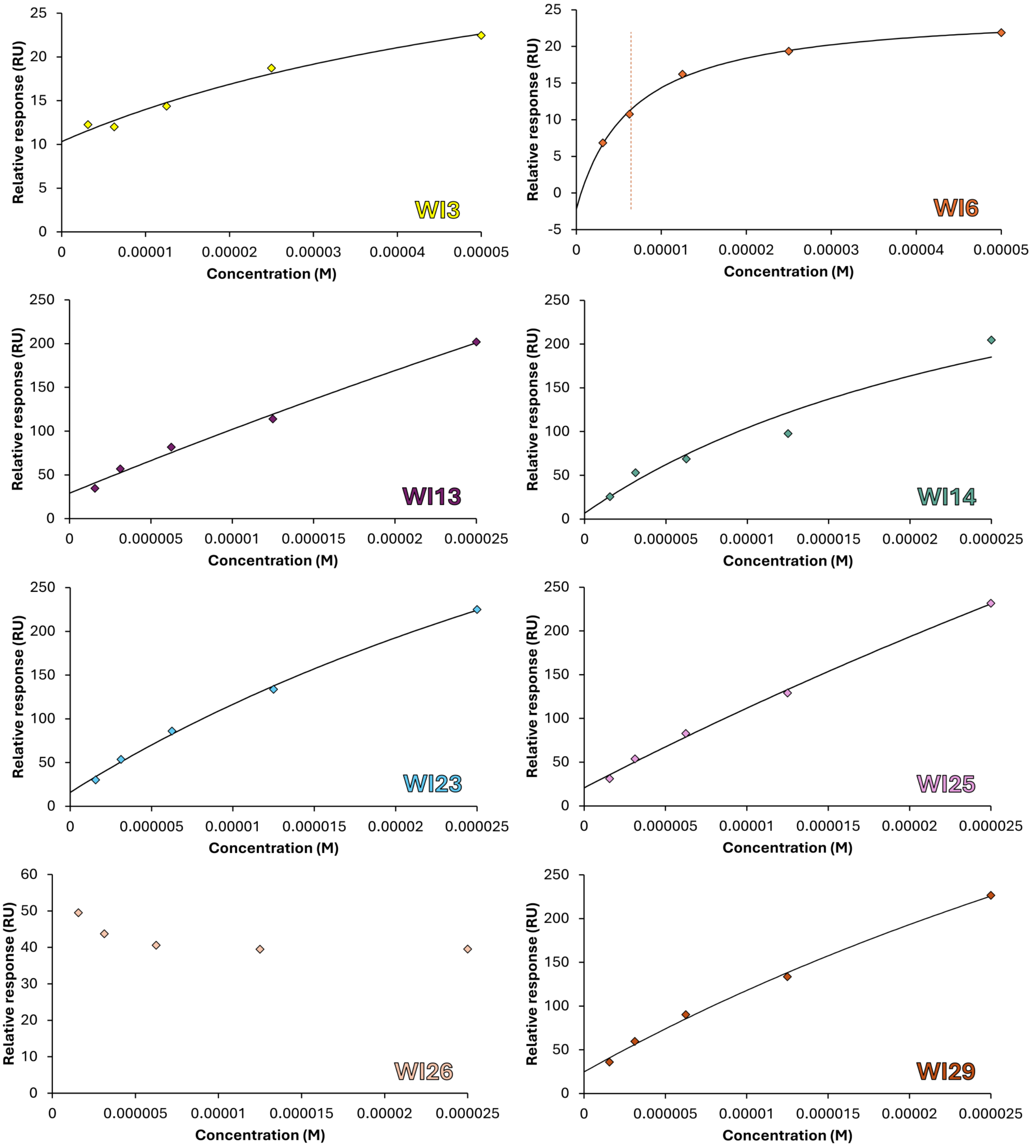


**Figure S3**. SPR experiments. All the tetrapeptides were injected in a single kinetic experiment at five concentration points (1.56, 3.12, 6.25, 12.5, and 25 µM), over the immobilized His$_6$-WWP1$_{HECT}$ protein at a flow rate of 30 µL/min. In the case of **WI3** and **WI6**, they were tested from 3.12 to 50 µM doubling the concentration at each point.

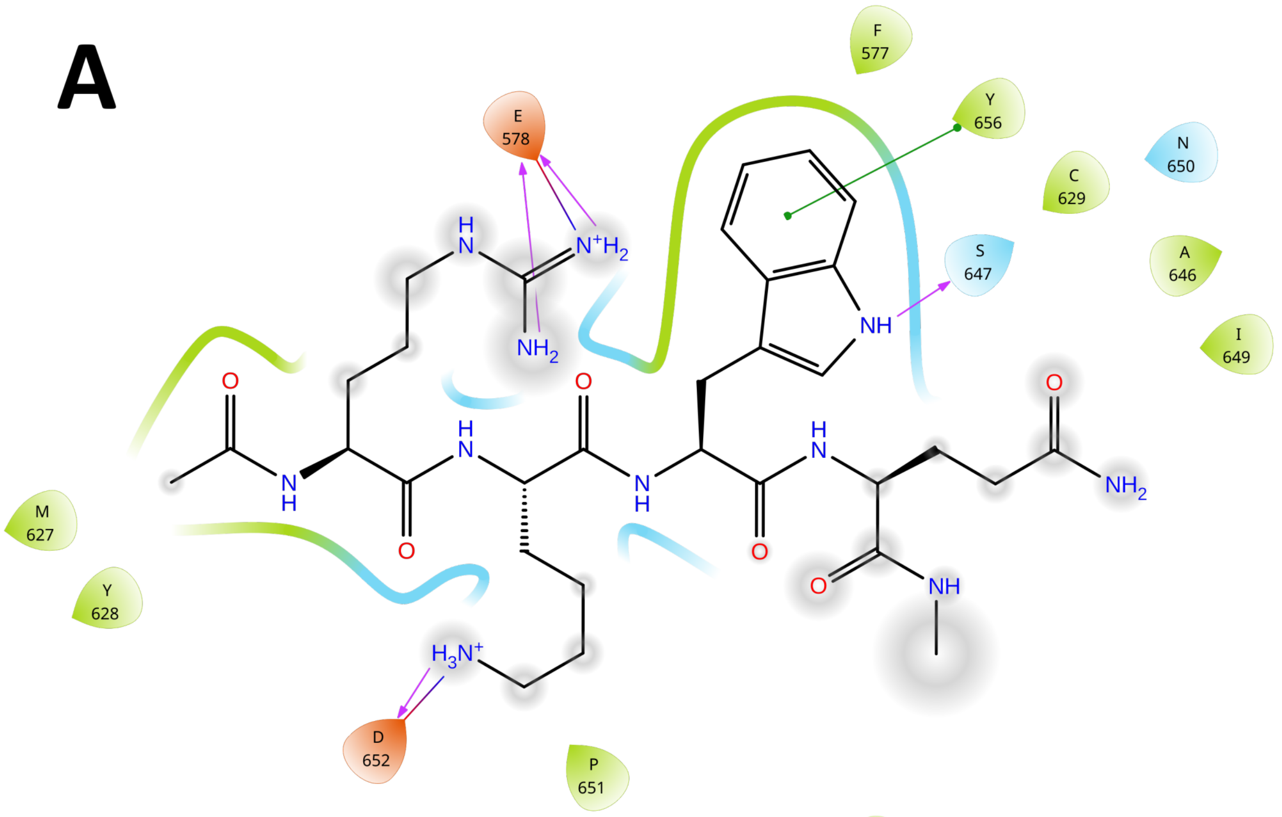


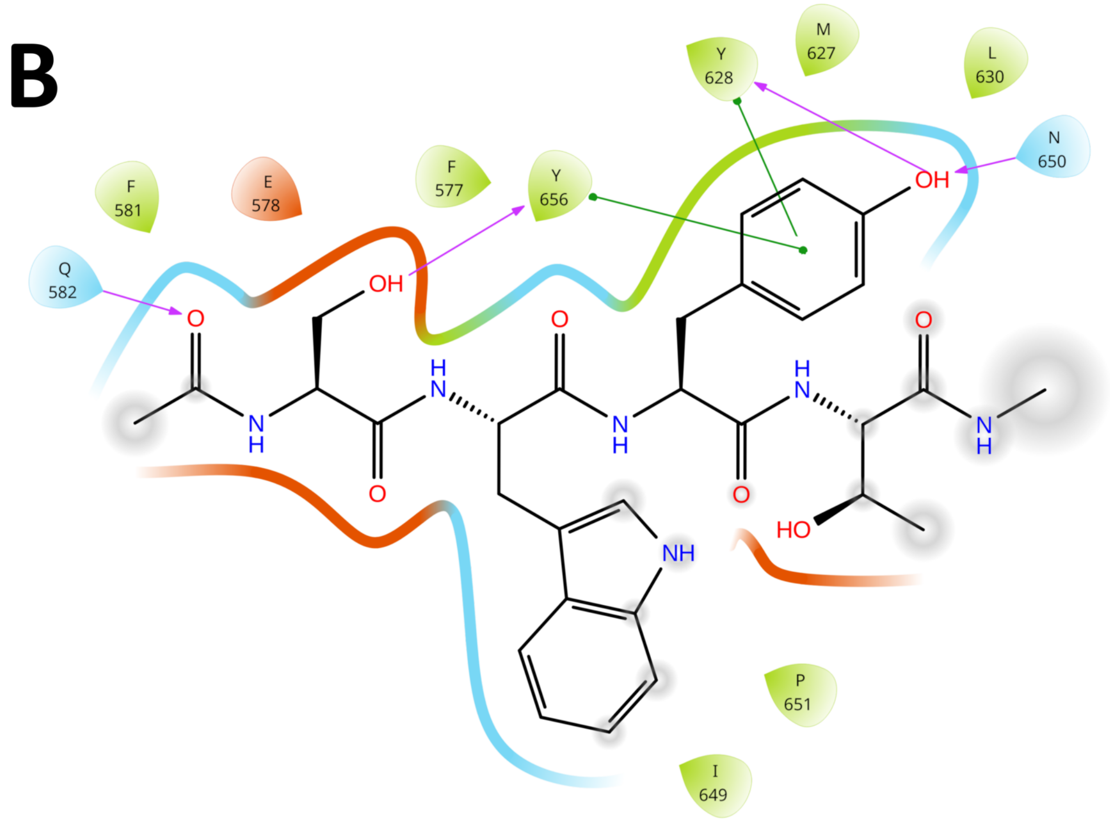


**Figure S4**. Ligand interaction diagram of **(A) WI23** and **(B) WI6** with WWP1 protein, applying a residue cutoff equal to 3.0 Å. H-bond interactions are highlighted by purple arrows, while salt bridges and π-cation interactions are indicated by blue/red and green lines, respectively. Atoms exposed to the solvent are highlighted by grey circles.

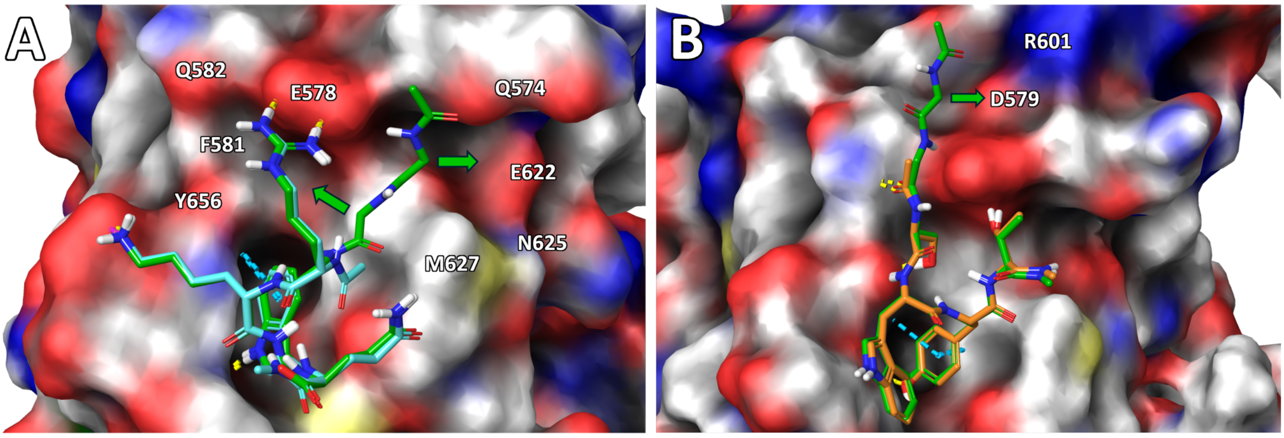


**Figure S5**. Hexapeptide analogues (green sticks) of **(A) WI23** (cyan sticks) and **(B) WI6** (orange sticks). These analogues are characterized by two additional GG residues at the N-terminus and were used as starting points for the affinity maturation protocol, with the aim of accessing additional residues that were previously inaccessible to the tetrapeptides.

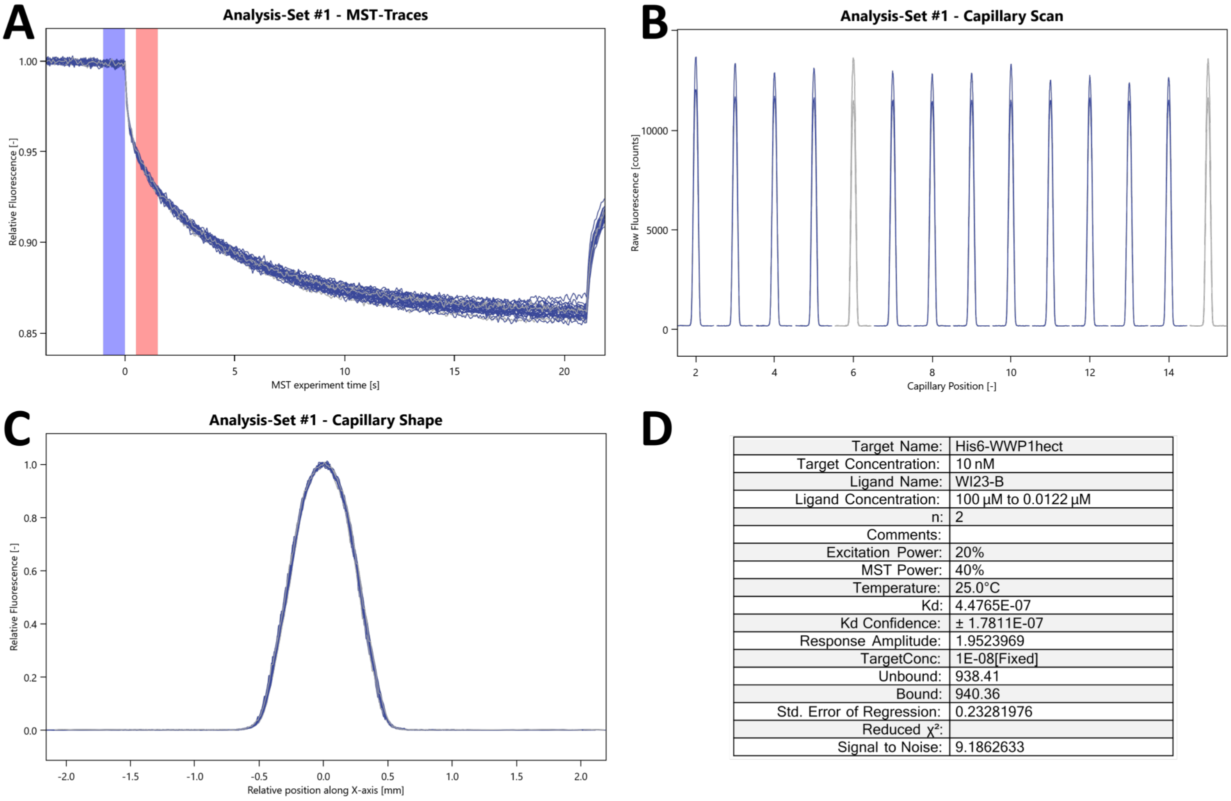


| | |
|---|---|
| Target Name: | His6-WWP1hect |
| Target Concentration: | 10 nM |
| Ligand Name: | WI23-B |
| Ligand Concentration: | 100 µM to 0.0122 µM |
| n: | 2 |
| Comments: | |
| Excitation Power: | 20% |
| MST Power: | 40% |
| Temperature: | 25.0°C |
| Kd: | 4.4765E-07 |
| Kd Confidence: | ± 1.7811E-07 |
| Response Amplitude: | 1.9523969 |
| TargetConc: | 1E-08[Fixed] |
| Unbound: | 938.41 |
| Bound: | 940.36 |
| Std. Error of Regression: | 0.23281976 |
| Reduced $\chi^2$: | |
| Signal to Noise: | 9.1862633 |

**Figure S6**. MST binding affinity assay report of **WI23-B** on His$_6$-WWP1$_{HECT}$. **(A)** MST traces, **(B)** capillary scan, **(C)** capillary shape, and **(D)** dataset overview. Concentration points highlighted in grey were excluded from the analysis as they were identified as outliers.

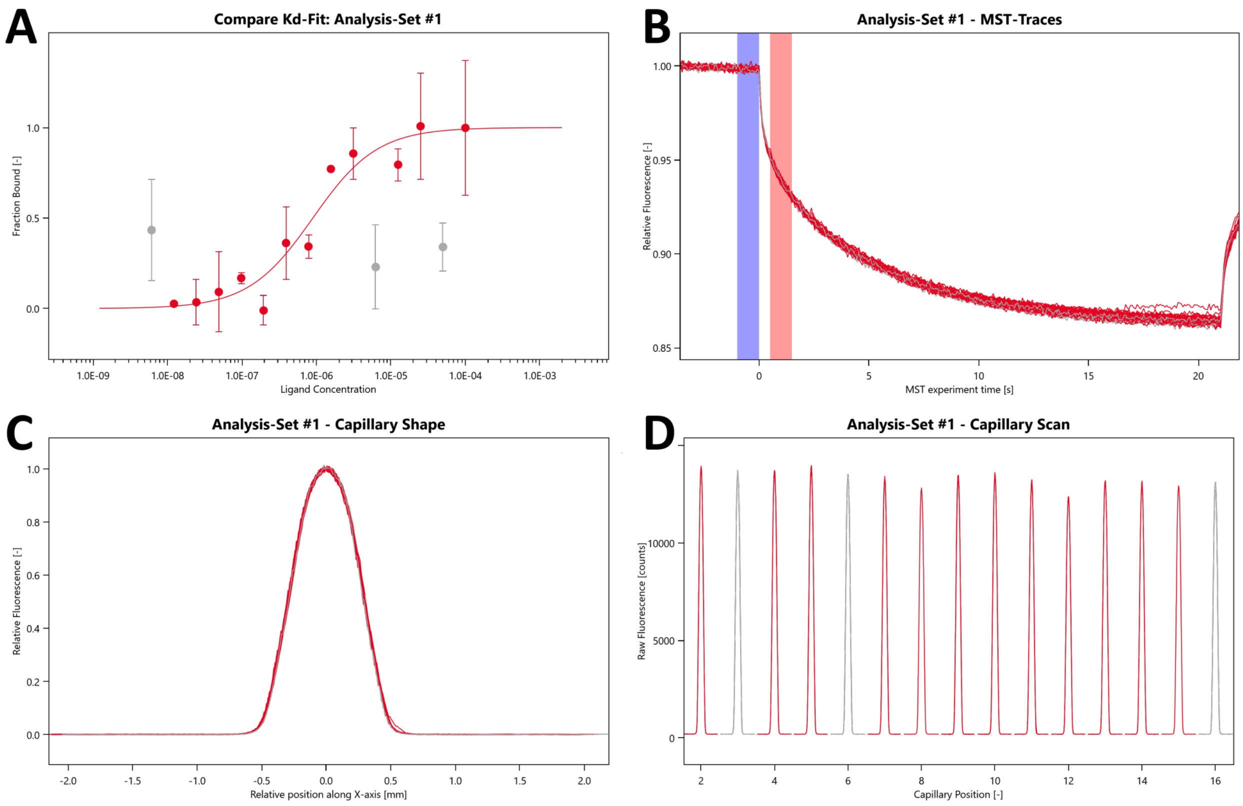


**E**

| | |
|---|---|
| Target Name: | His6-WWP1hect |
| Target Concentration: | 10 nM |
| Ligand Name: | WI23-E |
| Ligand Concentration: | 100 µM to 0.0122 µM |
| n: | 2 |
| Comments: | |
| Excitation Power: | 20% |
| MST Power: | 40% |
| Temperature: | 25.0°C |
| Kd: | 8.8899E-07 |
| Kd Confidence: | ± 3.2622E-07 |
| Response Amplitude: | 2.1062902 |
| TargetConc: | 1E-08[Fixed] |
| Unbound: | 940.16 |
| Bound: | 942.27 |
| Std. Error of Regression: | 0.22933466 |
| Reduced $\chi^2$: | |
| Signal to Noise: | 10.153683 |

**Figure S7**. MST binding affinity assay report of **WI23-E** on His$_6$-WWP1$_{HECT}$. **(A)** K$_d$ curve obtained by two independent experiments, **(B)** Binding MST traces, **(C)** capillary shape, **(D)** capillary scan, and **(E)** dataset overview. Concentration points highlighted in grey were excluded from the analysis as they were identified as outliers.

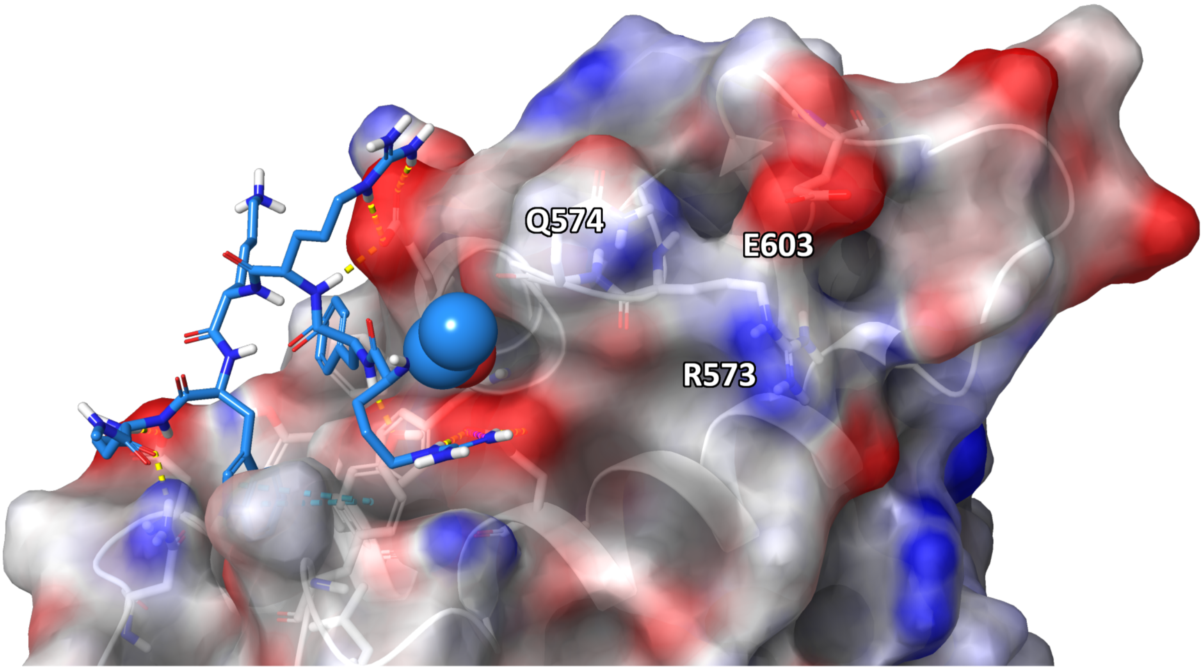


**Figure S8**. Representative structure of the most populated cluster of **WI23-B** (light blue sticks) in complex with WWP1. The N-terminus of **WI23-B** is shown as a van der Waals surface. The solvent-accessible surface of the protein is colored according to the electrostatic potential of the atoms: negatively charged regions are depicted in red, while positively charged regions are shown in blue.

#### HPLC chromatograms

Correspondence between the names of the peptides during synthesis with those reported in the manuscript.

| Manufacturer | Synthesis Name | Manuscript Name |
|---|---|---|
| Proteogenix | New_pep2 TE-4 | **WI3** |
| Proteogenix | New_pep1 ST-4 | **WI6** |
| Proteogenix | Enfa6 AC-WY-4 | **WI13** |
| Proteogenix | Enfa7 AC-QY-4 | **WI14** |
| Proteogenix | Enfa1 AC-RQ-4 | **WI23** |
| Proteogenix | Enfa2 AC-YQ-4 | **WI25** |
| Proteogenix | Enfa3 AC-FC-4 | **WI26** |
| Proteogenix | Enfa4 AC-YH-4 | **WI29** |
| *in-house* | WWP3 | **WI6-B** |
| Genscript | ENFA1-A | **WI23-B** |
| Genscript | ENFA1-B | **WI23-E** |
| *in-house* | CF-[WI23-B] | **CF-[WI23-B]** |

| | |
|---|---|
| Sample: | New_pep2 TE-4 |
| Lot. No.: | P201104-HS843563 |
| Column: | Gemini-NX 5µ C18 110A, 4.6*250mm |
| Solvent A | A: 0.1% Trifluoroacetic Acid in 100% Acetonitrile |
| Solvent B | B: 0.1% Trifluoroacetic Acid in 100% Water |

Gradient:

| | A | B |
|---|---|---|
| 0.0min | 25% | 75% |
| 25.0min | 50% | 50% |
| 25.1min | 100% | 0% |
| 30.0min | Stop | |

| | |
|---|---|
| Volume: | 10µl |
| Wavelength: | 220nm |
| Flow rate: | 1.0ml/min |

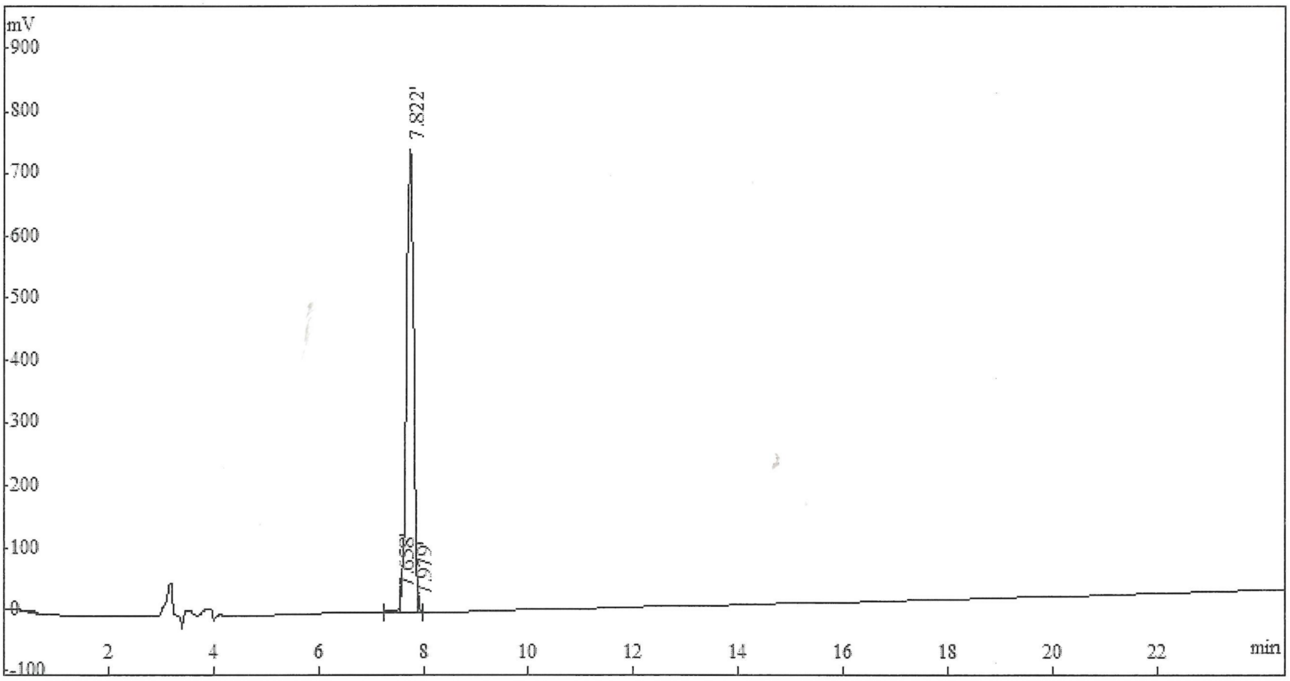


| Rank | Time | Conc. | Area | Height |
|---|---|---|---|---|
| 1 | 7.658 | 1.6164 | 104125 | 29624 |
| 2 | 7.822 | 97.9083 | 6307213 | 744664 |
| 3 | 7.979 | 0.4753 | 30620 | 19976 |
| Total | | 100 | 6441958 | 794264 |

Sample: New_pep1 ST-4
Lot. No.: P210518-CQ898185
Column: Kromasil 100-5C18, 4.6*250mm, 5μm
Solvent A A: 0.1% Trifluoroacetic Acid in 100% Acetonitrile
Solvent B B: 0.1% Trifluoroacetic Acid in 100% Water
Gradient:

| | A | B |
|---|---|---|
| 0.0min | 21% | 79% |
| 25.0min | 46% | 54% |
| 25.1min | 100% | 0% |
| 30.0min | Stop | |

Volume: 10μl
Wavelength: 220nm
Flow rate: 1.0ml/min

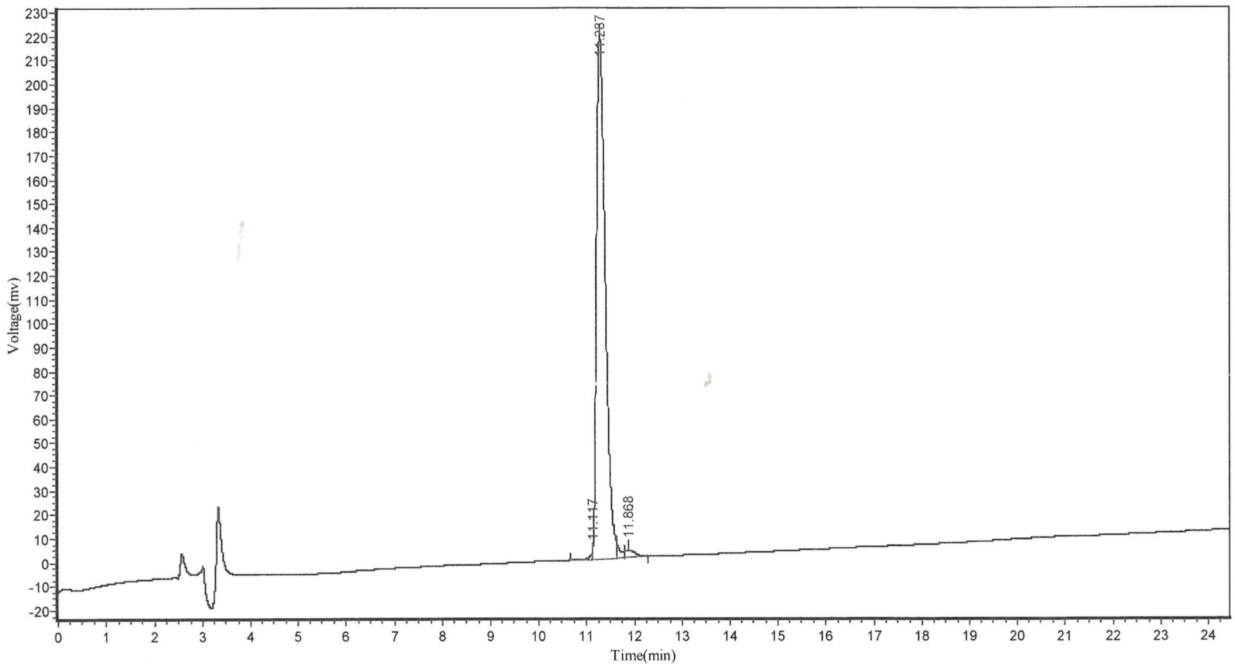


| Peak | Time | Height | Area | Conc. |
|---|---|---|---|---|
| 1 | 11.117 | 2514.728 | 7566.855 | 0.2578 |
| 2 | 11.287 | 218978.969 | 2851578.750 | 97.1539 |
| 3 | 11.287 | 6758.241 | 32701.406 | 1.1141 |
| 4 | 11.868 | 3009.975 | 43266.672 | 1.4741 |
| Total | | | | 100.0000 |

| | |
|---|---|
| Sample: | Enfa6 AC-WY-4 |
| Lot. No.: | P210305-CQ874402 |
| Column: | Kromasil 100-5C18, 4.6*250mm, 5μm |
| Solvent A | A: 0.1% Trifluoroacetic Acid in 100% Acetonitrile |
| Solvent B | B: 0.1% Trifluoroacetic Acid in 100% Water |

Gradient:

| | A | B |
|---|---|---|
| 0.0min | 22% | 78% |
| 25.0min | 47% | 53% |
| 25.1min | 100% | 0% |
| 30.0min | Stop | |

| | |
|---|---|
| Volume: | 10μl |
| Wavelength: | 220nm |
| Flow rate: | 1.0ml/min |

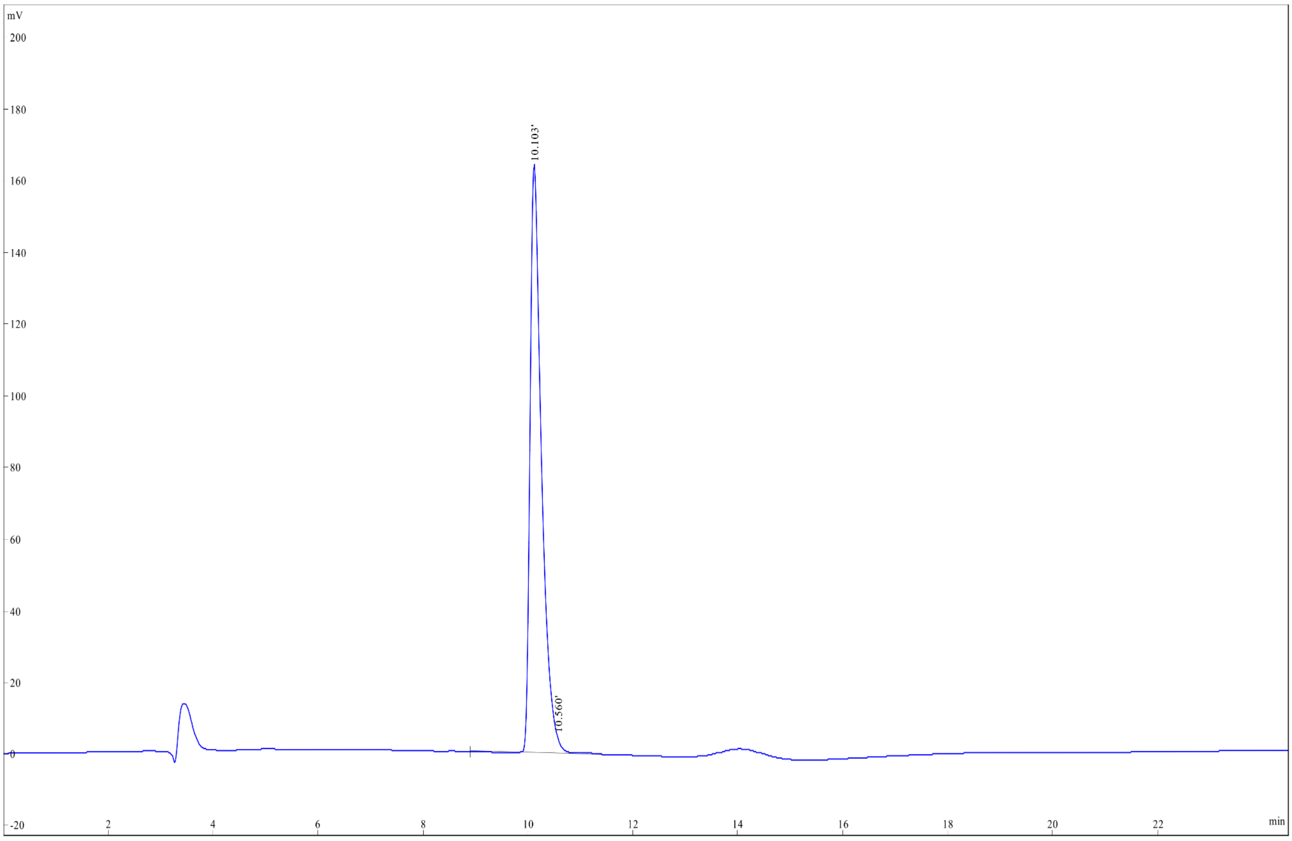


| Rank | Time | Conc. | Area | Height |
|---|---|---|---|---|
| 1 | 10.103 | 97.88 | 2456663 | 164667 |
| 2 | 10.560 | 2.113 | 53039 | 4222 |
| Total | | 100 | 2509702 | 168889 |

| | | | |
|---|---|---|---|
| Sample: | Enfa7 AC-QY-4 | | |
| Lot. No.: | P210305-CQ874403 | | |
| Column: | Kromasil 100-5C18, 4.6*250mm, 5μm | | |
| Solvent A | A: 0.1% Trifluoroacetic Acid in 100% Acetonitrile | | |
| Solvent B | B: 0.1% Trifluoroacetic Acid in 100% Water | | |
| Gradient: | | A | B |
| | 0.0min | 21% | 79% |
| | 25.0min | 46% | 54% |
| | 25.1min | 100% | 0% |
| | 30.0min | Stop | |
| Volume: | 10μl | | |
| Wavelength: | 220nm | | |
| Flow rate: | 1.0ml/min | | |

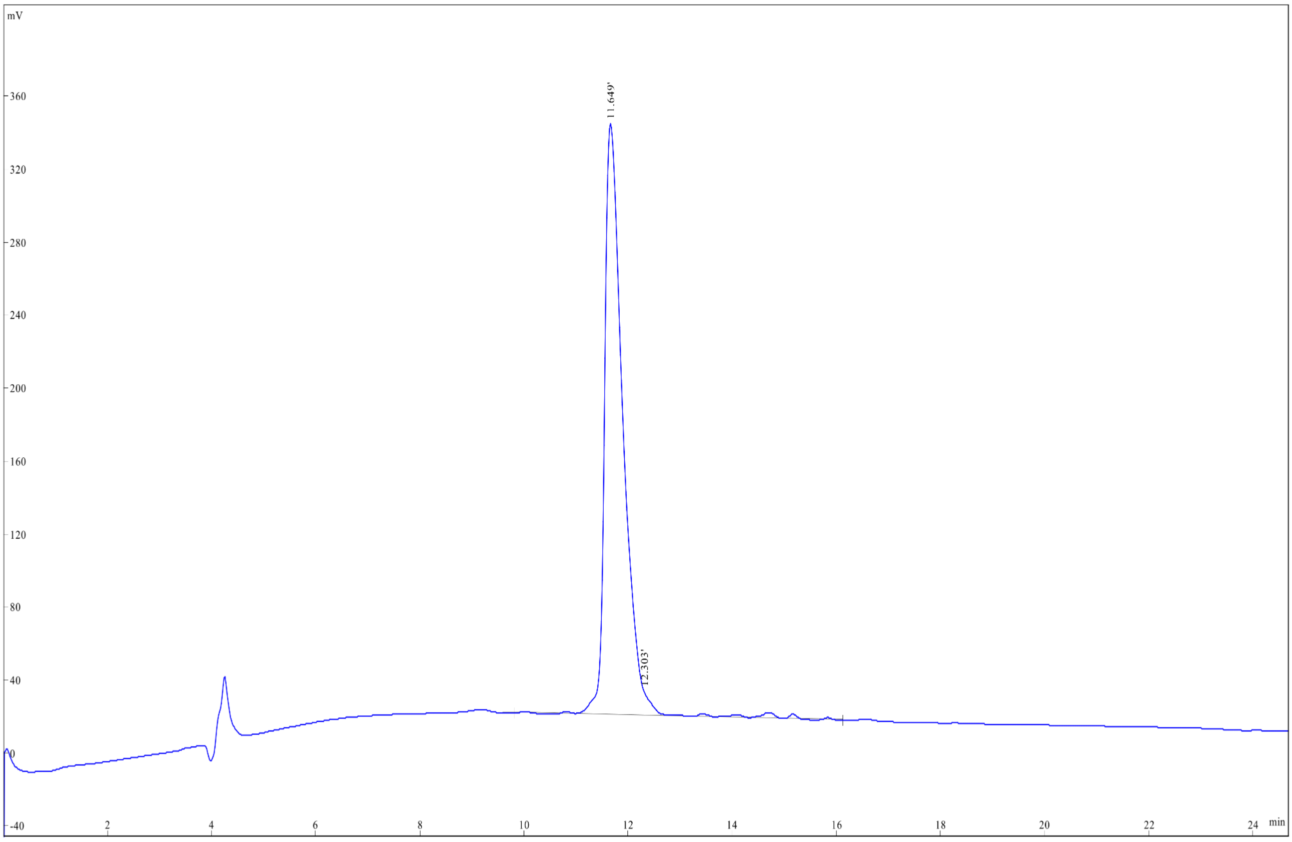


| Rank | Time | Conc. | Area | Height |
|---|---|---|---|---|
| 1 | 11.649 | 95.78 | 7864958 | 324479 |
| 2 | 12.303 | 4.214 | 346055 | 12903 |
| Total | | 100 | 8211013 | 337382 |

| | | | |
|---|---|---|---|
| Sample: | Enfa 1 AC-RQ-4 | | |
| Lot. No.: | P210305-CQ874397 | | |
| Column: | Kromasil 100-5C18, 4.6*250mm, 5μm | | |
| Solvent A | A: 0.1% Trifluoroacetic Acid in 100% Acetonitrile | | |
| Solvent B | B: 0.1% Trifluoroacetic Acid in 100% Water | | |
| Gradient: | | A | B |
| | 0.0min | 25% | 75% |
| | 25.0min | 50% | 50% |
| | 25.1min | 100% | 0% |
| | 30.0min | Stop | |
| Volume: | 10μl | | |
| Wavelength: | 220nm | | |
| Flow rate: | 1.0ml/min | | |

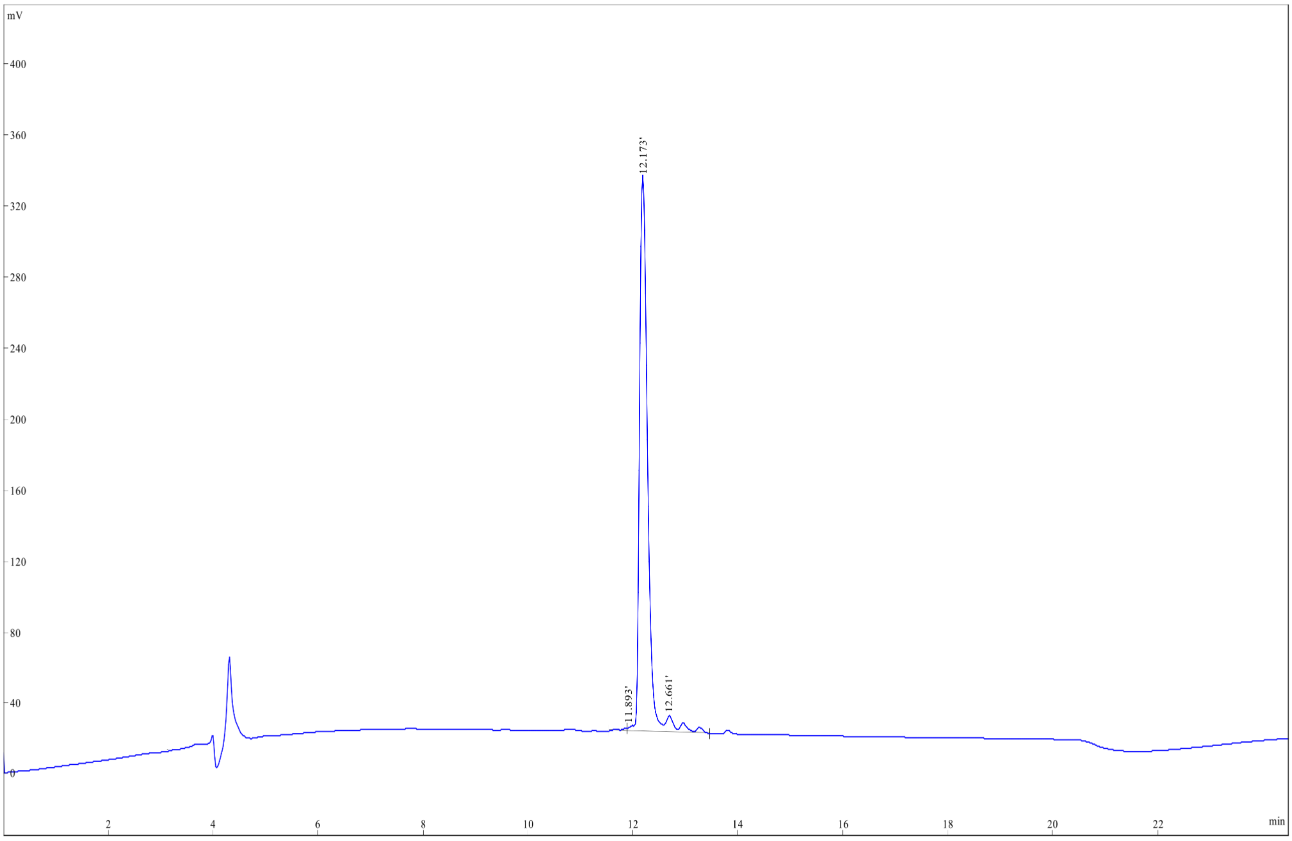


| Rank | Time | Conc. | Area | Height |
|---|---|---|---|---|
| 1 | 11.893 | 0.533 | 18447 | 1807 |
| 2 | 12.173 | 95.01 | 3288069 | 313931 |
| 3 | 12.661 | 4.451 | 154019 | 8836 |
| Total | | 100 | 3460535 | 324574 |

| | | | |
|---|---|---|---|
| Sample: | Enfa2 AC-YQ-4 | | |
| Lot. No.: | P210305-CQ874398 | | |
| Column: | Kromasil 100-5C18, 4.6*250mm, 5μm | | |
| Solvent A | A: 0.1% Trifluoroacetic Acid in 100% Acetonitrile | | |
| Solvent B | B: 0.1% Trifluoroacetic Acid in 100% Water | | |
| Gradient: | | A | B |
| | 0.0min | 22% | 78% |
| | 25.0min | 47% | 53% |
| | 25.1min | 100% | 0% |
| | 30.0min | Stop | |
| Volume: | 10μl | | |
| Wavelength: | 220nm | | |
| Flow rate: | 1.0ml/min | | |

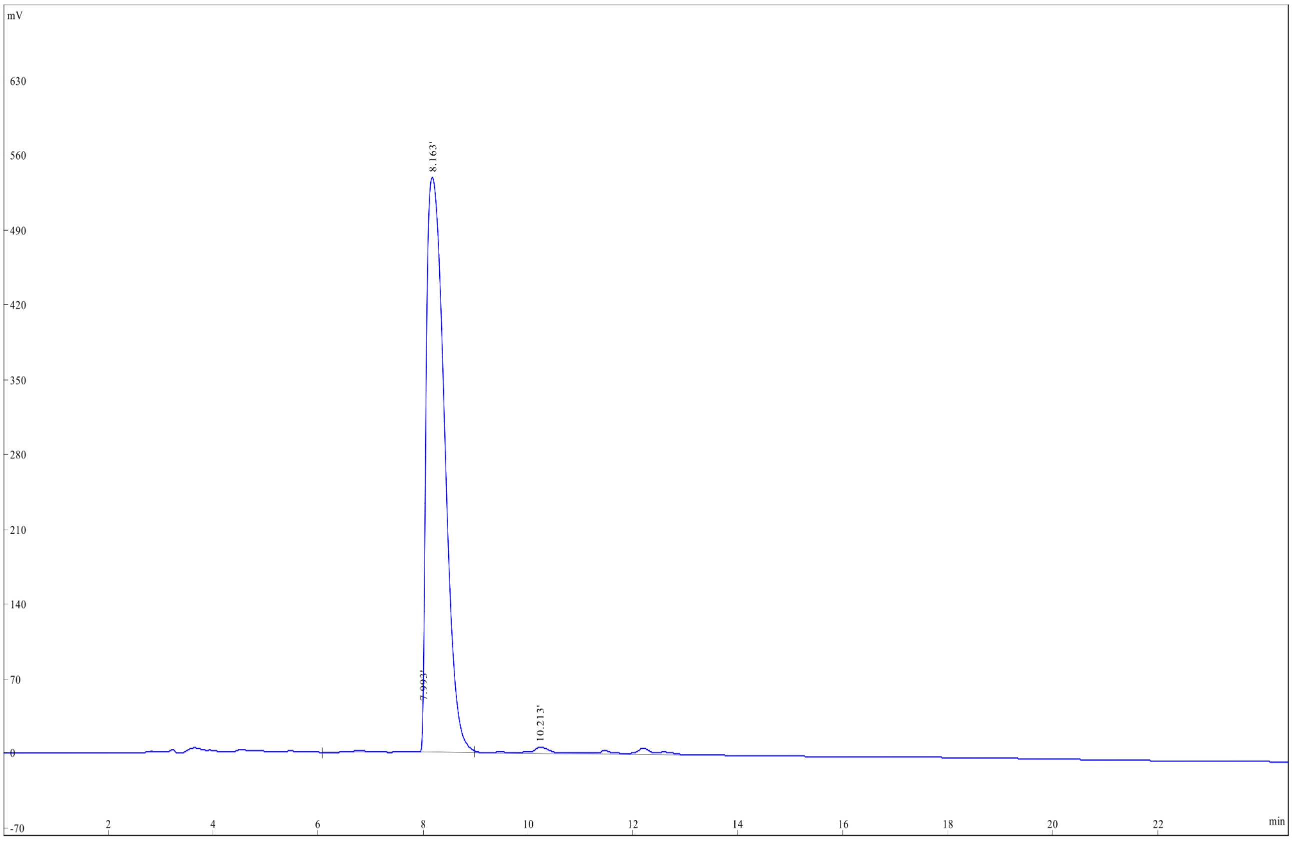


| Rank | Time | Conc. | Area | Height |
|---|---|---|---|---|
| 1 | 7.993 | 1.228 | 166807 | 69010 |
| 2 | 8.163 | 95.13 | 12925816 | 539101 |
| 3 | 10.213 | 3.643 | 494997 | 6824 |
| Total | | 100 | 13587620 | 614935 |

| | | | |
|---|---|---|---|
| Sample: | Enfa3 AC-FC-4 | | |
| Lot. No.: | P210305-CQ874399 | | |
| Column: | Kromasil 100-5C18, 4.6*250mm, 5μm | | |
| Solvent A | A: 0.1% Trifluoroacetic Acid in 100% Acetonitrile | | |
| Solvent B | B: 0.1% Trifluoroacetic Acid in 100% Water | | |
| Gradient: | | A | B |
| | 0.0min | 35% | 65% |
| | 25.0min | 60% | 40% |
| | 25.1min | 100% | 0% |
| | 30.0min | Stop | |
| Volume: | 10μl | | |
| Wavelength: | 220nm | | |
| Flow rate: | 1.0ml/min | | |

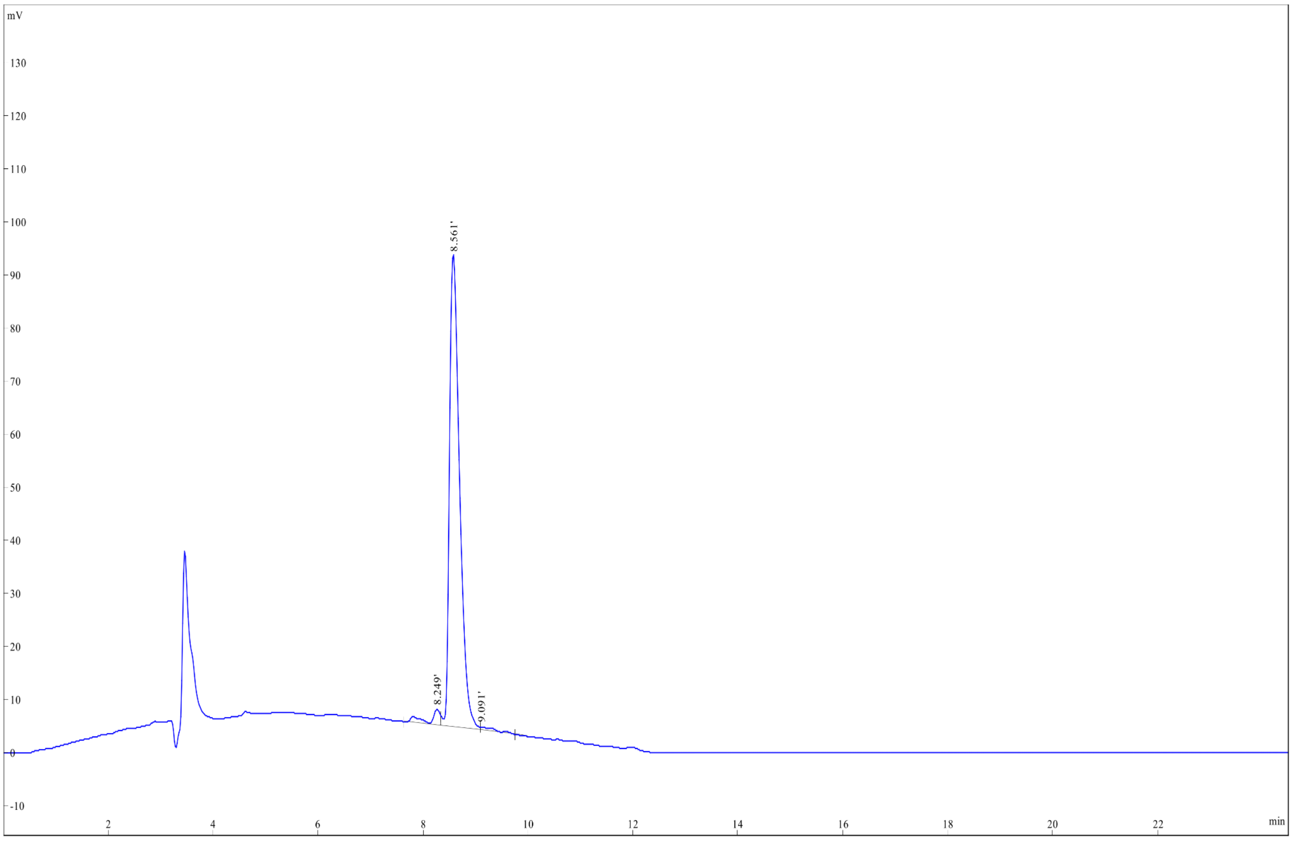


| Rank | Time | Conc. | Area | Height |
|---|---|---|---|---|
| 1 | 8.249 | 3.415 | 42976 | 2985 |
| 2 | 8.561 | 95.22 | 1198470 | 89213 |
| 3 | 9.091 | 1.364 | 17167 | 664 |
| Total | | 100 | 1258613 | 92862 |

Sample: Enfa4 AC-YH-4
Lot. No.: P210305-CQ874400
Column: Kromasil 100-5C18, 4.6*250mm, 5μm
Solvent A A: 0.1% Trifluoroacetic Acid in 100% Acetonitrile
Solvent B B: 0.1% Trifluoroacetic Acid in 100% Water
Gradient:

| | A | B |
|---|---|---|
| 0.0min | 20% | 80% |
| 25.0min | 45% | 55% |
| 25.1min | 100% | 0% |
| 30.0min | Stop | |

Volume: 10μl
Wavelength: 220nm
Flow rate: 1.0ml/min

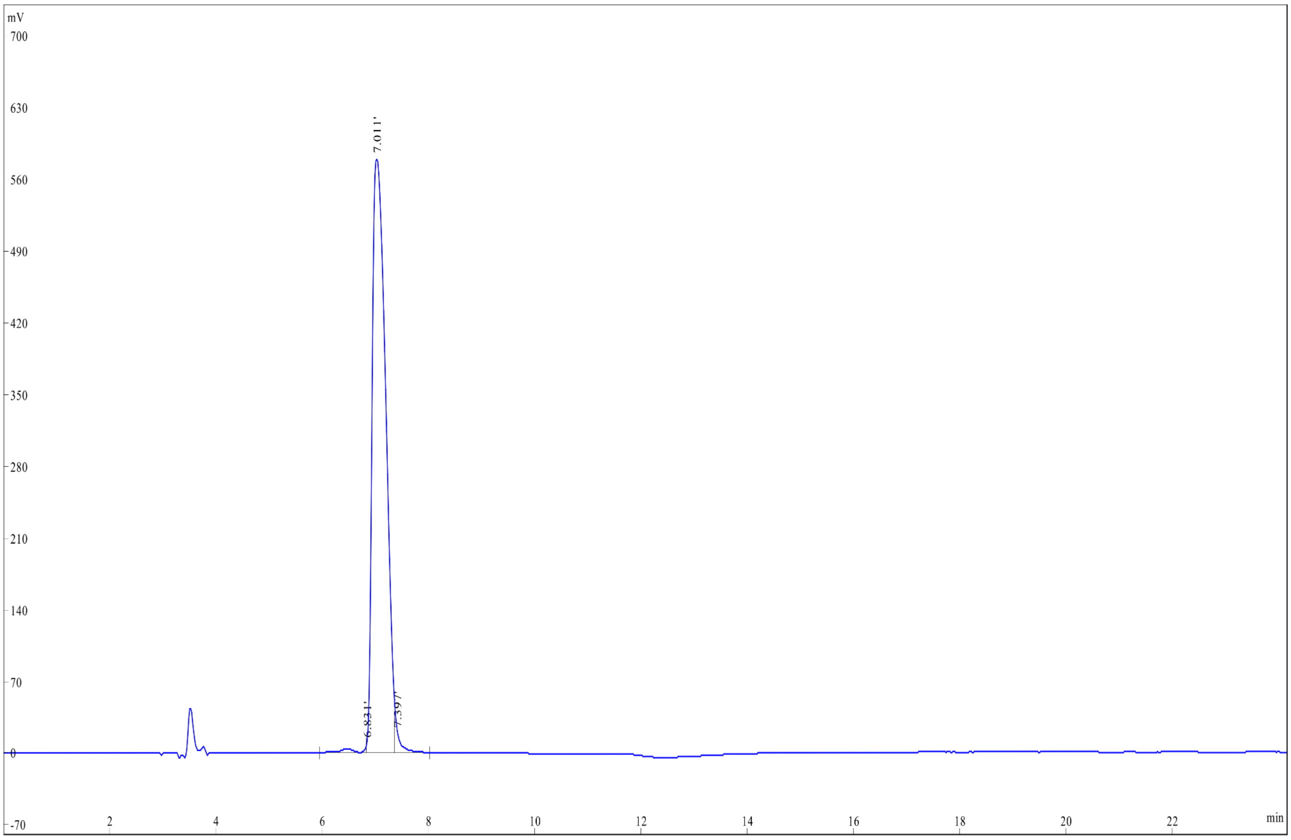


| Rank | Time | Conc. | Area | Height |
|---|---|---|---|---|
| 1 | 6.831 | 0.9124 | 90329 | 8812 |
| 2 | 7.011 | 97.3574 | 9638234 | 578203 |
| 3 | 7.397 | 1.7302 | 171290 | 18894 |
| Total | | 100 | 9899853 | 605909 |

### Analysis Report

#### <Sample Information>

Sample Name : WWP3
Sample ID : fr19-23
Data Filename : fr19-23.lcd
Method Filename : 90 to0 A_14min_1mL min_235nm.lcm
Batch Filename :
Vial # : -1
Sample Type : Unknown
Injection Volume : 100 uL
Date Acquired : 09/03/2022 13:57:04
Acquired by : System Administrator
Date Processed : 09/03/2022 14:18:05
Processed by : System Administrator

#### <Chromatogram>

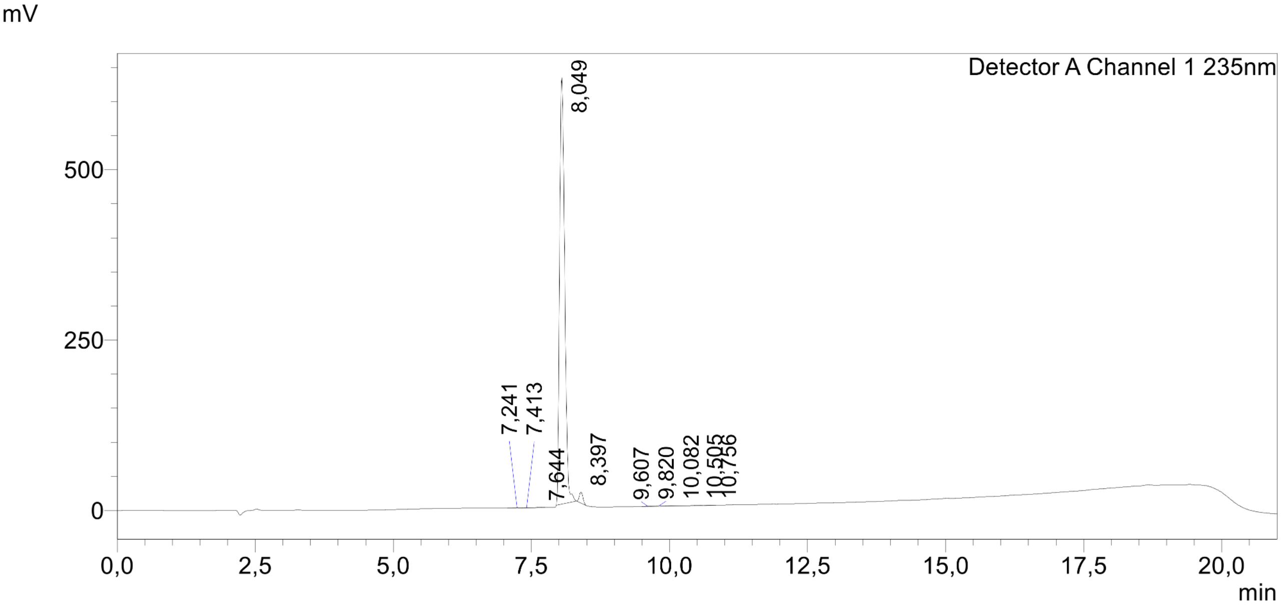


#### <Peak Table>

Detector A Channel 1 235nm

| Peak# | Ret. Time | Area | Height | Conc. | Unit | Mark | Name |
|---|---|---|---|---|---|---|---|
| 1 | 7,241 | 5574 | 680 | 0,135 | | | |
| 2 | 7,413 | 1341 | 205 | 0,032 | | V | |
| 3 | 7,644 | 4690 | 810 | 0,113 | | V | |
| 4 | 8,049 | 4036055 | 625631 | 97,515 | | M | |
| 5 | 8,397 | 73893 | 16047 | 1,785 | | | |
| 6 | 9,607 | 2651 | 420 | 0,064 | | | |
| 7 | 9,820 | 7275 | 871 | 0,176 | | V | |
| 8 | 10,082 | 1870 | 185 | 0,045 | | V | |
| 9 | 10,505 | 3587 | 565 | 0,087 | | V | |
| 10 | 10,756 | 1986 | 305 | 0,048 | | | |
| Total | | 4138923 | 645718 | | | | |

Sample Name :ENFA1-A
Sample ID :U918B027G0-29
Time Processed :15:19:42
Month-Day-Year Processed :04/20/2025

Pump A : 0.065% trifluoroacetic in 100% water (v/v)
Pump B : 0.05% trifluoroacetic in 100% acetonitrile (v/v)
Total Flow:1 ml/min
Wavelength:220 nm
<<LC Time Program>>

| Time | Module | Command | Value |
|---|---|---|---|
| 0.01 | Pumps | B.Conc | 5 |
| 25.00 | Pumps | B.Conc | 65 |
| 25.01 | Pumps | B.Conc | 95 |
| 27.00 | Pumps | B.Conc | 95 |
| 27.01 | Pumps | B.Conc | 5 |
| 35.00 | Pumps | B.Conc | 5 |
| 35.01 | Controller | Stop | |

<<Column Performance>>
<Detector A>
Column :Inertsil ODS-SP 4.6 x 250 mm
Equipment: ZJ17010509

#### <Chromatogram>

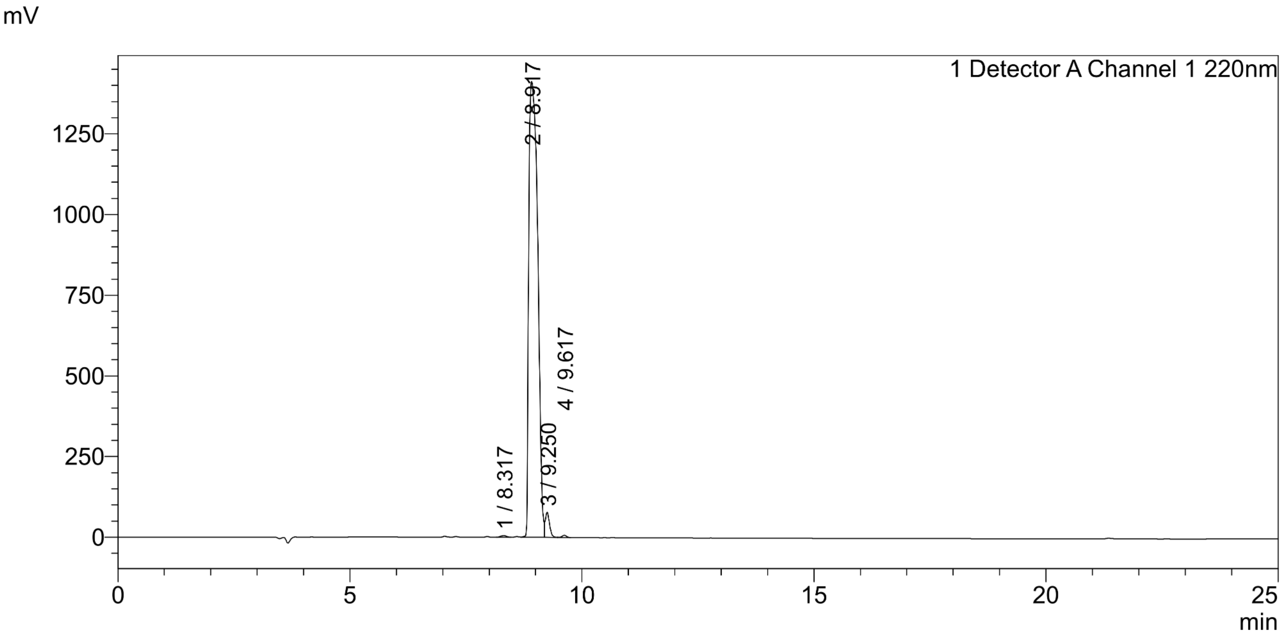


#### <Peak Table>

Detector A Channel 1 220nm

| Peak# | Ret. Time | Area | Height | Area% |
|---|---|---|---|---|
| 1 | 8.317 | 43404 | 5316 | 0.224 |
| 2 | 8.917 | 18827704 | 1412834 | 96.976 |
| 3 | 9.250 | 499054 | 77494 | 2.570 |
| 4 | 9.617 | 44554 | 6743 | 0.229 |
| Total | | 19414716 | 1502387 | 100.000 |

Sample Name :ENFA1-B
Sample ID :U918B027G0-33
Time Processed :10:36:26
Month-Day-Year Processed :04/30/2025

Pump A : 0.065% trifluoroacetic in 100% water (v/v)
Pump B : 0.05% trifluoroacetic in 100% acetonitrile (v/v)
Total Flow:1 ml/min
Wavelength:220 nm
<<LC Time Program>>

| Time | Module | Command | Value |
|---|---|---|---|
| 0.01 | Pumps | B.Conc | 5 |
| 25.00 | Pumps | B.Conc | 65 |
| 25.01 | Pumps | B.Conc | 95 |
| 27.00 | Pumps | B.Conc | 95 |
| 27.01 | Pumps | B.Conc | 5 |
| 35.00 | Pumps | B.Conc | 5 |
| 35.01 | Controller | Stop | |

<<Column Performance>>
<Detector A>
Column :Inertsil ODS-SP 4.6 x 250 mm
Equipment: ZJ21010376

#### <Chromatogram>

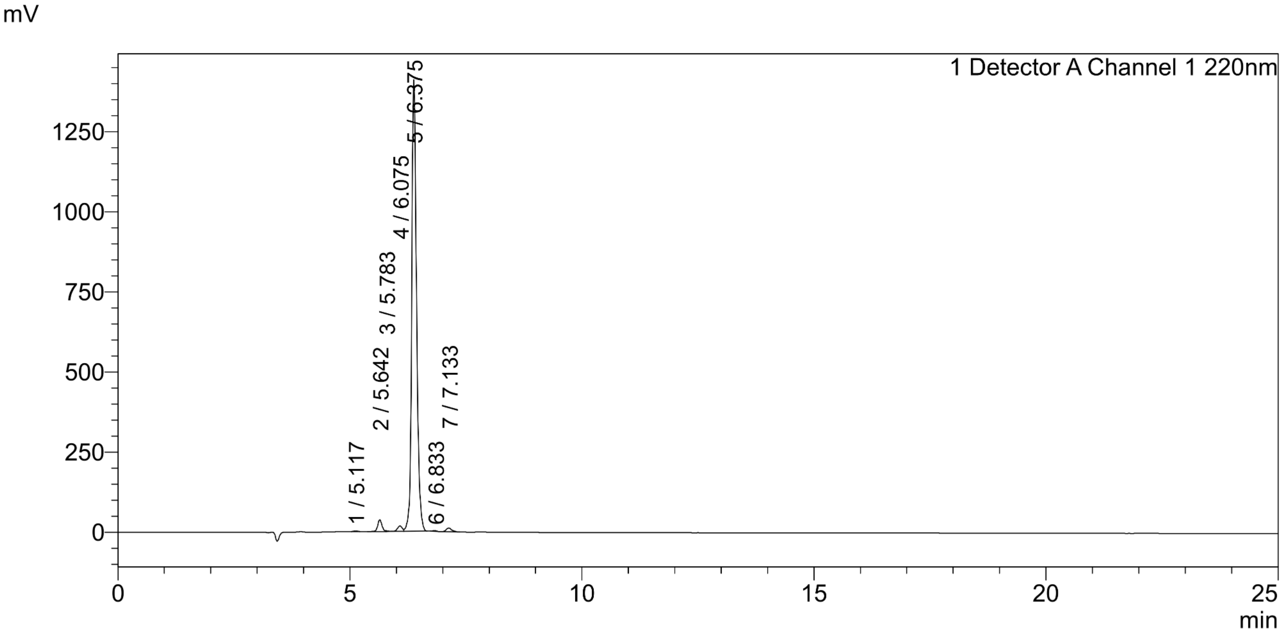


#### <Peak Table>

Detector A Channel 1 220nm

| Peak# | Ret. Time | Area | Height | Area% |
|---|---|---|---|---|
| 1 | 5.117 | 12830 | 2181 | 0.121 |
| 2 | 5.642 | 241374 | 37308 | 2.280 |
| 3 | 5.783 | 7087 | 2174 | 0.067 |
| 4 | 6.075 | 119452 | 16764 | 1.128 |
| 5 | 6.375 | 10089085 | 1409169 | 95.309 |
| 6 | 6.833 | 12057 | 2490 | 0.114 |
| 7 | 7.133 | 103737 | 12038 | 0.980 |
| Total | | 10585622 | 1482124 | 100.000 |

Sample Name: **CF-[WI23-B]**

**<Chromatogram>**

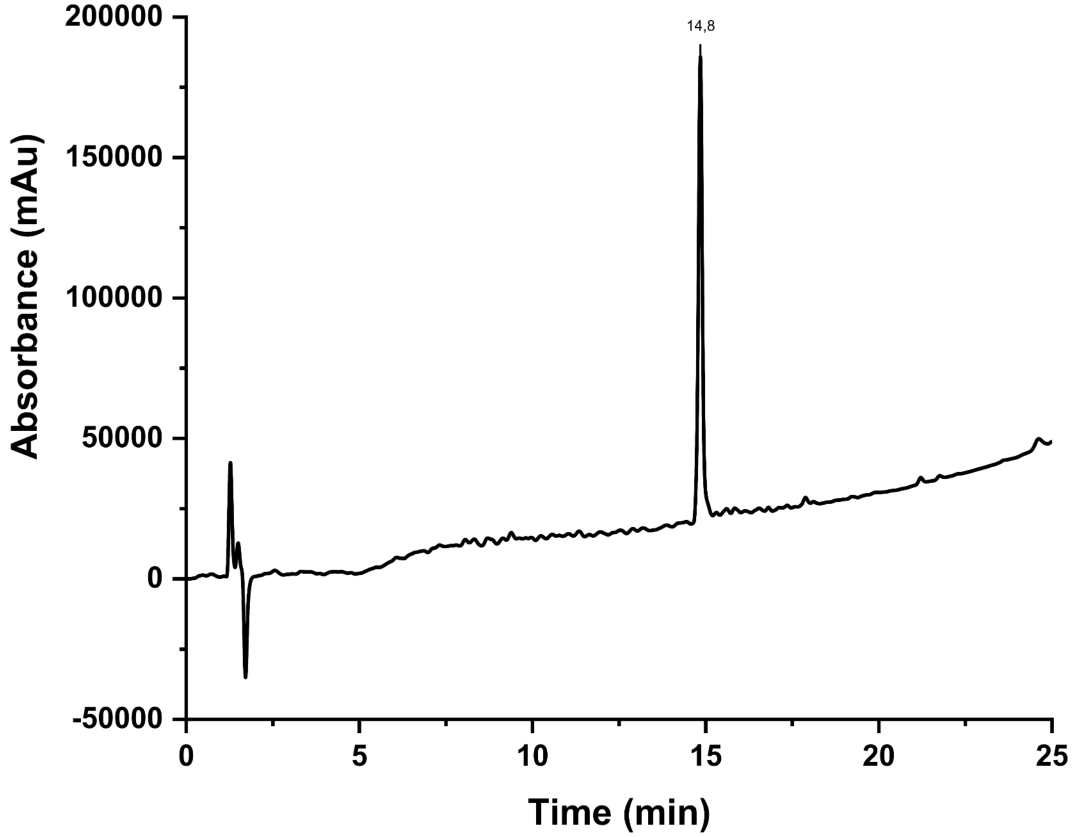

#### MS spectra

Correspondence between the names of the peptides during synthesis with those reported in the manuscript.

| Manufacturer | Synthesis Name | Manuscript Name |
|---|---|---|
| Proteogenix | New_pep2 TE-4 | **WI3** |
| Proteogenix | New_pep1 ST-4 | **WI6** |
| Proteogenix | Enfa6 AC-WY-4 | **WI13** |
| Proteogenix | Enfa7 AC-QY-4 | **WI14** |
| Proteogenix | Enfa1 AC-RQ-4 | **WI23** |
| Proteogenix | Enfa2 AC-YQ-4 | **WI25** |
| Proteogenix | Enfa3 AC-FC-4 | **WI26** |
| Proteogenix | Enfa4 AC-YH-4 | **WI29** |
| *in-house* | WWP3 | **WI6-B** |
| Genscript | ENFA1-A | **WI23-B** |
| Genscript | ENFA1-B | **WI23-E** |
| *in-house* | CF-[WI23-B] | **CF-[WI23-B]** |

### MASS SPECTROMETRY REPORT

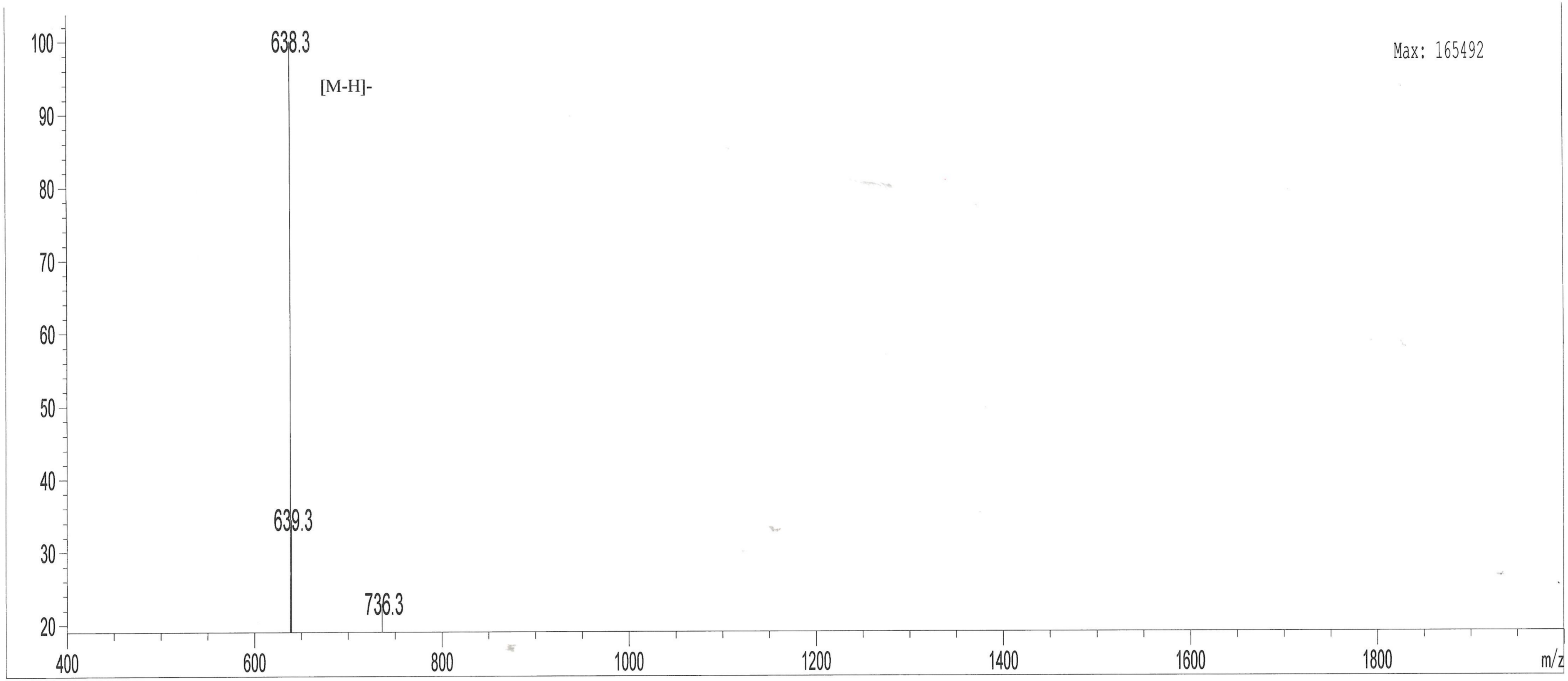


| | | | | | |
|---|---|---|---|---|---|
| Sample Information | | Probe: | ESI | Probe Bias: | +4.5kv |
| Injection Volume | : 1.00 μl | Nebulizer Gas Flow: | 1.5L/min | Detector: | 1.5kv |
| Sample: | New_pep2 TE-4 | CDL: | -20.0v | T. Flow: | 0.2ml/min |
| M.W.: | 639.65 | CDL Temp.: | 250 °C | B. Conc.: | 50%H2O/50%ACN |
| Lot. No.: | P201104-HS843563 | Block Temp.: | 200 °C | | |

### MASS SPECTROMETRY REPORT

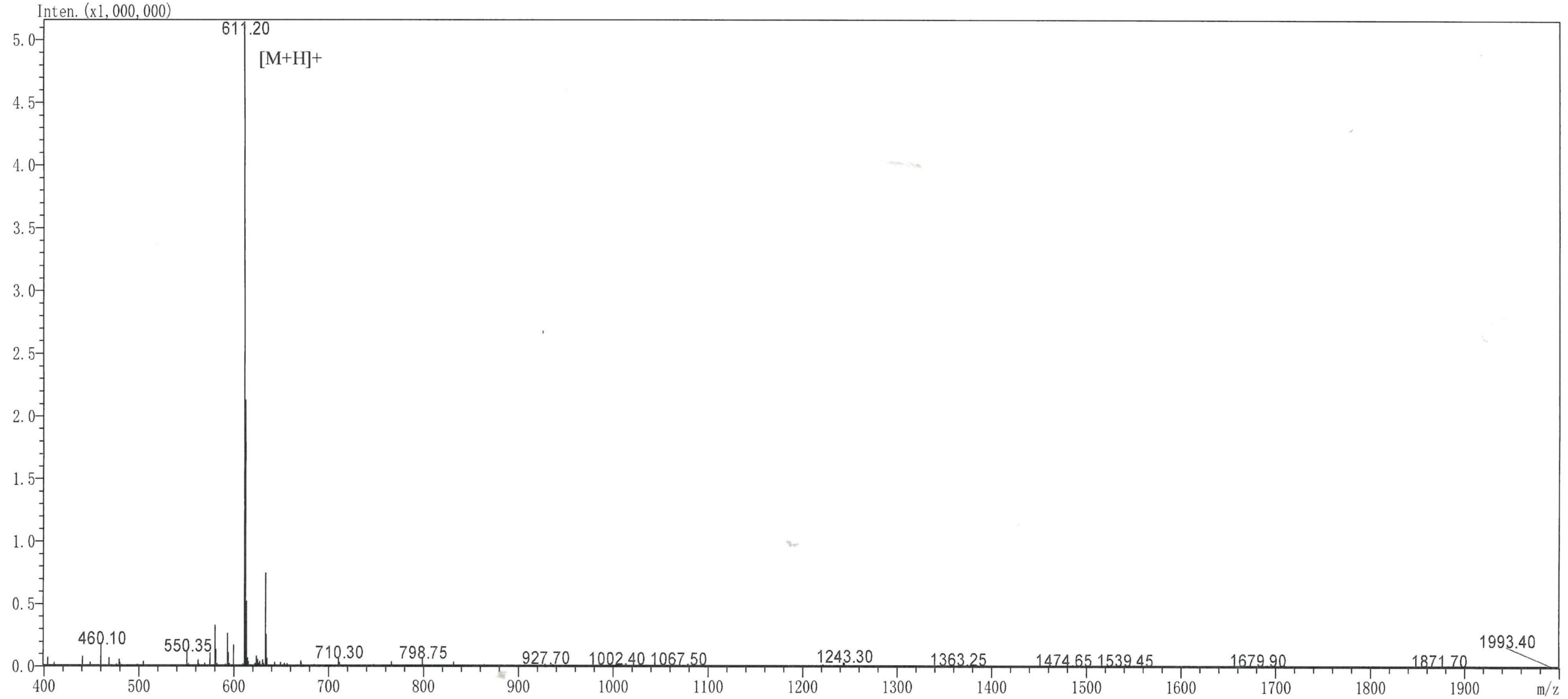


| | | | | | |
|---|---|---|---|---|---|
| Sample Information | | Probe: | ESI | Probe Bias: | +4.5kv |
| Injection Volume | : 1.00 µ l | Nebulizer Gas Flow: | 1.5L/min | Detector: | 1.2kv |
| Sample: | ST-4 | CDL: | -20.0v | T. Flow: | 0.2ml/min |
| M.W.: | 610.66 | CDL Temp.: | 250 °C | B. Conc.: | 50%H2O/50%ACN |
| Lot. No.: | P210518-CQ898185 | Block Temp.: | 400 °C | | |

### MASS SPECTROMETRY REPORT

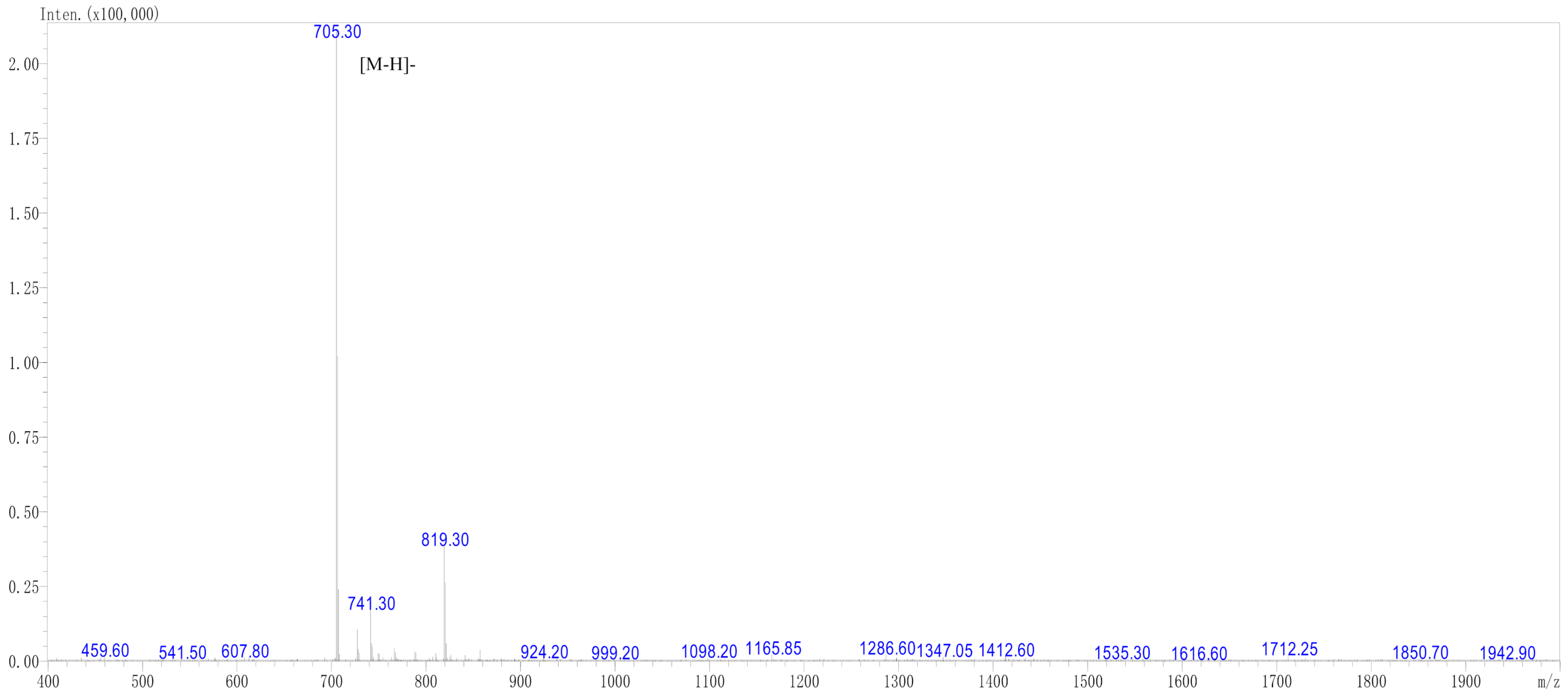


| Sample Information | | Probe: | ESI | Probe Bias: | +4.5kv |
|---|---|---|---|---|---|
| Injection Volume : | 1.00 μ l | Nebulizer Gas Flow: | 1.5L/min | Detector: | 1.2kv |
| Sample: | Ac-WY-4 | CDL: | -20.0v | T. Flow: | 0.2ml/min |
| M.W.: | 706.79 | CDL Temp.: | 250 °C | B. Conc.: | 50%H2O/50%ACN |
| Lot. No.: | P210305-CQ874402 | Block Temp.: | 400 °C | | |

### MASS SPECTROMETRY REPORT

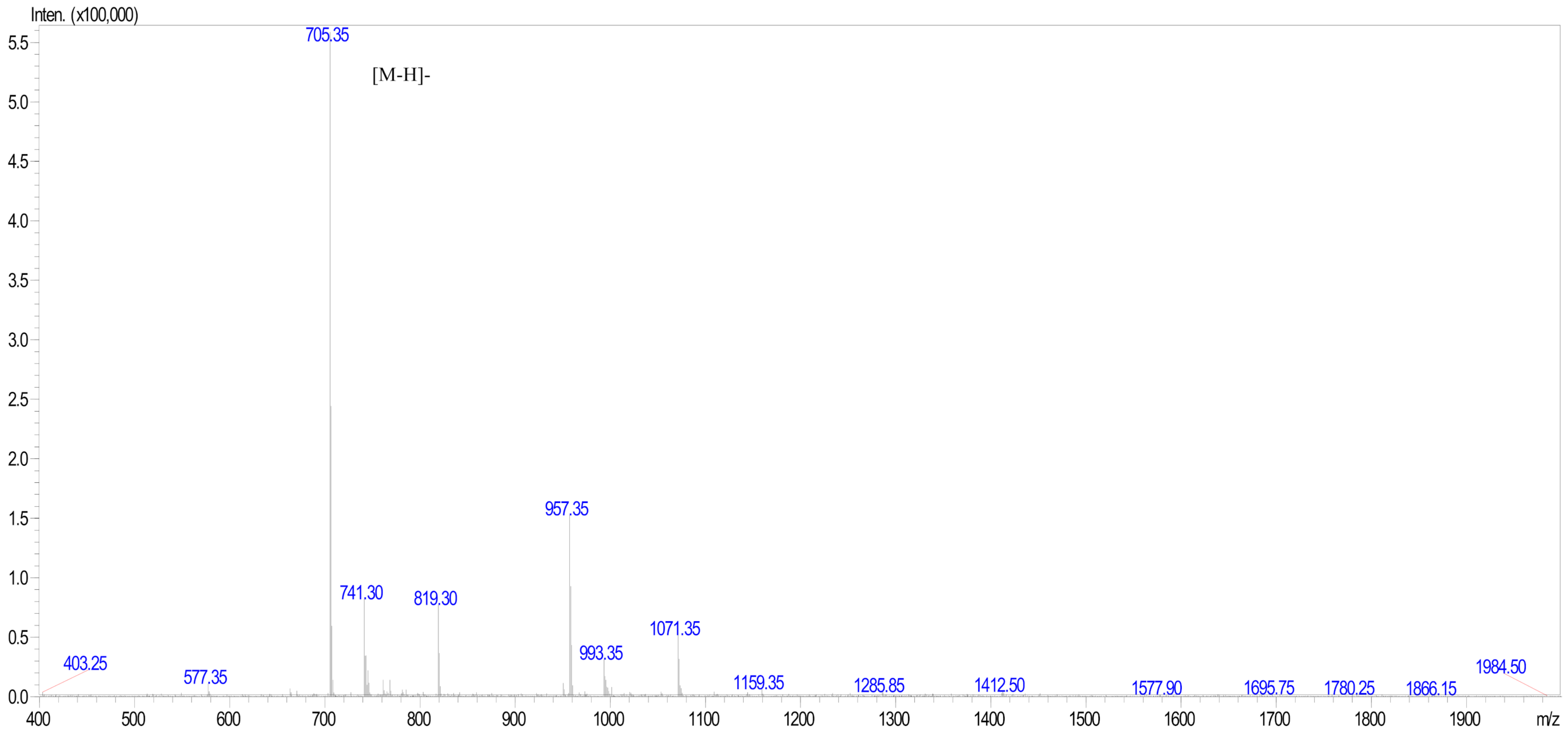


| | | | | | |
|---|---|---|---|---|---|
| Sample Information | | Probe: | ESI | Probe Bias: | ＋4.5kv |
| Injection Volume | : 1.00 μ l | Nebulizer Gas Flow: | 1.5L/min | Detector: | 1.2kv |
| Sample: | Enfa7 AC-QY-4 | CDL: | -20.0v | T. Flow: | 0.2ml/min |
| M.W.: | 706.79 | CDL Temp.: | 250 °C | B. Conc.: | 50%H2O/50%ACN |
| Lot. No.: | P210305-CQ874403 | Block Temp.: | 400 °C | | |

### MASS SPECTROMETRY REPORT

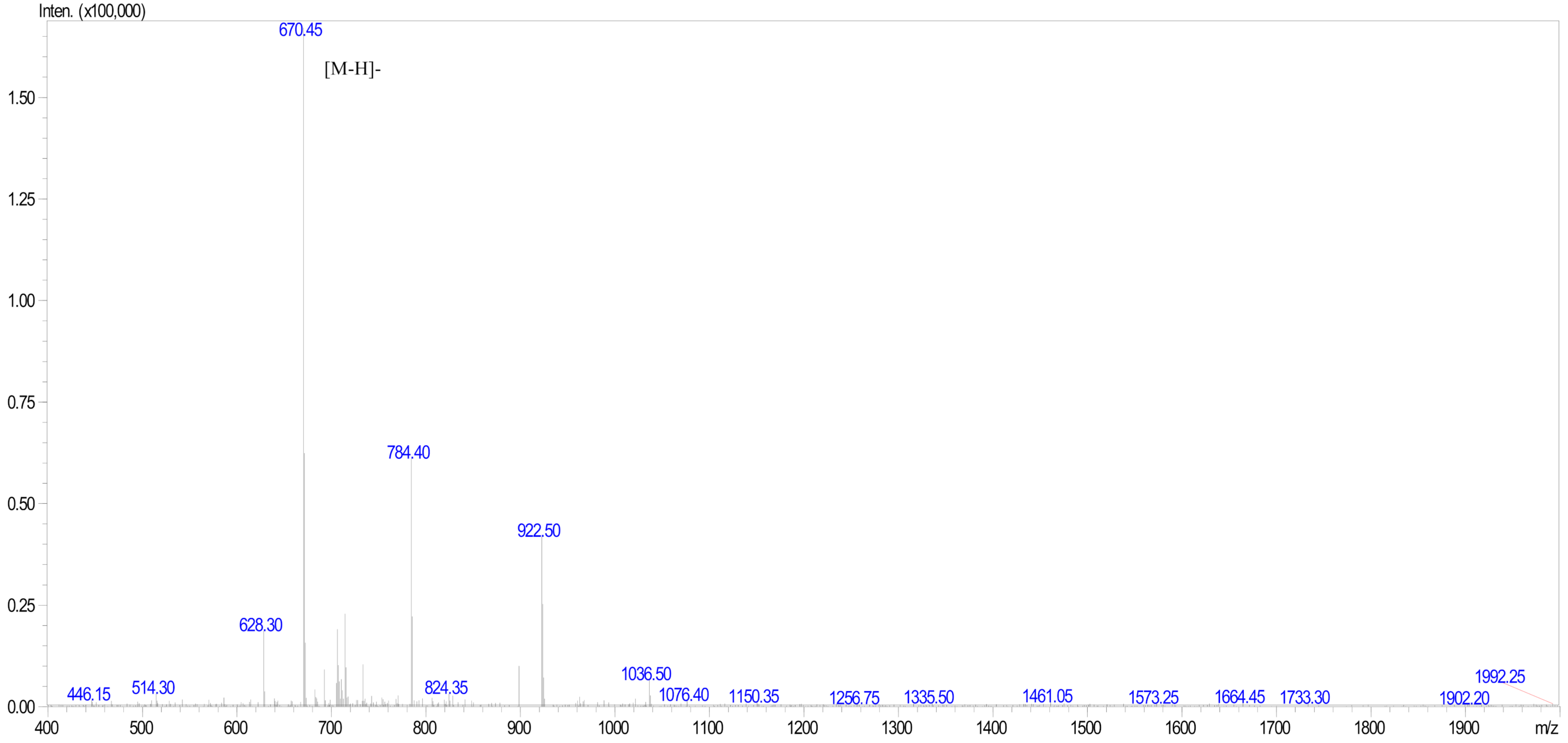


| | | | | | |
|---|---|---|---|---|---|
| Sample Information | | Probe: | ESI | Probe Bias: | +4.5kv |
| Injection Volume | : 1.00 µl | Nebulizer Gas Flow: | 1.5L/min | Detector: | 1.2kv |
| Sample: | enfa1 AC-RQ-4 | CDL: | -20.0v | T. Flow: | 0.2ml/min |
| M.W.: | 671.79 | CDL Temp.: | 250 °C | B. Conc.: | 50%H2O/50%ACN |
| Lot. No.: | P210305-CQ874397 | Block Temp.: | 400 °C | | |

### MASS SPECTROMETRY REPORT

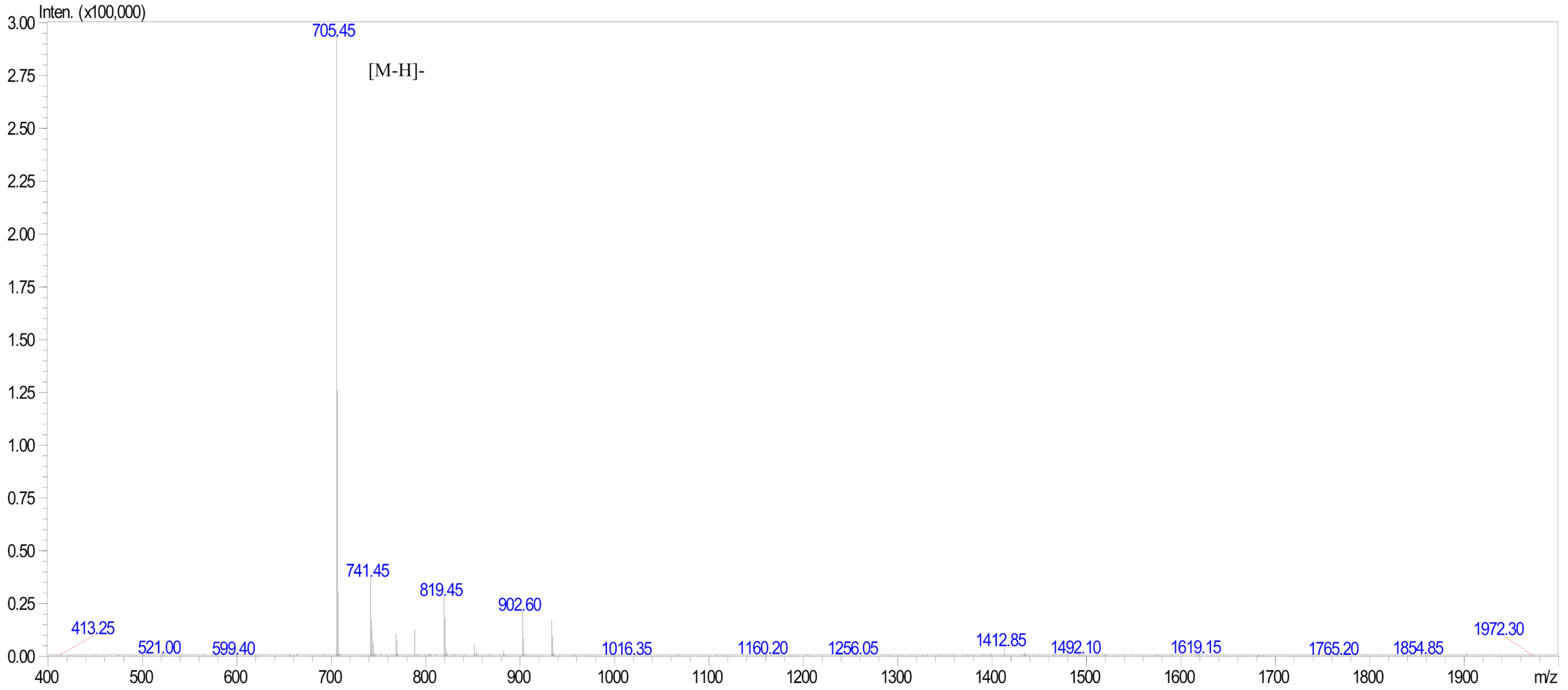


Sample Information
Injection Volume : 1.00 µl
Sample: ENFA2 AC-YQ-4
M.W.: 706.79
Lot. No.: P210305-CQ874398

Probe: ESI
Nebulizer Gas Flow: 1.5L/min
CDL: -20.0v
CDL Temp.: 250 °C
Block Temp.: 400 °C

Probe Bias: +4.5kv
Detector: 1.2kv
T. Flow: 0.2ml/min
B. Conc.: 50%H2O/50%ACN

### MASS SPECTROMETRY REPORT

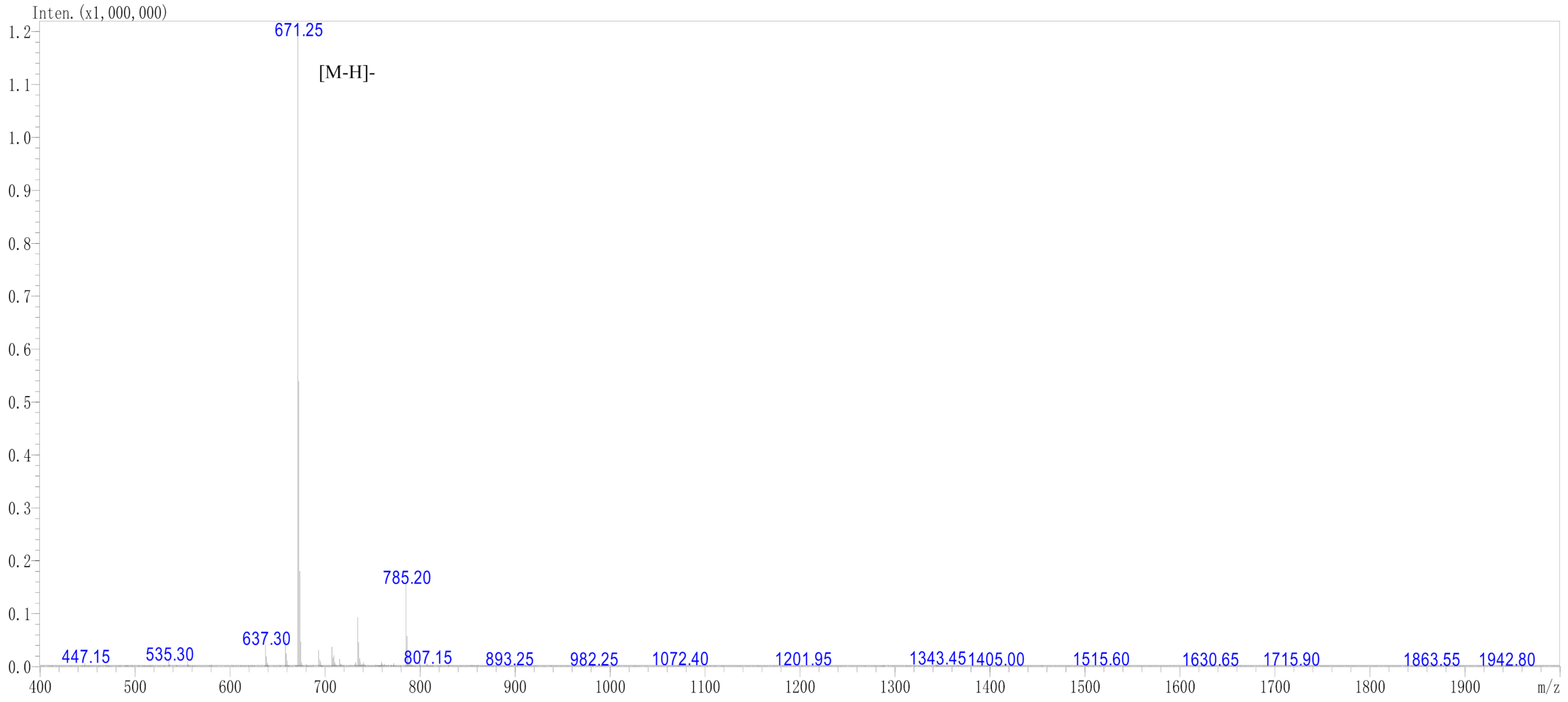


| | | | | | |
|---|---|---|---|---|---|
| Sample Information | | Probe: | ESI | Probe Bias: | +4.5kv |
| Injection Volume : | 1.00 µl | Nebulizer Gas Flow: | 1.5L/min | Detector: | 1.2kv |
| Sample: | Ac-FC-4 | CDL: | -20.0v | T. Flow: | 0.2ml/min |
| M.W.: | 672.79 | CDL Temp.: | 250 °C | B. Conc.: | 50%H2O/50%ACN |
| Lot. No.: | P210305-CQ874399 | Block Temp.: | 400 °C | | |

### MASS SPECTROMETRY REPORT

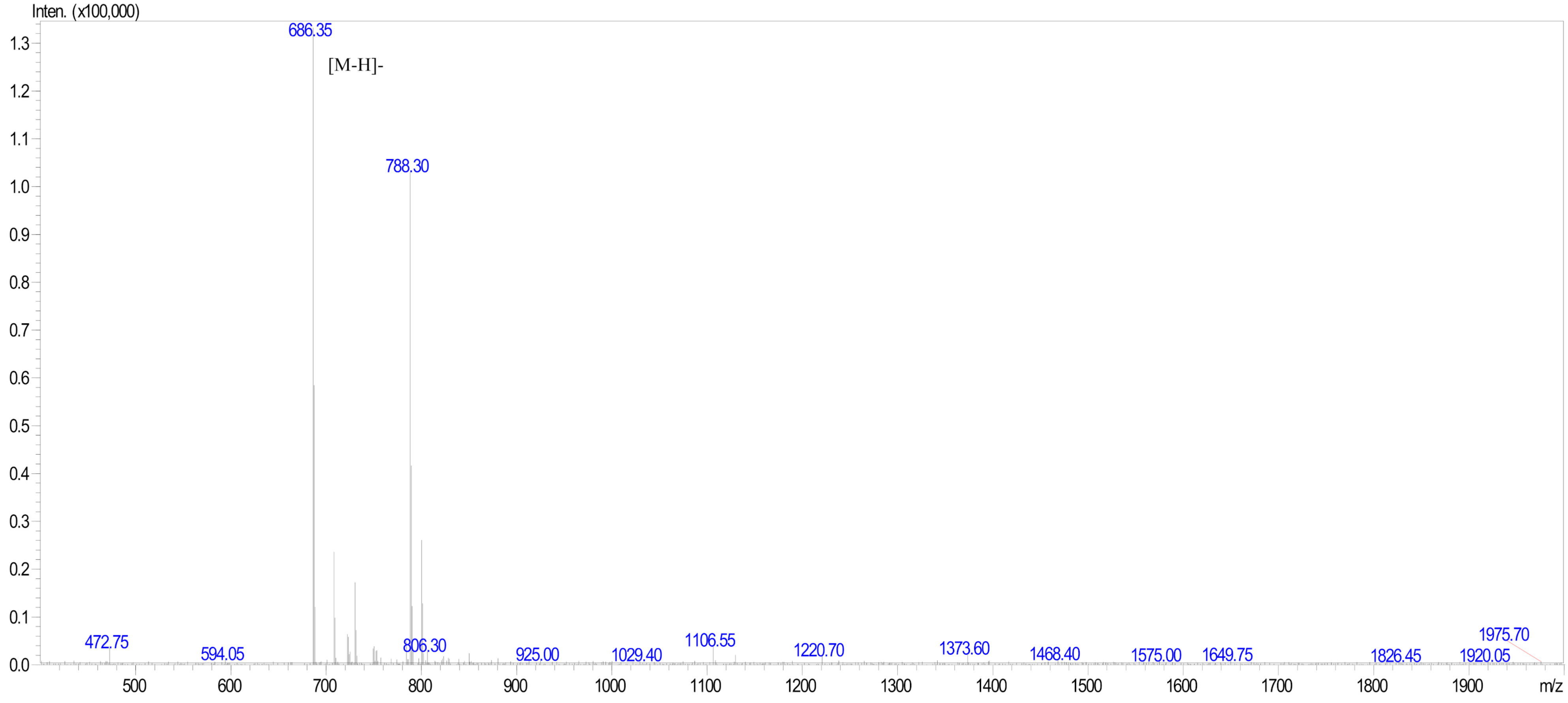


| | | | | | |
|---|---|---|---|---|---|
| Sample Information | | Probe: | ESI | Probe Bias: | +4.5kv |
| Injection Volume : | 1.00 μ l | Nebulizer Gas Flow: | 1.5L/min | Detector: | 1.2kv |
| Sample: | Enfa4 Ac-YH-4 | CDL: | -20.0v | T. Flow: | 0.2ml/min |
| M.W.: | 687.79 | CDL Temp.: | 250 °C | B. Conc.: | 50%H2O/50%ACN |
| Lot. No.: | P210305-CQ874400 | Block Temp.: | 400 °C | | |

Laboratory of Biomolecular Chemistry

Istituto di Scienze e Tecnologie Chimiche "Giulio Natta"

via M. Bianco 9, 20131 Milano (IT)
www.scitec.cnr.it

Sample Information

| | |
|---|---|
| Acquired by | : System Administrator |
| Date Acquired | : 14/03/2022 11:55:53 |
| Sample ID | : WWP3_1a500 |
| Data File | : WWP3_1a500.lcd |
| Method File | : 95 to 0_0.5mL_min.lcm |
| Date Processed | : 14/03/2022 12:25:59 |

MS Spectrum
WWP3_1a500

1

ESI Scan +Positive
Averaged

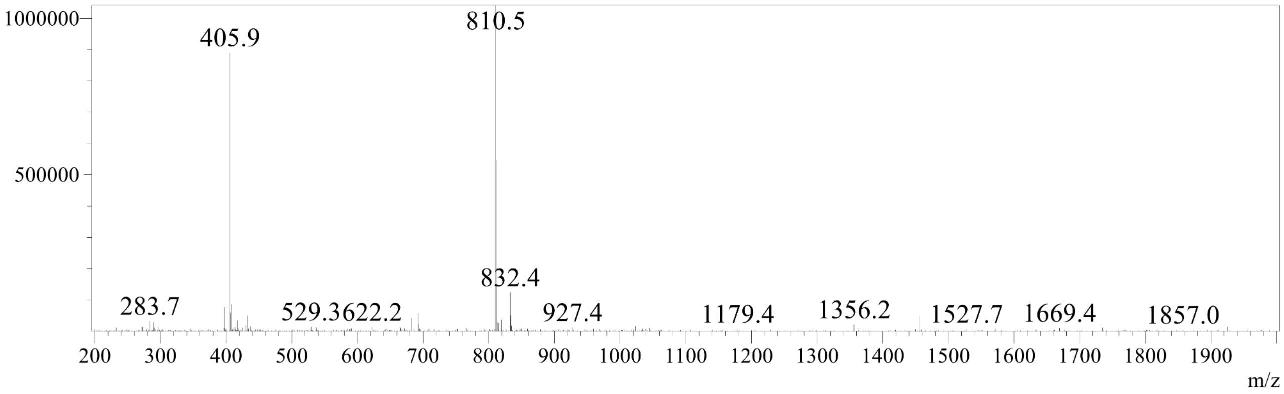

#### Mass Spectrum

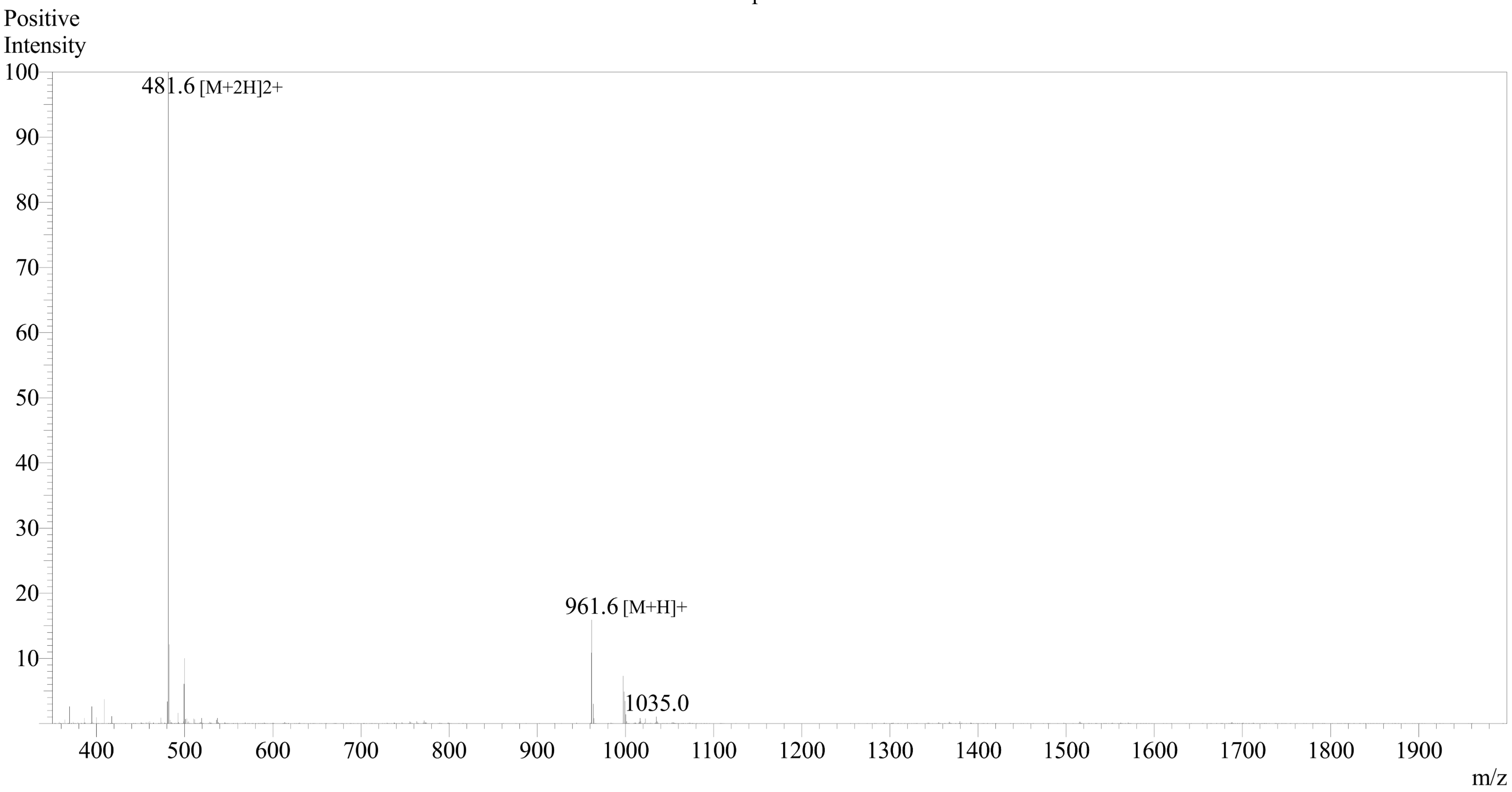


Sample Information
Month-Day Processed : 04/20/25
Time Processed : 16:34:10
Injection Volume : 0.2
Sample Name : ENFA1-A
Sample ID : U918B027G0-29
Theoretical MW : 961.14
Observed MW : 961.2

Interface :ESI
Nebulizing Gas Flow :1.5L/min
CDL Temp :250
Block Temp :200

Equipment : ZJ21010035
Interface Bias : +4.5 kV
Drying Gas Flow :5 L/min
T.Flow :0.2 ml/min
B.conc :50%H2O/50%MeOH

Mass Spectrum

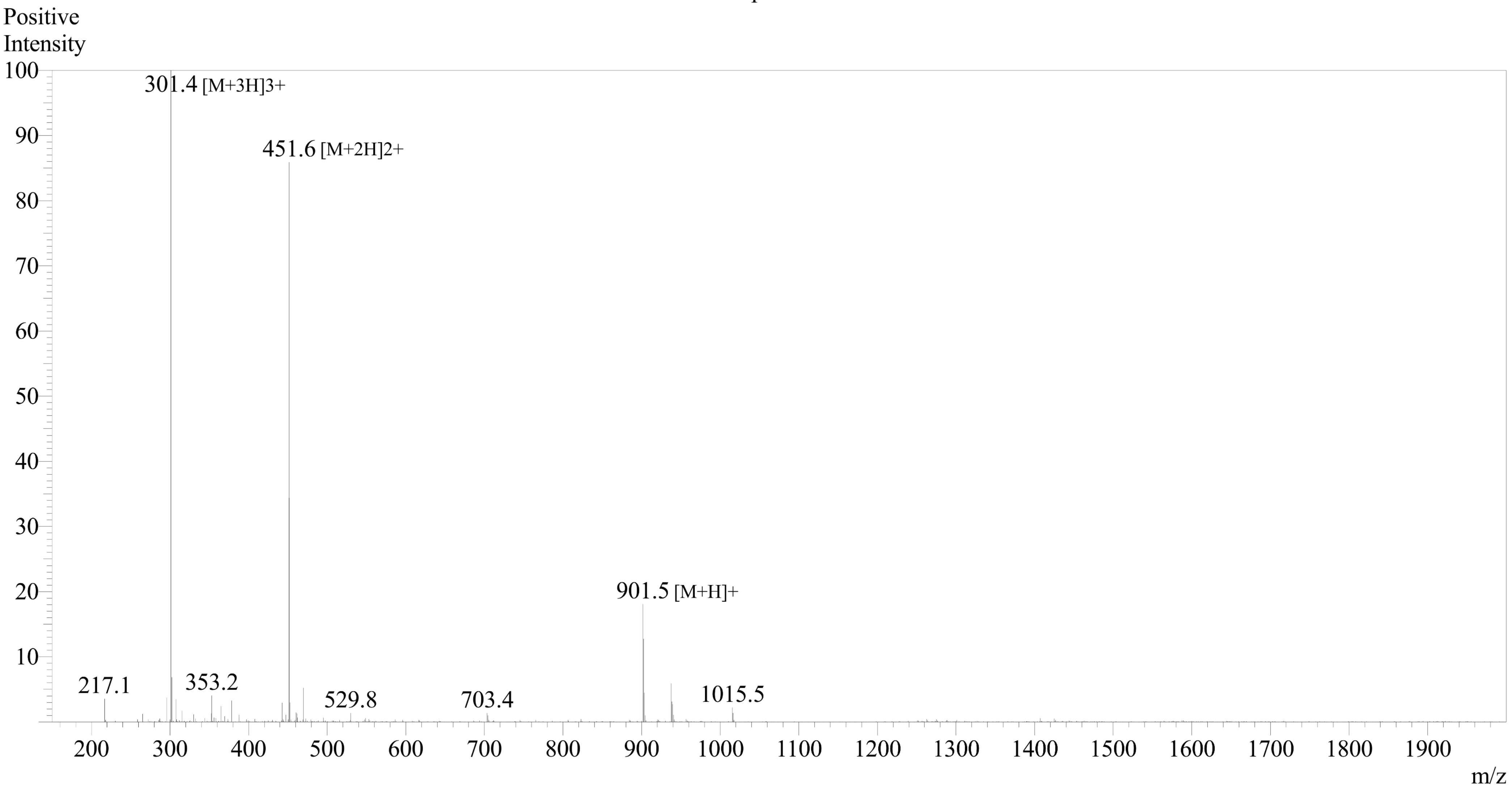


Sample Information
Month-Day Processed : 04/29/25
Time Processed : 20:22:59
Injection Volume : 0.2
Sample Name : ENFA1-B
Sample ID : U918B027G0-33
Theoretical MW : 901.04
Observed MW : 901.2

Interface :ESI
Nebulizing Gas Flow :1.5L/min
CDL Temp :250
Block Temp :200

Equipment :ZJ22010150
Interface Bias : +4.5 kV
Drying Gas Flow :5 L/min
T.Flow :0.2 ml/min
B.conc :50%H2O/50%MeOH

Sample Name: **CF-[WI23-B]**

**<ESI-MS spectra>**

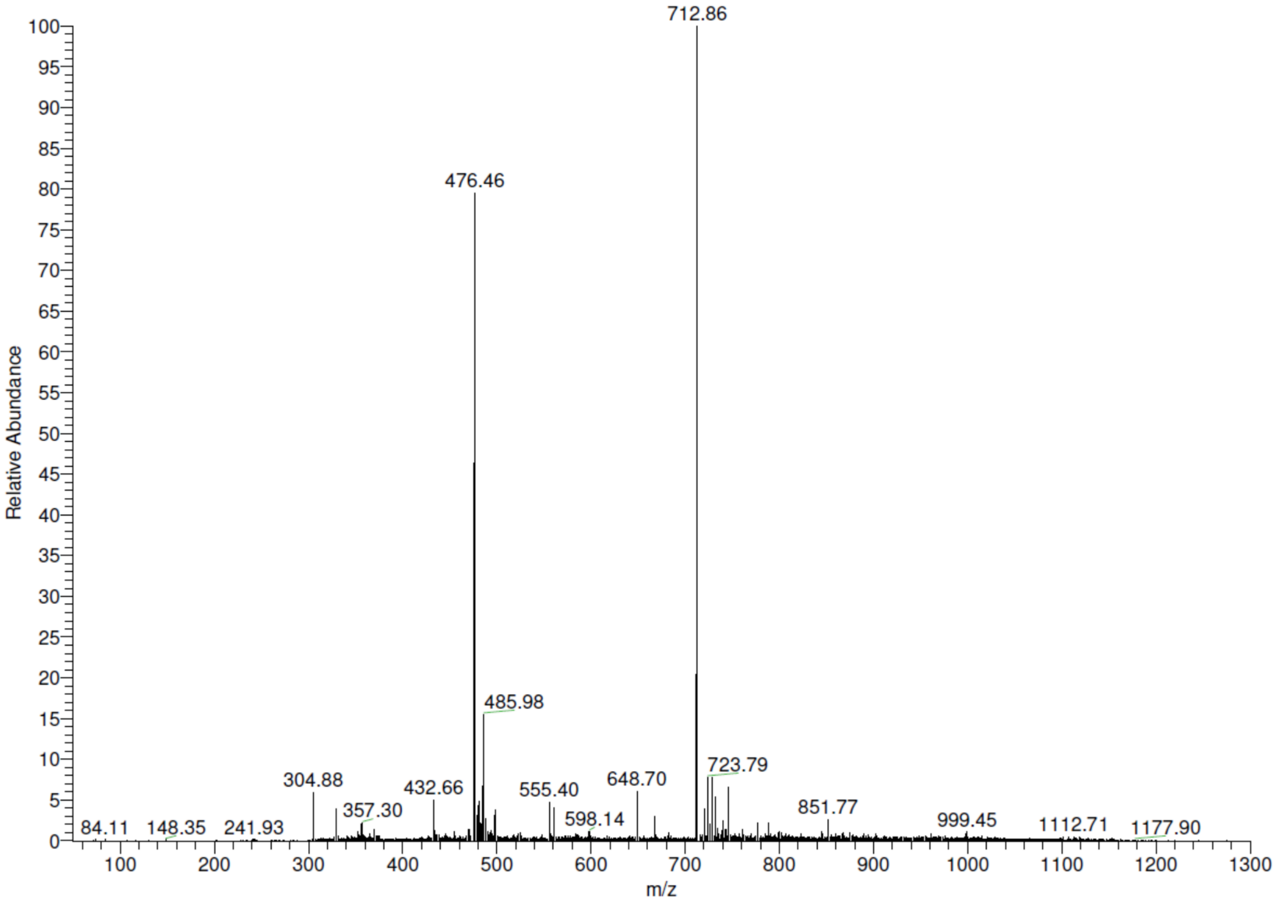


The peaks correspond to: $[M+2H]^{2+}$ (712.86 m/z) and $[M+3H]^{3+}$ (476.46 m/z).
